# Peax Interactive Visual Pattern Search in Sequential Data Using Unsupervised Deep Representation Learning

**DOI:** 10.1101/597518

**Authors:** Fritz Lekschas, Brant Peterson, Daniel Haehn, Eric Ma, Nils Gehlenborg, Hanspeter Pfister

## Abstract

We present Peax, a novel feature-based technique for interactive visual pattern search in sequential data, like time series or data mapped to a genome sequence. Visually searching for patterns by similarity is often challenging because of the large search space, the visual complexity of patterns, and the user’s perception of similarity. For example, in genomics, researchers try to link patterns in multivariate sequential data to cellular or pathogenic processes, but a lack of ground truth and high variance makes automatic pattern detection unreliable. We have developed a convolutional autoencoder for unsupervised representation learning of regions in sequential data that can capture more visual details of complex patterns compared to existing similarity measures. Using this learned representation as features of the sequential data, our accompanying visual query system enables interactive feedback-driven adjustments of the pattern search to adapt to the users’ perceived similarity. Using an active learning sampling strategy, Peax collects user-generated binary relevance feedback. This feedback is used to train a model for binary classification, to ultimately find other regions that exhibit patterns similar to the search target. We demonstrate Peax’s features through a case study in genomics and report on a user study with eight domain experts to assess the usability and usefulness of Peax. Moreover, we evaluate the effectiveness of the learned feature representation for visual similarity search in two additional user studies. We find that our models retrieve significantly more similar patterns than other commonly used techniques.

## 1 Introduction

Visually searching for patterns in sequential data can be challenging when the search space is large, the data is complex, or the search query is difficult to formalize. Visual query systems simplify the query formalization by allowing analysts to find regions in the data given a visual description of the pattern, often in the form of a sketch [32,50,67] or an example [7]. Using the query, the system retrieves regions that are most similar given some notion of similarity. But the search can fail when the analyst’s subjectively perceived similarity does not match the system’s similarity measure [17]. The larger the query region, the more likely it is that the query contains several distinct visual features, such as peaks, trends, or troughs, which can be hard to capture with current techniques (Figure 2). Not knowing what visual feature is important to the user makes similarity search even more challenging.

While well-known distance metrics, like Euclidean distance (ED) and dynamic time warping (DTW), in combination with one-nearest neighbor search are traditionally used for classification of sequential data [16], Correll and Gleicher have found that “no single algorithm accounts for human judgments of time series similarity” [12]. They instead suggest to combine several metrics and let the analyst choose which one to use for pattern search. To this end, the distance measures act as a feature representation of the sequential data. Such an approach has been used before [24,25,34], as a first step to address the issue. However, it assumes that the handcrafted distance metrics are able to capture many important pattern variations [12], that the analyst is aware of the visual features in the query pattern that they care about, and that the analyst knows which individual distance metric can identify or ignore the variations of interest. If the latter two do not hold, the visual query system can become inefficient and confuse the analysts rather than leading to successful exploration—known as the gulf of execution and evaluation [56]. Prior work on natural image search [21] has shown that user-provided binary relevance feedback can be a powerful approach to interactively learn which visual aspects are important without introducing a complicated user interface.

To address these challenges, we propose a novel feature-based approach for assessing the similarity of visual patterns in sequential data using convolutional autoencoder models. An autoencoder is a type of neural network that learns an identity function of some data. By introducing a lower-dimensional layer during the encoding step, the neural network is forced to find a compressed latent representation of the input data that captures as many visual details of the input data as possible (Figure 1.1). We use this learned latent representation as features to compare the similarity between patterns. Since the subjectively perceived similarity of an analyst is not known upfront we have developed Peax, a visual query system that interactively learns a classifier for pattern search through binary relevance feedback (Figure 1.3). After the analyst defines a query by example, Peax samples potentially interesting regions based on their distance in the learned latent space from the autoencoder models. We employ a sliding window approach with a user-defined window size and resolution to limit the search space. The analyst has to label the sampled windows as either interesting or not interesting to provide training data for a random forest classifier. After the classifier is trained, we employ an active learning strategy (Figure 1.2) to focus the labeling process on the windows that are close to the query window, located in dense areas in the latent space, and hard to predict by the classifier. Peax supports exploration of multivariate sequential data by concatenating multiple latent representations provided by potentially different autoencoders. This enables the analyst to adjust the combination data to be explored on the fly without having to re-train the autoencoder.

**Figure 1:**
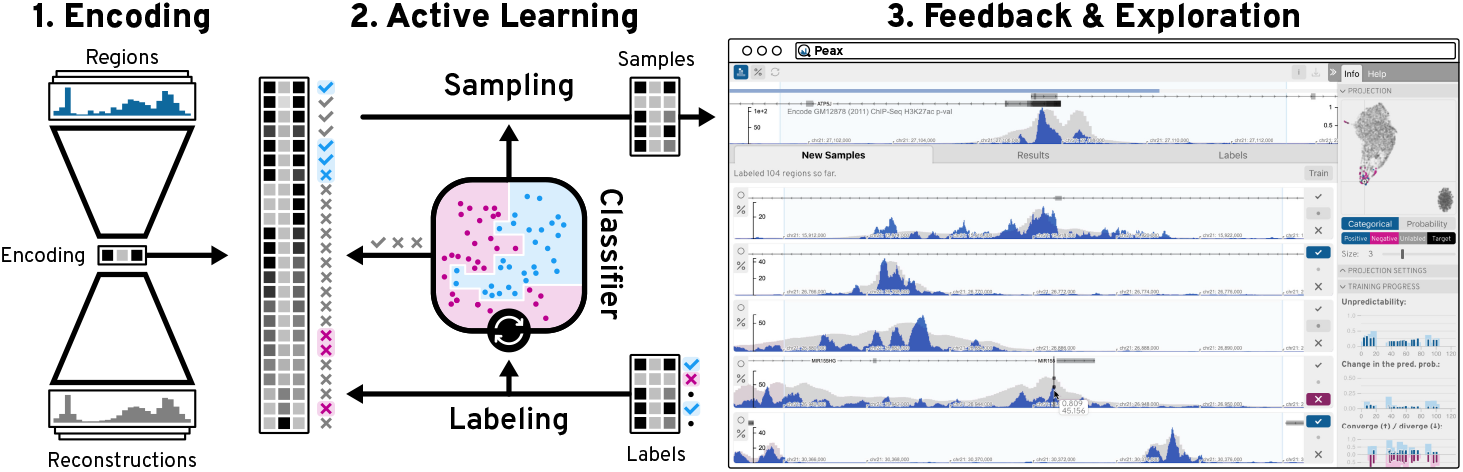
The Peax System. Using an autoencoder model (1), regions of sequential data are encoded into a compressed latent representation. Using this encoding, Peax employs an active learning strategy (2) to focus the labeling (3) on regions that are useful for training a classifier. This classifier is then iteratively trained on the user’s binary labels (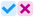) to predict the interestingness of other regions in the data.

We apply our technique on two epigenomic datasets and demonstrate how Peax can be used to explore biological phenomena (section 8). In epigenomics, biomedical researchers study how the genome of an organism is regulated through various modifications of the DNA and associated proteins that do not change the underlying DNA sequence. A better understanding of epigenomic modifications is crucial as, for example, genome-wide association studies have found that over 90% of all disease-associated DNA sequence variants are located in non-coding regions [39, 52, 63] that are most likely acting on gene regulation. Finding epigenomic patterns is challenging [23, 59] as the inter- and intra-dataset variance is high, the data is typically very noisy, and ground truth is missing in almost all cases. Furthermore, formally describing biological phenomena is often challenging due to the complexity of datasets or different interpretations of the same biological mechanism [22].

To evaluate our approach, we first assess the autoencoder’s ability to reconstruct visual features in sequential patterns and measure how the learned latent representation can be used for similarity search in two Mechanical Turk user studies. Compared to six commonly used techniques for similarity search, we find that our model retrieves patterns that are perceived significantly more similar by the participants than any other technique. Additionally, we evaluated the usability and usefulness of Peax through an in-person user study with eight domain experts in epigenomics. The results show that Peax is easy to learn and use, and that it offers a new and effective way of exploring epigenomic patterns. To the best of our knowledge, we present the first deeplearning-based approach for interactive visual pattern search in sequential data. The source code for Peax, instructions on how our autoencoder models are trained, and a set of 6 autoencoders [42] for two types of epigenomic data are available online at https://github.com/Novartis/peax/.

*Supplementary Figure and Table references are prefixed with an “S”*.

## 2 Related Work

**Similarity Measures** for pattern search in sequential data have been studied extensively [72]. Techniques for similarity search include distance-based and feature-based methods. Distance-based methods provide a single value for how different patterns are compared to each other. The two most widely used distance measures with consistently strong performance [16, 66] are Euclidean distance [19] (ED) and dynamic time warping [5] (DTW). Feature-based methods use a set of features describing the data in conjunction with a distance-based method or machine learning technique for comparison and classification [9,12,25,34–36,44]. Widely-used feature-based methods include, for example, piecewise aggregate approximation [35] (PAA) and symbolic aggregate approximation [44] (SAX), which are part of a class called symbolic representations. Both methods discretize sequential data into segments of equal size and aggregate these segments into a new representation. We apply PAA (Figure 4.1c) to downsample the segmented data as a preprocessing step. Other feature-based approaches combine several distance metrics [12, 34] into a feature representation assuming that a combination will capture more variations. Fulcher and Jones [24, 25] took this approach to an extreme and proposed a supervised feature extraction technique that represents a time series by a combination of over 9000 analysis algorithms. Christ et al. [10] extended this approach but we find that it does not provide comparable performance in an unsupervised setting (subsection 9.2). General purpose dimensionality reduction techniques such as t-SNE [49] or UMAP [53] can also be used to learn a lower-dimensional embedding as a feature representation. But while they are useful for visualization purposes, we show that their embedding is not effective for visual pattern search (Figure 8).

Finally, autoencoder [4] (AE) models provide a data-driven approach to learn a feature presentation. It was shown that AEs can extract a hierarchy of features from natural images [51], textual data like electronic health records [54], or scatter plot visualizations [48]. AEs have also been applied on time series data for classification [20] and predictions [27,47,60]. Furthermore, AEs are also used in genomics for general feature learning [68] and prediction [71] of gene expression data. In this paper, we extend upon these works and leverage AEs to learn feature representations for interactive similarity search in sequential data.

**Visual Query Systems** (VQS) can be divided into two general approaches that let the user query for patterns by example or by sketching. Time-Searcher [29, 30] is an early instance of a query-by-example system that supports value-based filtering and similarity search [7] using rectangular boxes drawn on top of a time series visualization. QueryLines [62] is a sketch-based filtering technique for defining strict or fuzzy value limits based on multiple straight lines. Time series that conflict with a limit are either filtered or de-emphasized visually. QuerySketch [67] is one of the first systems that supports similarity search by sketching. The user can directly draw onto the visualized time series to find similar instances using the ED. Holz and Feiner [32] combined the ideas of QuerySketch and QueryLines and developed a tolerance-aware sketch-based system. Their technique measures shape and time deviation during sketching to determine which parts of the sketched query should be relaxed. Recently, Mannino and Abouzied presented Qetch [50] for querying time series data based on scale-free hand-drawn sketches. Qetch ignores global scaling differences and instead determines the best hit according to local scale and shape distortions. However, Lee et al. [40] find that sketch-based systems are rarely used in real-world situations as users are often unable to articulate precisely what they are looking for and, therefore, have a hard time sketching the correct query. Peax does not use a sketch-based strategy as it is hard to to accurately draw complex patterns as shown in Figure 2.

**Figure 2:**
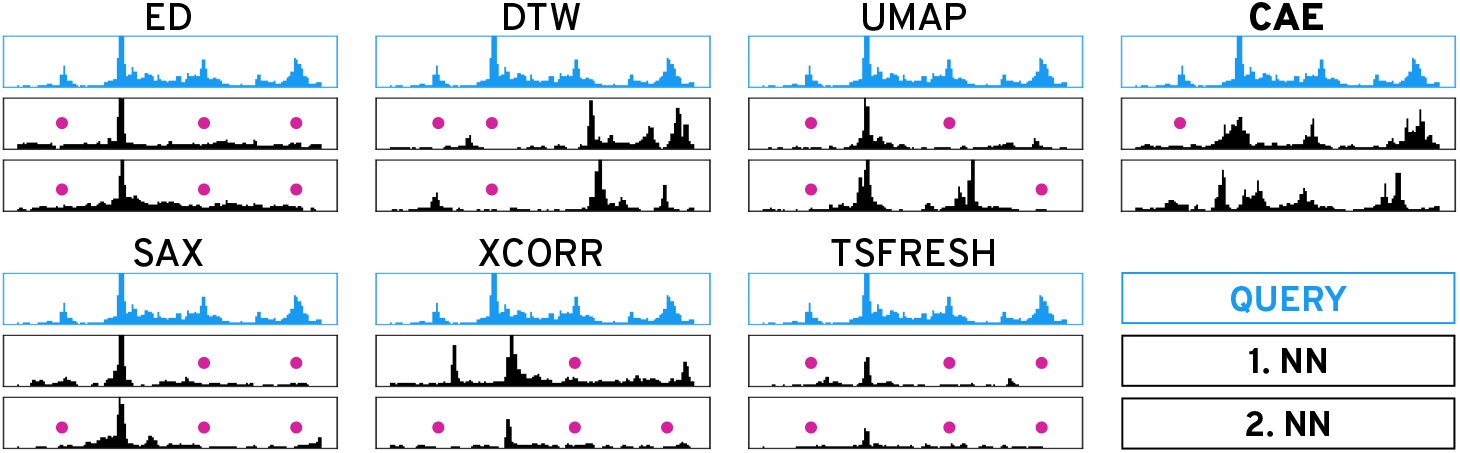
Similarity Search. Given the query (blue), the two nearest neighbors of the six methods on the left fail to capture many salient peaks (pink dots). While our autoencoder-based technique (CAE) also misses the small left-most peak it finds more similar instances.

Apart from the query interface, Keogh and Pazzani [36,37] show that the notion of similarity can heavily depend on the user’s mental model and might not be captured well with an objective similarity measure. Eichmann and Zgraggen [17] extend this line of work and show that the objective similarity can differ markedly from the perceived similarity, even for simple patterns. Corell and Gleicher [12] build upon these findings and define a set of ten *invariants*, i.e., visual variations or distortions that a user might want to ignore during pattern search. They evaluate three distance measures and find that “no single algorithm accounts for human judgments of time series similarity” [12]. This indicates that it might be impossible to develop a single distance metric that captures all perceived notions of similarity.

**Interactive Visual Machine Learning** is a powerful technique to adjust the results of a VQS through user interactions. For example, Keogh and Pazzani [36, 37] propose relevance-feedback driven query adjustments for time series. Given a query, their system asks the user to rank the three nearest neighbors. The ranking is then used to change the query using a weighted average of the original query and the ranked nearest neighbors. A similar approach has been proposed by Behrisch et al. [3] for scatter plots, where the user interactively trains a classifier based on the user’s feedback to learn to capture the interestingness of scatter plot views. Recently, Dennig et al. [15] proposed a system to interactively learn the best combination of feature descriptors and a distance function for pattern separability, assuming the existence and user’s knowledge about the feature descriptors. In contrast, CueFlik [21] is a tool for interactive concept learning in image search, which allows the user to rank the results of a text-based image search using simple binary feedback. CueFlik employs an active learning strategy to speed up the learning process by finding images that potentially help to separate interesting from non-interesting images. In our work we use an active learning sampling strategy for interactive binary labeling based on uncertainty sampling [64]. We extend this strategy with a distance-dependent term (section 5) to ensure that more similar regions are labeled before the exploration space is broadened.

## 3 Goals and Tasks

The primary goal of Peax is to find regions in sequential data that show an instance of the target pattern. The target pattern can either be defined manually by example or by a set of already existing labels derived elsewhere. In both cases, the user might either be interested in finding the *k* most similar matches or in retrieving all windows that exhibit a pattern matching the target. The main difference between the two goals is whether recall is negligible (first case) or essential (second case). To achieve either goal, the user needs to know which windows are predicted to match the target pattern. Therefore, it is crucial that the user understands what concept the classifier has learned. Since the classifier is interactively trained based on the user’s subjective labels, there is no objective metric for when to stop the training process. Thus, the user needs to be aware of the training progress to decide when to stop training.

As summarized in Figure 3, the user might either start with a query by example or a set of labeled data. To arrive at a set of matching windows the user has to provide labels in order to train the classifier and steer the pattern search. We identified four main avenues for training the classifier. First, users need to be able to explore unlabeled windows, ideally with a sampling strategy that chooses windows intelligently to reduce the labeling efforts. It also needs to be possible to manually select windows for targeted labeling. Once a classifier is trained, the user needs to be allowed to provide immediate feedback on the results to confirm or correct what the classifier assumes is interesting. Finally, examining and potentially re-labeling already-labeled windows is important to resolve conflicts between the user’s and predicted labels. We assume that the first two labeling strategies are more common when building a classifier from scratch. The latter two strategies seem more relevant when starting with already labeled data.

**Figure 3:**
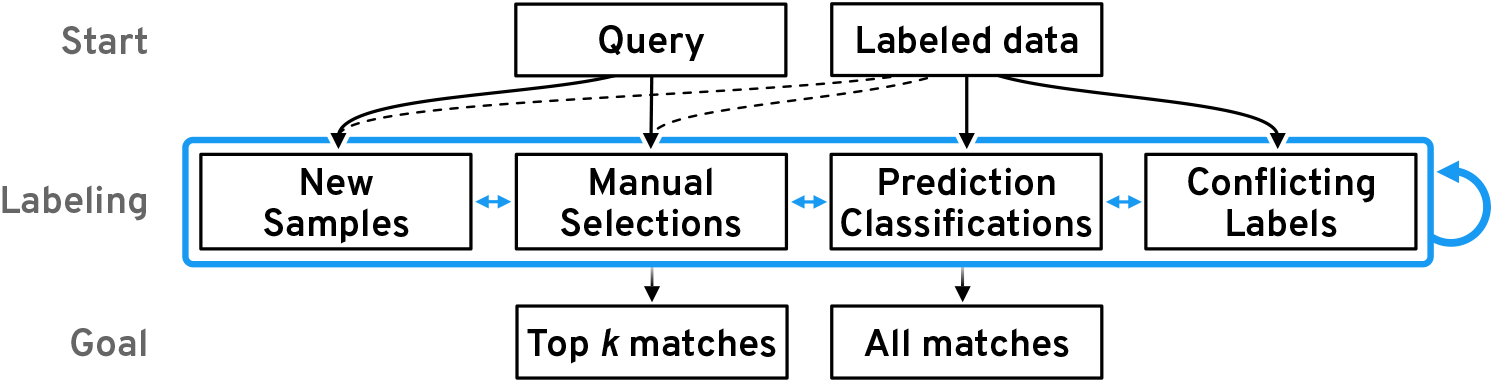
Workflow. A new search can be started with a query by example or already labeled data. The user then needs to provide and adjust labels to steer the classifier’s learning process.

Following these goals and this workflow (Figure 3), we have identified a set of high-level interaction tasks that underlie the design of Peax (section 6):

**T1 Freely browse and explore** to be able to gain an overview of the data and to find search queries.
**T2 Identify the classifier’s predictions** to be able to understand what the currently matching pattern is.
**T3 Compare windows** to identify similarities or dissimilarities among potentially related windows.
**T4 Contextualize windows** to understand the broader impact of a pattern, to improve confidence of the manually assigned labels, and to find related windows.
**T5 Visualize the learning progress** to highlight the impact of labeling and inform about the status of the trained classifier.
**T6 Show the capabilities of the latent representation** to realize if the visual features of the query pattern were captured.

## 4 Representation Learning

Instead of relying on handcrafted feature descriptors that do not always capture complex patterns well (Figure 8), we learn the feature representation in an unsupervised fashion using a convolutional autoencoder (CAE) model. To limit the search space and improve learnability, we employ a sliding window approach with a fixed window size and resolution as shown in Figure 4.1.

**Figure 4:**
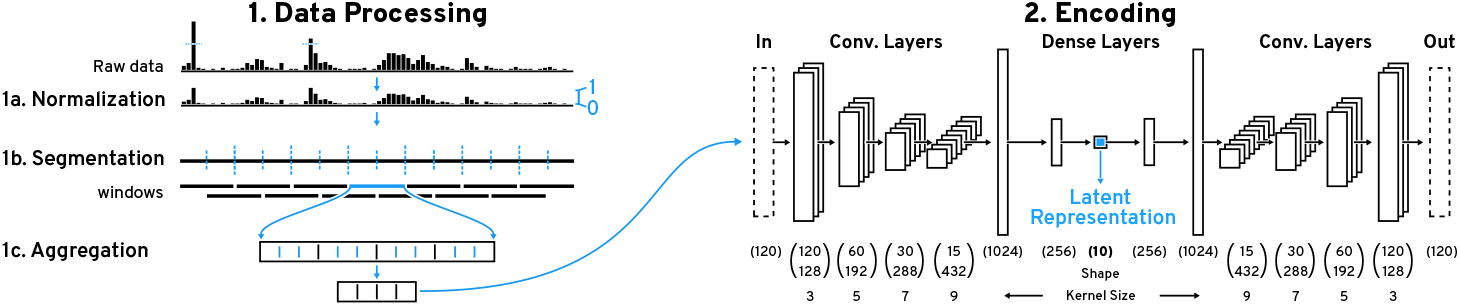
Data Processing and Encoding Pipeline. (1) We employ a sliding window approach for pattern search on (1a) piecewise (1b) aggregated (1c) windows. (2) The windows are encoded with our convolutional autoencoder (CAE) to obtain the latent representation. The CAE consists of a 4-layer convolutional encoder with an increasing number of filters and kernel size followed by two dense layers.

### 4.1 Data Preprocessing

We cap values at the 99.9th percentile to remove rare outliers and then scale the data to [0,1]. Then we split the data into overlapping windows w of fixed length *l*. The overlap is controlled by the step size *s* and step frequency *f*, where 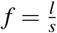. The step frequency *f* can be adjusted during the search process but the window length *l* is fixed. Therefore, *l* is critical and depends on the application (see section 8 for an in-depth example) as the CAE will only be able to recognize co-located patterns that appear within the window. Ideally one would want to use the highest possible *l* to capture more context but in practice the level of detail that the CAE can capture decreases with larger *l* (Figure 7). Thus, we suggest *l* to be slightly bigger than the largest pattern that is expected to be found. After segmentation, we downsample each window by a factor of *r* to allow the CAE to learn on data that looks similar to what a user would see when visualizing the data on a common computer screen. Also, downsampling has been shown to be an effective similarity search strategy on its own [35]. Finally, depending on the type of sequential data it might be necessary to filter out windows that contain very little to no visual features or that are highly overrepresented (section 8).

### 4.2 Convolutional Autoencoder

The preprocessed windows are used to train a CAE. The CAE consists of an encoder and a decoder model, where the encoder is trying to learn a transformation of the input data that the decoder is able to reconstruct into the original input as best as possible. Having the encoder output a vector that has fewer dimensions than the input forces the encoder to compress redundant information and to effectively learn features that are useful for reconstruction. Our encoder model (Figure 4.2) consists of four convolutional layers followed by two fully-connected layers. The four convolutional layers have 128, 192,

288, and 432 filters with kernel sizes of 3, 5, 7, and 9 respectively. Following Springenberg et al.’s approach [65] we use a striding of 2 instead of max pooling to shrink the convolved output per layer. Such architectures have previously been shown to enable hierarchical feature learning [51]. While the kernel size is typically unchanged in models for natural images, we found that larger kernel sizes lower the reconstruction error for sequential data (Supplementary Table S5). The two fully-connected layers consist of 1024 and 256 units. The final layer consists of 10 units and outputs the latent representation. All layers use the ReLu activation function. The CAE’s decoder model consists of the same layers just in reverse order and features an additional final layer with a sigmoid activation function to ensure that the output is within [0,1].

## 5 Active Learning

Peax employs two sampling strategies to select unlabeled windows that are subsequently shown to the analysts for labeling. Both strategies operate in the learned latent space of the CAE (subsection 4.2). The first sampling strategy is only used upon starting a new search when no classifier has been trained yet. Supplementary Figure S1 illustrates this initial sampling strategy. As classifier-related metrics such as uncertainty are unknown, the initial sampling strategy only relies on the average distance of a window w to its *k* nearest neighbors (default: 5) and the windows’ distance to the query in the latent space. Formally the average distance is defined as follows where *nn* is a function returning the ith nearest neighbor, and *norm* scales the input to [0,1].

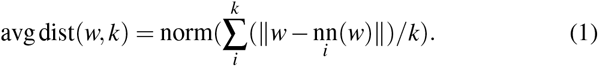

During the initial sampling strategy, we gradually increase the search radius such that positive and negative windows are sampled. The search radius r (default: 10) is defined as the number of r nearest neighbors that are taken into account in each initial sampling round. To sample from a relevant but also broad spectrum, we double the radius *r* in each of the *m* sampling rounds (default: 5). To not sample twice from the same neighborhood, we exclude windows from the previous search radii. In each round, we iteratively sample *n* windows (default: 5). We select the ones with lowest average distance to k nearest neighbors that maximize the distance from the already chosen samples S. Formally, we select the unlabeled window *u* from S, the complement of S, with the lowest following score:

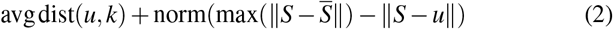

Sampling positive and negative windows is essential because we need both types of labels for the first training; otherwise, the initial classifier could be highly biased towards negative or positive windows. We include the average distance term to avoid sampling outliers, and we maximize the pairwise distance between sampled windows to let the user annotate diverse samples. After the first classifier has been trained, Peax switches to an active learning sampling strategy. This strategy extends the previous sampling approach with a distance-to-query and an uncertainty term. See Supplementary Figure S2 for a visual example. The distance-to-query term stands for the distance of a window *w* to the search query q. The uncertainty term, which is inspired by uncertainty sampling [64], is defined as

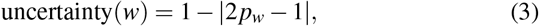

with the classifier’s prediction probability *p_w_* of a window w.

Finally, our active learning sampling strategy iteratively selects *n^a^* (default: 10) unlabeled windows. In each step, the unlabeled window *u* with the lowest following score is selected:

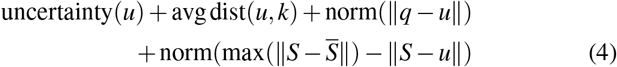

We sample uncertain windows that are within dense neighborhoods of the latent space to reduce the overall uncertainty of the classifier after the next training. The distance-to-query term is added to balance exploration and exploitation during the sampling strategy. Sampling highly uncertain regions broadens the exploration, especially at the beginning, when the classifier is generally more uncertain. On the other hand, sampling windows that are close to the query are expected to retrieve similar windows (subsection 9.2). Default values are derived from experience with our use cases (section 8).

### 5.1 Classifier

Given the small number of labels when training a new classifier from scratch and the repeatedly online training during the search, we choose to use a random forest classifier. Also, the random forest classifier shows superior performance compared to other popular binary classifiers (Supplementary Figure S49). This allows for efficient re-training each time the labels have changed or after requesting a new set of unlabeled windows based on the active learning sampling strategy. For searching across multiple sequential datasets, as shown in Figure 6, we concatenate the windows’ feature vectors from each datasets to avoid having to train a new CAE. Ultimately, we use the predicted classes to define the resulting set of matching windows.

**Figure 5:**
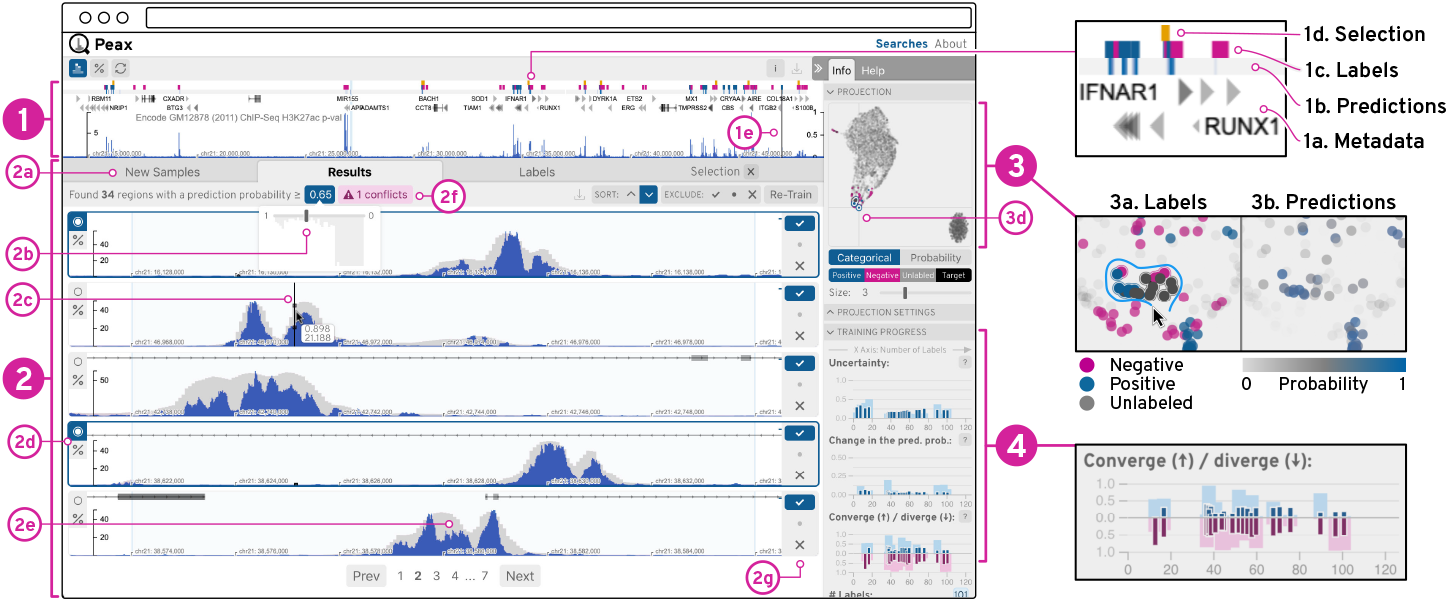
User Interface. The sequential data is visualized as blue bar charts (1&2), which can be superimposed with the CAE’s reconstruction as gray bars (2e). Additionally, the query view (1) can show metadata (1a-d) as region-based annotations. The list view (2) shows windows for comparison and labeling. The embedding view (3) represents windows as dots on a 2D canvas. The progress view (4) visualizes training metrics of the classifiers as composed bar charts (4), where the dark and light bars are relative to the labeled and all windows respectively.

**Figure 6:**
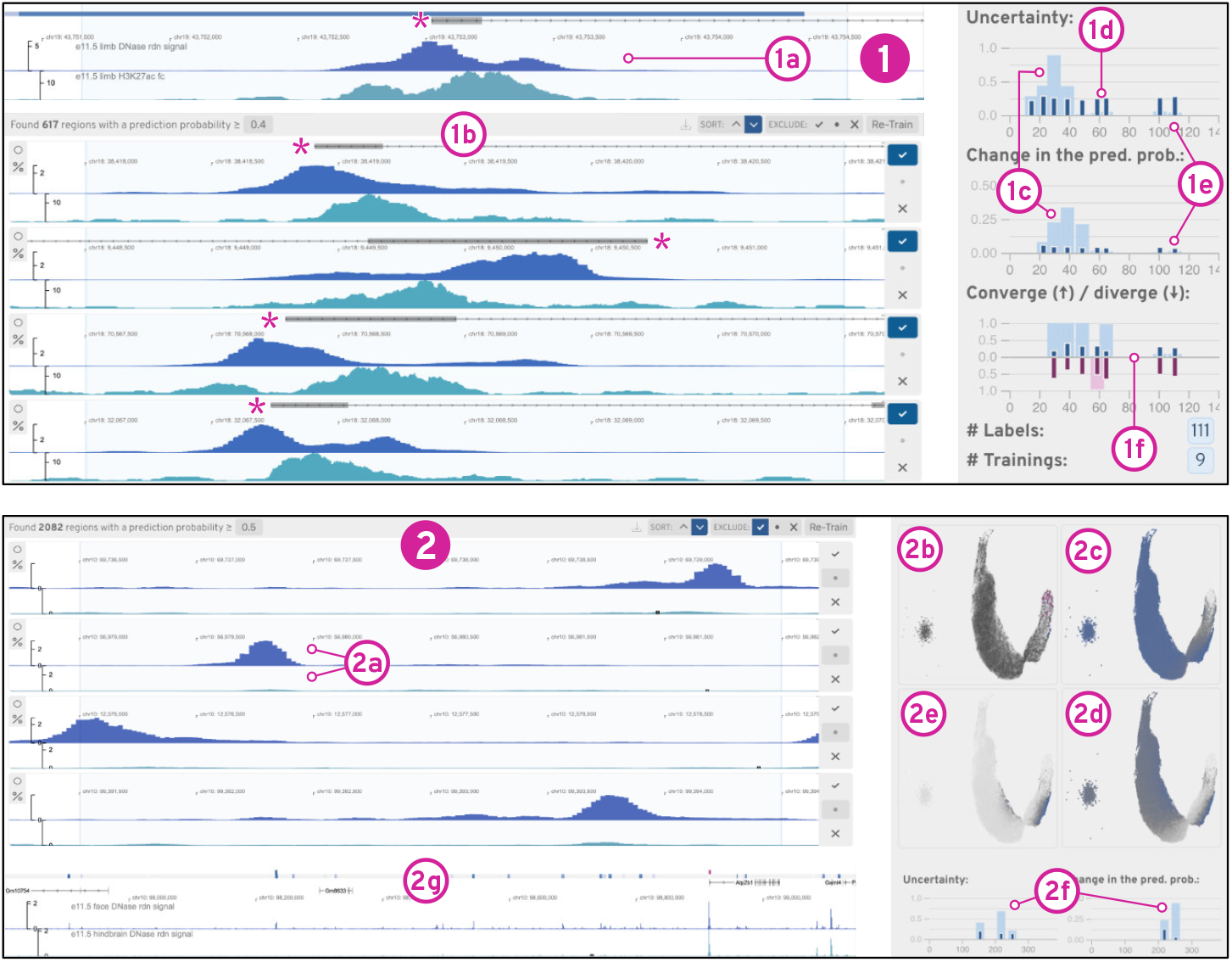
Findings from the Use Cases. (1) shows the result and progress of our asymmetrical peak detector training. In (2) we refined existing labels to build a differential peak detector (2a) after resolving an initially high uncertainty in the existing labels (2b-c).

**Figure 7:**
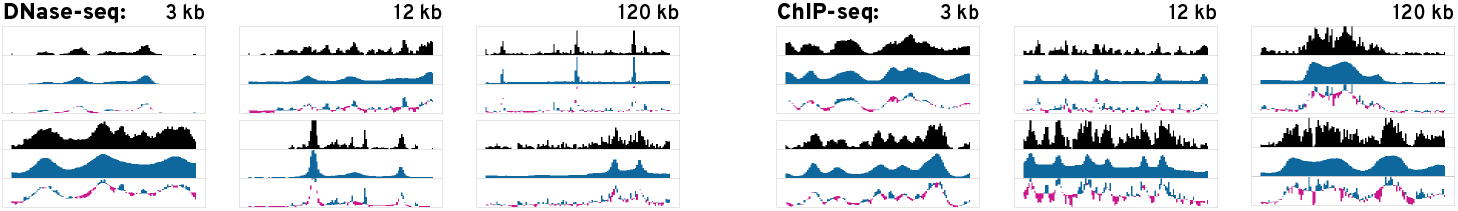
Example Reconstructions. The black bars represent the input and the blue bars visualize the reconstruction. The third chart plots the differences, where blue indicates missing data and pink indicates over-estimation of the reconstruction. As the window size increases the reconstruction gets less accurate. Since DNase-seq data contains less frequent peaks the reconstruction is more accurate.

### 5.2 Training Progress

To measure the classification process, we record overall uncertainty, the total change in prediction probability, and the divergence of prediction probabilities. The overall uncertainty is defined as the arithmetic mean of the class prediction uncertainty (Equation 3). The classifier’s uncertainty can indicate if the classifier learned to reliably detect a certain pattern type and if any progress is made over several iterations. The prediction probability change, which is defined as the arithmetic mean of the difference between the per-window prediction probability of the current and previous classifier, can provide insights about the impact of a training iteration. Finally, convergence is determined as the overall number of windows for which the prediction probability consecutively decreases or increases over the last 3 classifiers. All non-converging windows are considered diverging if the change in the prediction probability is larger than 0.01. The convergence and divergence rate can inform the user about the overall stability of the classifier. For formal definitions see Supplementary Table S1.

## 6 The User Interface

Based on the identified tasks T1-T6 (section 3), we have developed Peax to provide a visual interface for feedback-driven interactive visual pattern search and exploration of sequential data. The user interface consists of four main components shown in Figure 5.

### Query View

The system shows the selected search query or the highest ranked window by default in the query view (Figure 1) to provide content to the targeted pattern search (T4). This view is not static but allows the user to interactively browse the entire dataset (T1) for contextualization (Figure 5.1). The user can also choose to visualize additional metadata (Figure 5.1a) such as gene annotations to add further context (T4) during exploration. Additionally, after having trained a classifier for the first time, the view features a onedimensional heatmap track (Figure 5.1b) that color-encodes the prediction probability for regions in the data (T2). The main focus during navigation is on positive hits and we use a max-binning approach to show positively classified regions even when the user is zoomed out to view an entire chromosome (Figure 5.1a). In addition to the prediction probability, the query view shows two more types of region-based annotations: blue and pink rectangles indicate positive and negative labels (Figure 5.1c) that were defined by the user and yellow highlights selected windows (Figure 5.1d). Finally, the sequential data track can be superimposed with the reconstruction of the CAE model (Figure 1 and Figure 5.2e) for understanding which visual features are captured by the CAE (T6). Knowing how well the learned features represent the data can inform the user whether a search for a specific pattern is feasible. Therefore, this feature can be used for debugging when a pattern search fails. It is also useful during the development and testing of new (auto)encoder models.

### List View

The list view (Figure 5.2) visualizes several independent windows for labeling and comparison (T3). These windows are stacked vertically and are aligned to the query view for visual comparisons (Figure 1). Each window has three buttons (Figure 5.2g) to label it as interesting 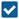, not interesting 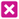, or inconclusive 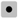. The two buttons on the left side of each window (Figure 5.2d) allow the user to select a window (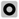) for separate comparison and to re-scale (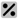) all currently visible windows to the associated window such that the y-scales of all windows are the same for value-based comparison (T3). By default, each window and the query view are individually scaled to the window’s minimum and maximum value to emphasize the pattern shapes. The list view consists of three permanent and one temporary set of windows that are organized under different tabs (Figure 5.2a). The *new samples* tab on the left is selected upon starting a new search and contains unlabeled windows that are sampled by the active learning strategy (section 5). The *results* tab in the middle lists the windows that are predicted by the classifier to match the query. The results tab includes two additional features to improve recall and debug data. First, it allows to dynamically adjust the probability threshold at which a window is considered a positive hit (Figure 5.2b). And in case the classifier and the user’s labels disagree, a pink button appears (Figure 5.2f) to inform the user about the potential conflicts. The *labels* tab on the right holds the set of already labeled windows. Additionally, when the user selects some windows a fourth tab is shown to the right of the labels tab allowing the user to compare the selected windows (T3).

### Embedding View

To provide an overview of the entire dataset and allow users to contextualize (T4) windows by their similarity in the latent space, Peax provides a view of the 2D-embedded windows (Figure 5.3). The embedding is realized with UMAP [53]. It takes into account the window’s latent representation and the user’s labels. Windows are represented as points that are either color-encoded by their label (Figure 5.3a) or by their prediction probability (Figure 5.3b). The view supports pan and zoom interaction as well as dynamic selection via a click on a dot or a lasso selection that is activated by holding down the shift key and the left mouse button while dragging over points (Figure 5.3a). Selecting windows is important for comparing (T3) local neighborhoods during exploration and for debugging already labeled windows.

### Progress View

To inform the user of the training progress and provide guidance on when to stop the training (T5), we visualize metrics for the uncertainty and stability (Figure 5.4) as defined in subsection 5.2. Each metric is applied separately to the entire set of windows and to the labeled windows. Visually this information is encoded as superimposed bars where the smaller but more saturated bars (Figure 5.4a) represent the metric in regards to the labeled windows. The wider bar represents all windows (Figure 5.4b), to indicate how well the trained classifier generalizes to the entire dataset. The y-scale is normalized to [0,1] and the x-scale stands for the number of labeled windows that were used to train a classifier. The diverging bar plot for the convergence metric simultaneously visualizes the convergence score in the upper part and the divergence score in the lower part. In general, as the training progresses one would expect each score to decrease. Large bars either indicate that the classifier is still uncertain or unstable. It is also possible that the target pattern changed or that the latent representation is not able to properly represent the features of the query.

### View Linking

The query, list, and embedding view are highly interlinked to foster contextualization (T4) and comparison (T3) of windows. For instance, moving the mouse over a window in the list view (Figure 5.2c) highlights the same location in the query view (Figure 5.1e) and the corresponding point in the embedding view (Figure 5.3d). The highlighted position of a window in the embedding view informs the user how windows compare to each other in terms of their latent representations and the shared mouse location highlights the windows’ spatial locality in terms of the underlying sequence.

## 7 Implementation

Peax’s code base is highly modular. The frontend application is implemented in JavaScript and uses React [18]. The bar charts are visualized with HiGlass [38]. The embedding view [41] uses Regl [46] for WebGL rendering. The Python-based backend server is built upon Flask [61] and can be configured via a JSON file. We use Scikit-Learn’s implementation [57] of the random forest classifier and persistently stored search results in a Sqlite [28] database. For training the CAEs we use Keras [8] with TensorFlow [1]. Documentation on how we trained our CAEs is provided in the form of iPython Notebooks [58]. All of the source code and detailed instructions on how to get started are available at https://github.com/Novartis/peax/.

## 8 Use Cases: Epigenomics

The human genome is a sequence of roughly 3.3 billion chemical units, called base pairs (bp), which encode for protein and RNA genes. But genes make up only less than 2% of the human genome, the remaining fraction contains the regulatory information for when and where genes are expressed. The regulatory information consists of genomic regions, called regulatory elements [55], and associated protein complexes. Gene expression is then controlled by modification of the degree of compaction of DNA and modification of the core proteins of the DNA and protein complex, called histones. These modifications are collectively called the “epigenome”. Studying the epigenome involves, for example, the genome-wide measurement of DNA accessibility (e.g., DNase-seq [13]), DNA protein binding and histone modifications (e.g., ChIP-seq [2]), gene expression (e.g., RNA-seq [45]), or spatial chromatin conformation (e.g., Hi-C [43]). Consortia like ENCODE [11], Roadmap Epigenomics [6], or 4D Nucleome [14] provide a large collection of these assays across many cell types and tissues. But while strict protocols and processing pipelines are applied, the variance between measurements and biological samples remains high and detecting regulatory elements is highly challenging [23, 59].

As our driving use case, we focus on the exploration of regulatory elements [55]. DNA accessibility and histone modifications are often used as proxies for detecting these elements, where peak-like patterns (Figure 6) show the strength of the epigenomic marks. Multiple datasets are typically explored in parallel, as multivariate sequential data, since no single method provides enough evidence to confirm the presence of a regulatory function. To explore such data we trained six CAEs [42] for 100 epochs on 343 histone ChIP-seq (modifications H3K4me1, H3K4me3, H3K27ac, H3K9ac, H3K27me3, H3K9me3, and H3K36me3) and 120 DNase-seq datasets, on 3 kilobase pairs (kb), 12 kb, and 120 kb windows, with bin sizes of 25, 100, and 1000 bp respectively. We chose the above mentioned window sizes (*l*) to capture local regulatory interactions as well as long-range regulatory domains. See Supplementary Tables S3–S5 for details.

### 8.1 Finding Peaks

We demonstrate how Peax can be used to find regulatory regions in epigenomic data. In recent work exploiting epigenomic data to detect regulatory enhancer patterns, Fu et al. [23] found that DNase-seq and histone mark ChIP-seq peak callers retrieve and rank peak patterns very differently and that the overall accuracy for finding regulatory elements is still quite low. Inspired by their findings, we first show how Peax can be used to build a classifier for asymmetrically co-occurring peak patterns from scratch. We then illustrate how existing peak annotations can be used as a starting point to find patterns of differentially activated regulatory regions.

### Building a Classifier From Scratch

We choose a lung DNase-seq and H3K27ac histone mark ChIP-seq dataset from ENCODE and encode it with our 3 kb DNase and ChIP-seq CAEs. We begin by selecting an asymmetrically co-occurring peak through brushing to start the search (Figure 6.1a) See our supplementary video and Supplementary Figure S22–S33 for complementary screenshots. We follow the left side of our workflow (Figure 3) and use our active learning strategy (section 5) to label 65 windows. The progress view (Figure 6.1c-f) shows that the uncertainty first increases (Figure 6.1c) and then gradually decreases (Figure 6.1d) before it stabilizes (Figure 6.1e), indicating that the classifier is learning to detect a certain pattern type. To get an overview of all windows, we compute the 2D embedding, which shows several clusters. Next, we switch the results, exclude our own positive labels, and find the classifier is able to detect patterns similar to our target. To assess the recall of our classifier, we invert the sort order of the results to find windows with a prediction probability close to 0.5. We find that these windows still match our target pattern and lower the threshold to 0.35. Peax now warns us about potential false-negative and false-positive conflicts, which we subsequently inspect and fix. Finally, we manually identify neighborhoods of windows showing instances of our query pattern using the embedding view. Using the prediction probability color encoding, we find a neighborhood of windows that are predicted to match our search target and select these windows using the lasso tool to improve the labeling further.

### Refine Existing Labels to Build an Improved Classifier

Peax also supports interactive refinement of existing labels. This time we search for differentially-accessible peaks in two DNase-seq datasets from face and hindbrain. Such peaks are shown [23] to be highly predictive of tissue-specific regulatory elements. Using algorithmically-derived peak annotations, we define positive labels as regions that have a reported peak annotation in the face dataset but not the hindbrain dataset and negative labels as regions that share peak annotations. See Supplementary Figure S34-S48 for complementary screenshots. We start by training a classifier with our pre-loaded labels. Since we have not specified a query region, Peax shows the region with the highest prediction probability in the query view. We find that the top results indeed show a pronounced peak in the top track and no peak in the bottom track (Figure 6.2a). Given the surprisingly high number of positive matches, we investigate the prediction probability landscape in the embedding view and see that almost every window has a probability higher than 0.5 (Figure 6.2c). After realizing that most results with a probability threshold close to 0.5 do not contain a differentially-accessible peak, we increase the probability threshold to 0.85. As a consequence, several incorrect labels appear as conflicts, which shows that the pre-loaded peak annotations are far from perfect. After resolving the conflicts and retraining the classifier, we observe a strong change in the average prediction probability and notice that the uncertainty for labeled windows decreased while the uncertainty for unlabeled data increased (Figure 6.2f), which is also reflected in the embedding view (Figure 6.2d dark-gray dots). Using the embedding view’s lasso tool, we assign labels local neighborhoods with high uncertainty, which leads to a notable reduction in the overall uncertainty (Figure 6.2e). Finally, using query view we explore the entire dataset to assess the spatial location of the found windows (Figure 6.2g).

## 9 Evaluation

### 9.1 Reconstruction

First, we study the reconstruction quality with data from our use case (section 8) to assess the general performance of our CAE models. While the reconstruction is not expected to be perfect given the 12-fold compression, it should capture the main visual properties to be useful for visual similarity search. Figure 7 exemplifies the reconstruction quality of the six CAEs from our use case (section 8). More example reconstructions are available in Supplementary Figure S3–6. It appears that the model is able to learn the visual properties of the epigenomic datasets. Most of the salient peaks are captured by the CAEs. As the window size and resolution increase from 3 kb over 12 kb to 120 kb (with a binning of 25 bp, 100 bp, and 1000 bp) the reconstruction quality decreases. Hence, the model is not able to encode high-frequency variations. Numerically this is captured by an increase in the reconstruction loss for larger window sizes. Since absolute loss is hard to interpret, we report *R*2 scores to assess how much variability is captured by the reconstruction, where a score of 1 stands for a perfect reconstruction. All scores are computed on the test data. For the CAEs trained on DNase-seq data the *R*^2^ is .98, .90, and .78 for 3 kb, 12 kb, and 120 kb windows respectively. Similarly, the *R*2 scores for the CAE trained on histone mark ChIP-seq data are .84, .69, and .73 for 3 kb, 12 kb, and 120 kb windows respectively.

**Figure 8:**
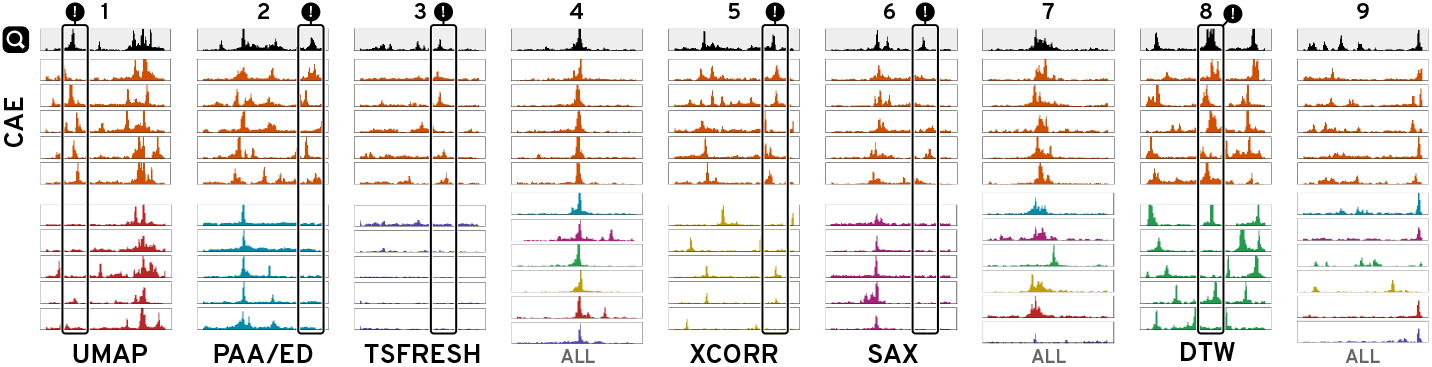
Similarity Comparison. Given a query (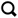) at 12 kb, we ran a 5-nearest neighbors search with our model (CAE), four distance-based methods (PAA/ED, SAX, DTW, and zero-normalized cross-correlation (XCORR)), and two feature-based methods (UMAP and TSFRESH). All techniques are able to detect the most distinct visual feature but many fail to match secondary features, as highlighted with (!). For simple patterns (4) many methods are equally good and we only show the top results. See Supplementary Figure S15 for all results.

### 9.2 Similarity Comparison

Knowing that the CAEs are able to successfully reconstruct most of the visual features, we wanted to know whether the learned representation is more effective for visual similarity search in terms of the perceived similarity compared to other techniques. Given a DNase-seq dataset from our use case, we manually identified 9 diverse query patterns. Using a 5-nearest neighbor search, we subsequently retrieved the 5 windows that are closest to the queries using the latent representation of our CAEs and six other techniques (Supplementary Table S6). ED [19], SAX [44], DTW [5], and zero-normalized cross-correlation (XCORR) are four distance-methods we compare against. Additionally, we included UMAP [53] and TSFRESH [9] as two feature-based techniques. Figure 8 and S15 shows the nine queries for 12 kb windows together with the 5-nearest neighbors of every method. Subjectively, the results from our model consistently find more similar patterns on average. While simple pattern, like Figure 8.4, are well captured by several techniques, every other method has at least one instance where the five 5-neighbors are missing important visual features compared to our CAE (Figure 8 “!”). Supplementary Figure S16 and S17 show the comparison for 3 kb and 120 kb windows.

To determine whether our subjective findings hold true, we evaluated the similarity in an online user study on Amazon Mechanical Turk. For each of the 27 patterns (9 patterns × 3 window sizes), we generated an image showing the 5-nearest neighbors for each technique together with the query pattern (Supplementary Figure S18). We asked the participants to “select the group of patterns that *on average* looks most similar and second most similar to the target pattern”. The order of target patterns and search results were fixed but we randomized the order in which techniques were shown. We asked for the most and second most similar group as a pilot study revealed that the task can be hard when there are two or more groups with similarly good results. In total, participants had to make 18 choices by comparing 7 × 9 groups of patterns. We paid $1 for an estimated workload of 6 minutes. Participation was restricted to master workers with a HIT approval rate higher than 97% and at least 1000 approved finished HITs. In total, we collected 75 responses. We hypothesize (H1) that our CAEs outperform other techniques by finding more similar patterns on average.

Using a Pearson’s chi-squared test, we find significant differences between the techniques (3 kb: *χ*^2^(1,N=25)=206.4, *p* <.001; 12 kb: *χ*^2^(1,N=25)=284.5, *p* <.001; 120kb: *χ*^2^(1,N=25)=319.0, *p* <.001). In a post-hoc analysis using Holm-Bonferroni-corrected [31] pairwise Pearson’s chi-squared tests between our and the other six methods (Figure 9 and S20), we find that our CAEs retrieves significantly more often patterns that are perceived most or second most similar to the target compared to SAX (3 kb: *χ*^2^(1,N=25)=19.8, *p* <.001; 12 kb: *χ*^2^(1,N=25)=44.6, *p* <.001; 120kb: *χ*^2^(1,N=25)=76.8, *p* <.001),DTW (3 kb: *χ*^2^(1,N=25)=57.3, *p* <.001; 12 kb: *χ*^2^(1,N=25)=49.5, *p* <.001; 120kb: *χ*^2^(1,N=25)=41.4, *p* <.001), UMAP (3 kb: *χ*^2^(1,N=25)=23.6, *p* <.001; 12 kb: *χ*^2^(1,N=25)=53.4, *p* <.001; 120kb: *χ*^2^(1,N=25)=100.3, *p* <.001), TSFRESH (3 kb: *χ*^2^(1,N=25)=126.2, *p* <.001; 12 kb: *χ*^2^(1,N=25)=134.4, *p* <.001; 120kb: *χ*^2^(1,N=25)=117.4, *p* <.001), and XCORR (3 kb: *χ*^2^(1,N=25)=72.0, *p* <.001; 12 kb: *χ*^2^(1,N=25)=97.7, *p* <.001; 120kb: *χ*^2^(1,N=25)=122.0, *p* <.001). We find no significant differences between CAE and ED. Therefore, H1 does not hold true given the strong performance of ED. In general, we found that for simple patterns it is more likely that any method performs well.

**Figure 9:**
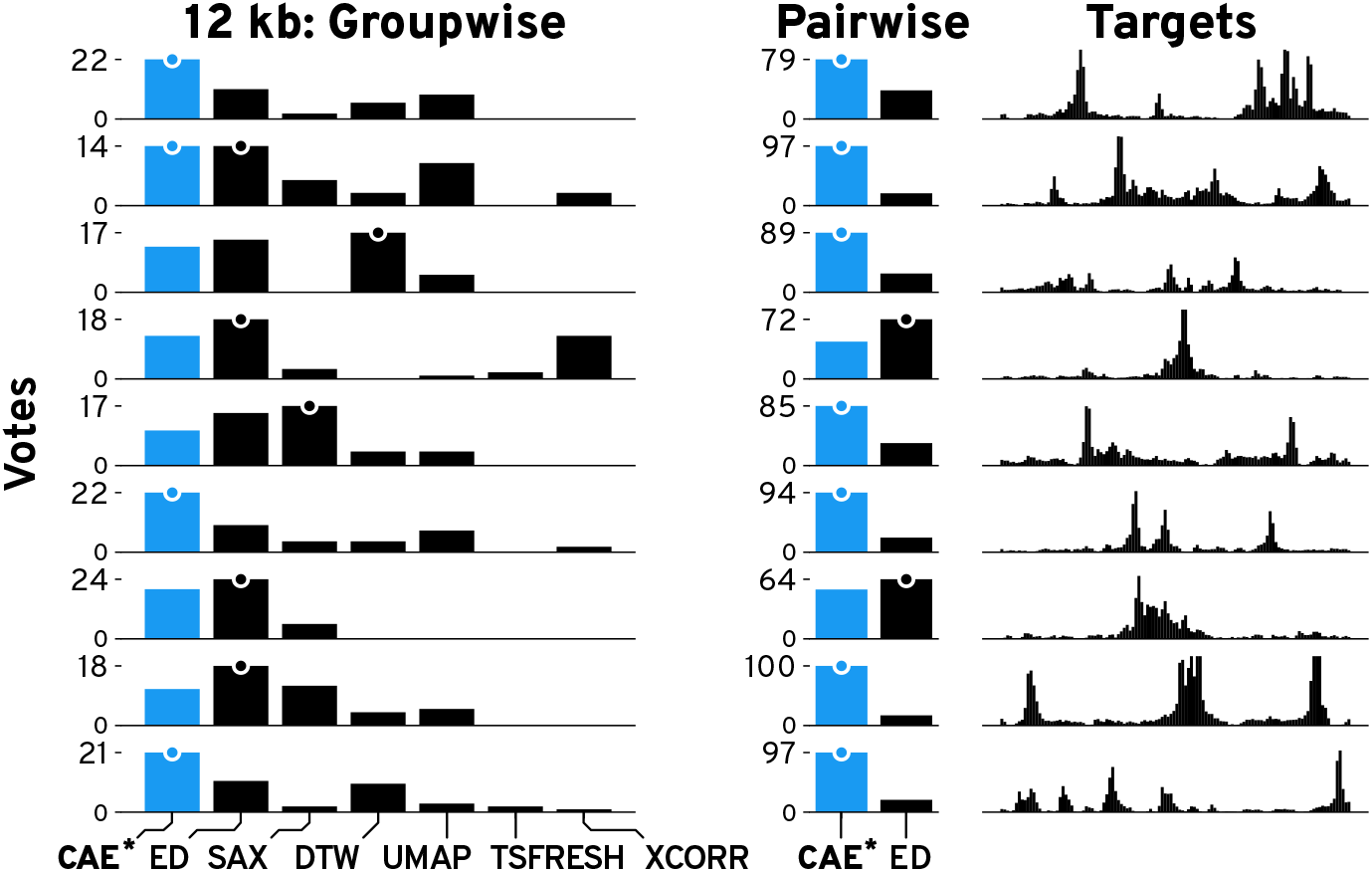
Results of the Similarity Comparison. Bars show the votes per technique per target pattern for 12 kb patterns. More votes are better. The highest votes are indicated by a dot. Results for our method (CAE) are drawn in blue.

We were surprised by some results where ED outperformed our model, e.g., Supplementary Figure S15.8. We hypothesize that for complex patterns groupwise comparison can be too challenging. To gain a more detailed picture, we conducted a follow-up study on pairwise similarity search between the 5-nearest neighbors of just our model and ED. For each of the 27 patterns, we generated images showing the target pattern together with one pattern found with CAE and ED. The pairing of patterns is determined by the order in which they were found, e.g., the first nearest neighbor from CAE is paired with the first nearest neighbor from ED, etc. The order in which the patterns are drawn was randomized but the order of targets was fixed as in the first study (Supplementary Figure S19). We asked the participants to “select the pattern that looks more similar to the target” for each of the 5-nearest neighbor pairs. This resulted in a total of 45 comparisons (9 patterns × 5 pairs). We paid participants $1 for an estimated workload of 6 minutes. Participation was limited to the same population as in the first user study and we again collected 75 responses. We hypothesize (H2) that our CAE model outperforms ED in pairwise comparisons on average. However, we also expect mixed results for simple patterns. Using the same analysis from the first user study, we found that across all 5-nearest neighbors, our model retrieves significantly more similar patterns compared to ED for 3 kb (*χ*^2^(1,N=25)=70.3, *p* <.001), 12 kb (*χ*^2^(1,N=25)=171.5, *p* <.001), and 120 kb (*χ*^2^(1,N=25)=85.2, *p* <.001) window sizes. Thus, we can accept H2. See Supplementary Figure S20 and S21 for detailed votes.

### 9.3 Usability & Usefulness

We conducted an in-person user study with 8 domain experts (P1-8) to investigate the usability and usefulness of Peax. All experts are computational biologists who work with epigenomic data on a regular basis (Supplementary Table S9). In each individual one hour-long session, we first introduced Peax using our supplementary video, summarized the primary use cases for genomics, and provided a brief demo of features not covered in the video (15 minutes). Afterward, each expert completed two tasks on building and evaluating pattern classifiers. The experts had 15-20 minutes to complete each task, and we asked them to think aloud while we recorded the screen and audio. See Supplementary Table S8 for a detailed description of the procedure.

#### Task 1

We asked the participants to build a classifier to find windows that contain a peak pattern in the first two out of three datasets (Supplementary Figure S50). To assess the performance of the classifiers, we simulated three ChIP-seq experiments (Supplementary Table S7) with ChIPsim [33] and used our 3kb ChIP-seq CAE for encoding. As a guideline, we mentioned that 50-100 labels are typically needed to produce reasonable results. See Supplementary Table S11 for the participants’ actions. All but one expert started by sampling and labeling windows for the first 5-10 minutes before they switched to the results. In the results tab, they typically labeled windows of the first few pages. Six experts also labeled windows that were close to the prediction probability threshold. While assessing the results, six experts noticed potential conflicts highlighted by Peax (Figure 5.2f) and adjusted the labels if necessary. Also, two experts increased the prediction probability threshold (Figure 5.2b) after inspecting the conflicts. Finally, six experts computed the embedding view and explored windows within this view. Interestingly, one expert (P2) switched to the results after only one round of training. As the results were not good yet, P2 continued labeling and retraining the classifier in the results tab.

We compared the ground-truth labels from our simulated data against the expert-generated labels and found (Supplementary Figure S51) high sensitivity of 0.98 (SD=0.02) and specificity of 0.95 (SD=0.07) on average. P2 and P5 had a relatively high number of *false-positive* labels, which indicates that their imagined target pattern might have differed from ours. Next, we evaluated the performance of the classifiers in terms of the area under the receiver operating characteristic curve (AUC-ROC), average precision (AP), and Matthews correlation coefficient (MCC) with a probability threshold of 0.5. When training the classifiers on our experts’ labels, we found an average AUC-ROC of 0.93 (SD=0.04), AP of 0.56 (SD=0.09), and MCC of 0.49 (SD=0.12) (Figure 10). The MCC scores for classifiers trained on P2’s and P5’s labels are significantly lower, as their search targets diverged. We then compared the performance against classifiers trained on ground truth labels sampled with our active learning strategy to study the impact of a human-in-the-loop. Interestingly, we found that on average, AUC-ROC (+0.13) and AP (+0.14) are higher for classifiers trained on the experts’ labels (see blue versus gray bars in Figure 10), which indicates that the expert-steered training process is more effective. The differences for the MCC scores are minor, except for P2 and P5, which indicates that a probability threshold of 0.5 might not be ideal.

**Figure 10:**
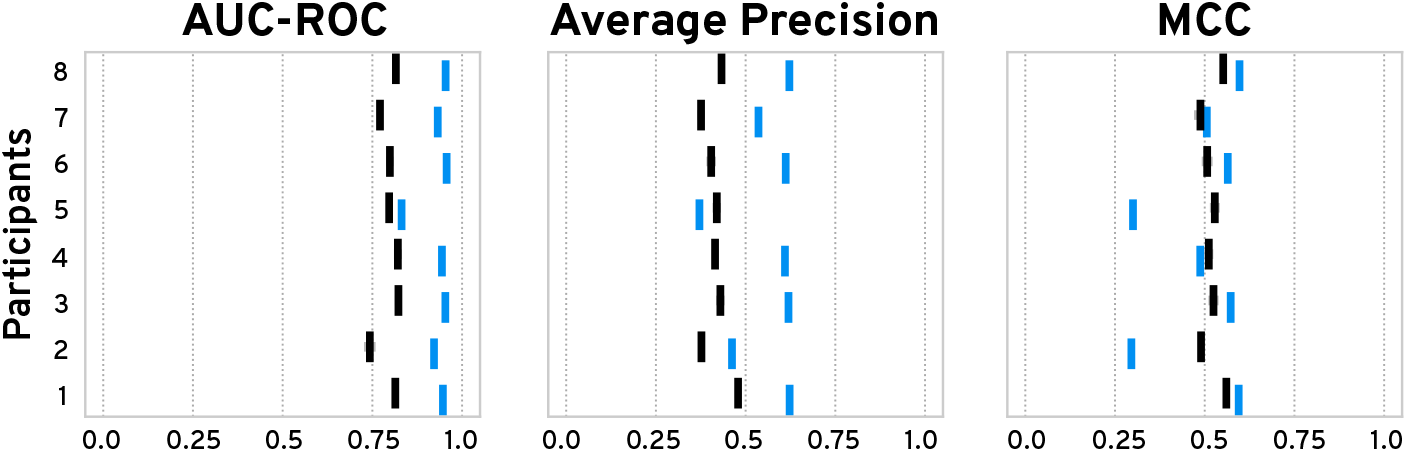
User Study Task 1 Classifier Performance. We trained our classifier 10 times on the experts’ labels (blue) and on ground-truth labels sampled with our active-learning strategy (black). Vertical and horizontal bars show the mean and standard error.

#### Task 2

We asked the participants to evaluate and improve a classifier trained on a set of pre-loaded labels. The dataset and labels were taken from our second use case scenario (section 8) to assess Peax’s usability on a real-world dataset. See Supplementary Table S12 for the participants’ actions. In the beginning, all experts acknowledged that the classifier correctly predicts differentially-accessible peaks and that they don’t have to label new samples. Subsequently, all experts assigned labels to the top results and noticed that the number of matching windows was too high. Interestingly, different actions made the experts question the number of results. Three experts noticed that results with a prediction probability close to the threshold did not match the target pattern. Two experts discovered that a lot of regions had a prediction probability close to 0.5 after examining the embedding view. Another two experts investigated the threshold after inspecting conflicts. Finally, one expert decided to retrieve new samples and realized that the classifier is uncertain about regions with low signal, that typically appear in high numbers. Eventually, all experts increased the threshold to reduce false positives results.

In general, through the two tasks and the post-study questionnaire (Q7 and Q9) we found that Peax’s interactive pattern search is an effective and useful new exploration approach. None of the experts is aware of any tool that supports a similar pattern exploration approach. P6 and P8 mentioned that they currently have to browse and annotate their data manually. We also found that Peax is easy to learn and use. The 15-minute introduction was sufficient to enable the experts to successfully build and refine pattern classifiers. Expert P1 mentioned that “for having just seen a quick video and a couple of quick instructions, it’s pretty easy to work through”. After the study, we addressed smaller usability issues that were mentioned by the experts (Supplementary Table S13). However, use case-specific guidelines would be helpful to lower the initial learning curve, as mentioned by P2, P3, and P8. While the manual labeling and training process requires more work than a purely computational search, the classifier comparison from Task 1 suggests that a human-in-the-loop can make the labeling process more effective. Also, P8 noted that the labeling “is really awesome to play around with. It’s like playing a game.”, and we also see increasing trust in the results of Peax compared to fully automatic peak and feature detectors (see Figure S52 Pre-Study Q5 versus Post-Study Q5).

## 10 Discussion

Instead of using deep learning techniques to predict the presence of a specific pattern type in a fixed combination of datasets, we use deep learning to augment human intelligence to search for patterns in multivariate sequential data. Our approach improves the performance of visual pattern search by providing a learned latent representation together with an actively-learned classifier. Having a tool for general exploration of large sequential datasets by pattern similarity allows to ask new questions and find previously undetectable patterns, and complements efforts for highly specialized pattern detectors.

While the overall results of our user study suggest that the user-steered learning process leads to more effective classifier training, subjective interpretation of the target pattern can influence the classifier’s performance strongly. Visualizing the impact of a label on the classifier might help to stabilize the labeling between users. Despite its complexity, we show that Peax is easy to learn and use, given a short introduction to Peax’s general search approach. While all participants actively used at least two different labeling strategies to broaden the exploration, more in-depth training might be required to utilize all functionality of Peax. Carefully-designed notifications could also inform the user if their exploration strategy is too narrow.

Peax is not limited to any of the specific autoencoders that we presented in this paper. In fact, the system is explicitly designed to facilitate extension with custom encoders, and we hope that analysts will use Peax as a tool to evaluate their own (auto)encoders for pattern search. Thus, it is up to the user to normalize the data before loading the data into Peax. Moreover, while the use case presented in this paper is about epigenomic data, the ideas apply equally to other sequential or time series data, given that the autoencoder is purely data-driven and does not require any prior knowledge. For other use cases, the most important consideration is the window size. The windows need to fully include the target pattern to allow the CAE to extract relevant features. In our current implementation, we use a random forest classifier given its superior performance (Supplementary Figure S49). But Peax supports any classifier that implements Scikit-Learn’s API [57].

While we found that the reconstruction is not perfect, it is hard to tell what a “perfect” reconstruction in regards to visual pattern search might be. In general, our CAEs work best at encoding distinct patterns, e.g., medium-frequent peaks with a noticeable difference in magnitude compared to the background signal. High-frequency patterns are smoothed out by the CAE, which can be beneficial when the underlying phenomena is independent of the background noise. While there are legitimate reasons to assess the “randomness” of the background noise, like in anomaly detection, systems supporting such use cases should not be steered visually by humans, as judging randomness is a hard task [69,70] for humans. In general, Peax’s ability to find patterns is bound by the extracted features of the input data. As we aim for generic search, the unsupervised feature learning employs a general loss function, which emphasizes high compared to low-magnitude differences. As a consequence, a feature that is not visible with our eyes stays undetected by the CAEs. To address specific search scenarios, the user can provide a custom (auto)encoder model.

Currently, Peax scales well to up to one million windows. To ensure interactivity, Peax preprocesses the data, which takes about 25–30 minutes for one million windows on a regular laptop. The main limiting factor is the computation of the nearest neighbors and UMAP embedding. For datasets with more than 100,000 windows, we switch to an approximate nearest-neighbor algorithm. Cluster-based data preprocessing can further reduce the startup time for larger datasets. Visually, the main limitation is the embedding view, which works smoothly until one million dots but can slow down above 1.5 million dots. For larger datasets, interactive filtering techniques could alleviate overplotting and rendering issues. Finally, the amount of labels needed to train a reasonable classifier depends on the complexity of the search query. As an example, roughly 25–50 labels can be enough to retrieve reasonable results if the latent representation effectively captures a single-dataset query for a distinct pattern. This number can increase linearly in a multi-dataset search but ultimately depends on the complexity of the datasets.

Finally, the choice of the similarity search metric should depend on the data type and search goals. The larger the data, the more likely it is that different instances of the same pattern type exist. If the goal is to find the top *k* hits than many different techniques will likely yield equally good results. If, on the other hand, one seeks to find all pattern instances or the data are highly variable then a feature-based approach can be more effective as shown by our evaluations and similarity comparisons.

## 11 Conclusions and Future Work

We have presented Peax, a novel technique and tool for interactive visual pattern search in sequential data. We found that our convolutional autoencoder models learn a latent representation that is more effective at similarity search than existing techniques. In future work we want to compare the performance of different autoencoders such as different variational autoencoders, recurrent autoencoders, and other deep learning approaches such as generative adversarial networks. Especially the latter would allow visual judging of the encodings to improve debugging and sensemaking. In addition to the encoding efforts, there are several avenues to extend the active learning strategy. For example, when training a classifier from an existing set of peak annotations, a deep-learningbased strategy similar to Gonda et al. [26] might prove to be superior. Finally, in our in-person user study, we found that the articulated and imagined target pattern can differ between people. An exciting avenue for future work is to systematically test the variance of human labels within and between different subjects. Also, it would be interesting to expand Peax to a multi-user system and study the impact of multiple users on the labeling and learning process.

## Acknowledgement

We wish to thank all participants from our user studies who helped us evaluate Peax. We also like to express our gratitude to Mark Borowsky for his ideas and support. This work was supported in part by Novartis Institutes for BioMedical Research and the National Institutes of Health (U01 CA200059 and R00 HG007583).

