## Supplementary Material for "Peax Interactive Visual Pattern Search in Sequential Data Using Unsupervised Deep Representation Learning"


Fritz Lekschas

Harvard School of Engineering and Applied Sciences

Brant Peterson

Novartis Institutes for BioMedical Research

Daniel Haehn

Harvard School of Engineering and Applied Sciences
University of Massachusetts Boston

Eric Ma

Novartis Institutes for BioMedical Research

Nils Gehlenborg

Harvard Medical School

Hanspeter Pfister

Harvard School of Engineering and Applied Sciences

### Supplementary Figures

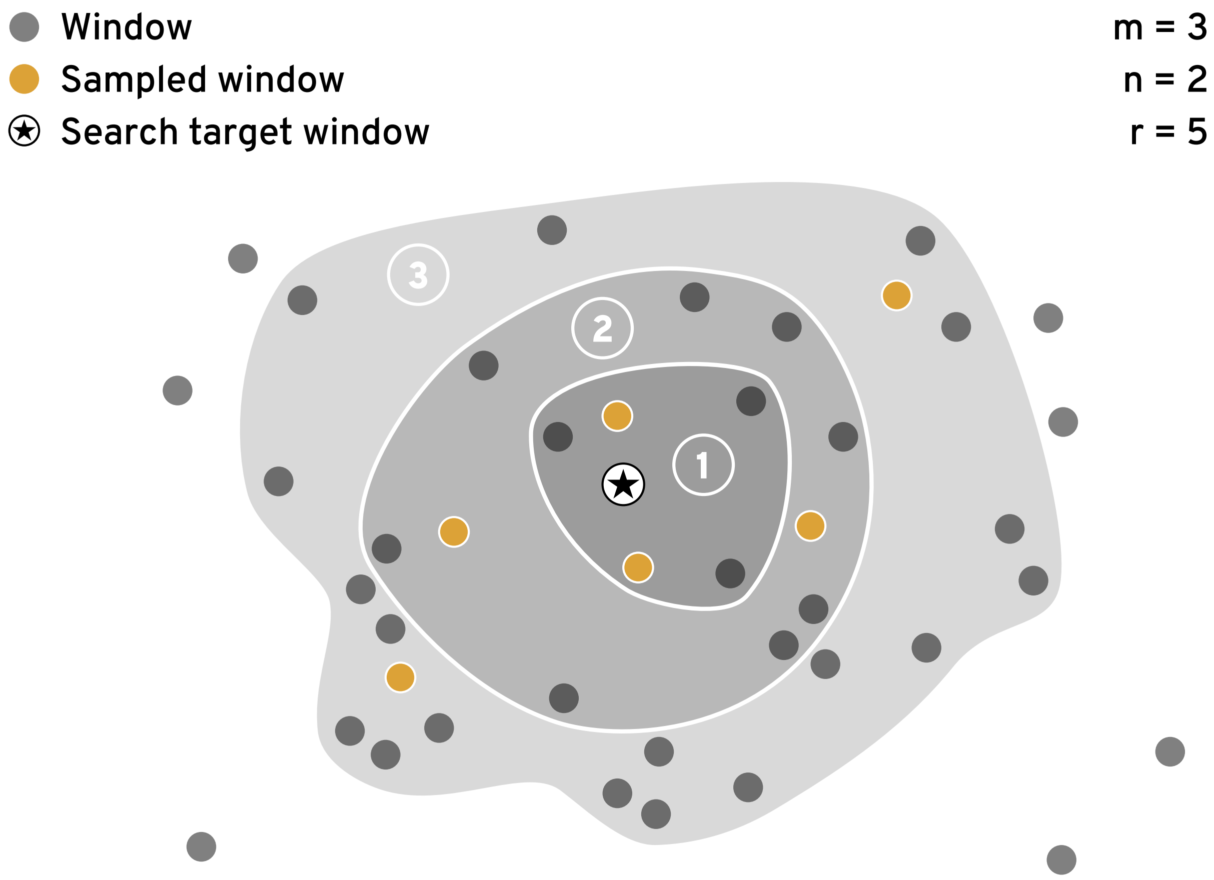


Figure S1: **Initial sampling.** An illustration of the initial sampling strategy in a 2D latent space. In this example we conduct $m = 3$ rounds of sampling. In every round, we iteratively sample $n = 2$ windows from the currently active search neighborhood (indicated by the shaded background) that are located in dense neighborhoods and far from already sampled windows. The search neighborhood includes the $r$ windows that are closest to the search query. The "radius" $r$, that is initially set to 5, is doubled in every iteration. Finally, to avoid sampling from the same neighborhood multiple times, windows that appeared in previous search neighborhoods are excluded from subsequent rounds. Our rationale is to select a diverse set of sample while still ensuring that the sample set includes positive windows that are very close to the search target. This is important for balancing the label set and to keep the user motivated during the initial labeling.

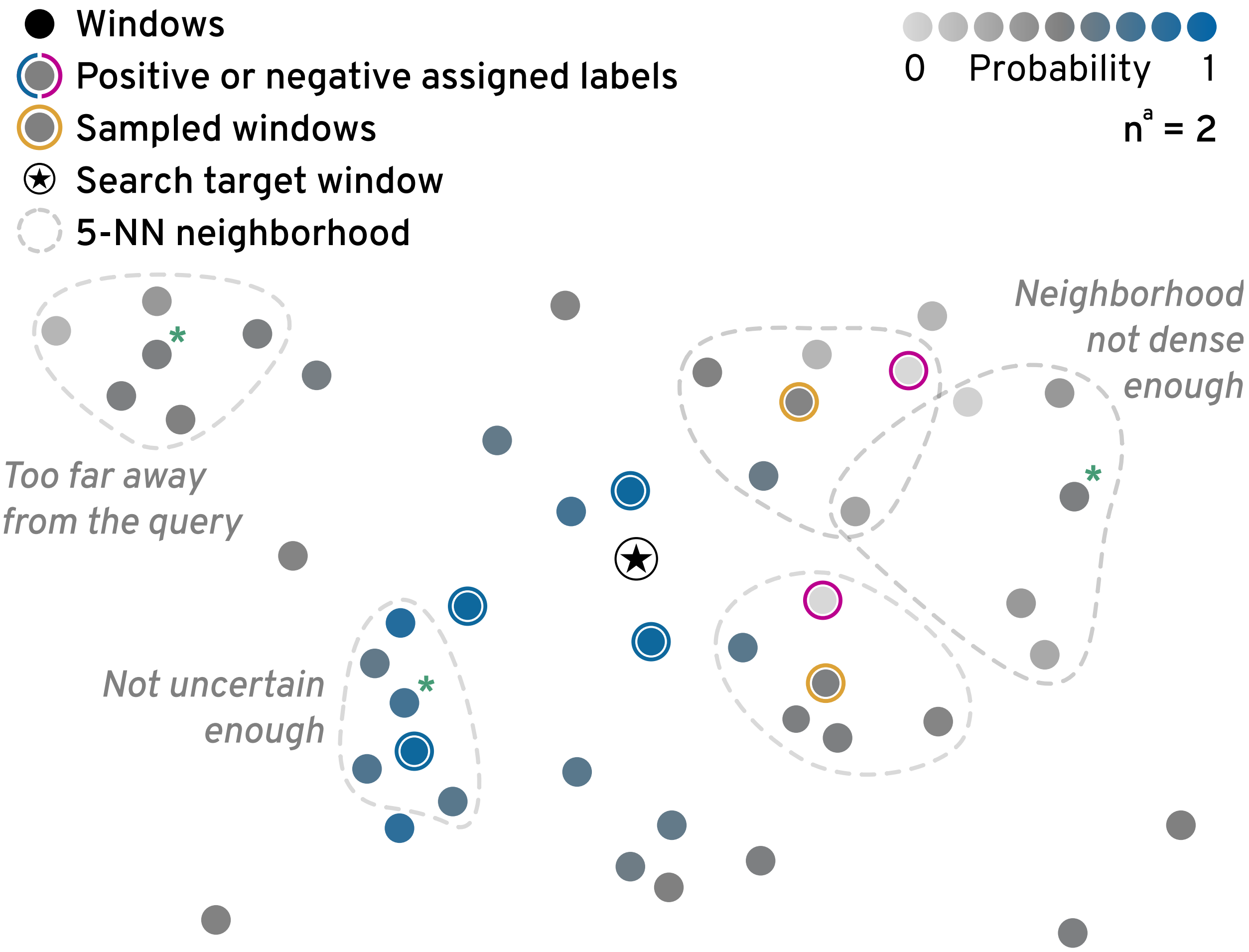


Figure S2: **Initial sampling.** An illustration of the uncertainty sampling strategy in a 2D latent space. In this example, we have already labeled 6 windows as indicated by the blue and pink outline around 6 dots. We are not about to sample $n^a = 2$ windows. The goal is to sample windows that are close to the target, are located in dense $k$-NN neighborhoods, are far away from already sampled windows, and have high uncertainty. In this example, we samples the two windows with the yellow outline since they are both relative close to the target, highly uncertain, and in dense neighborhoods. Other potential candidates, which are highlighted with a green asterisks, are not sampled because they are either too far away from the search query, are not uncertain enough, or are too far away from their 5-nearest neighbors.

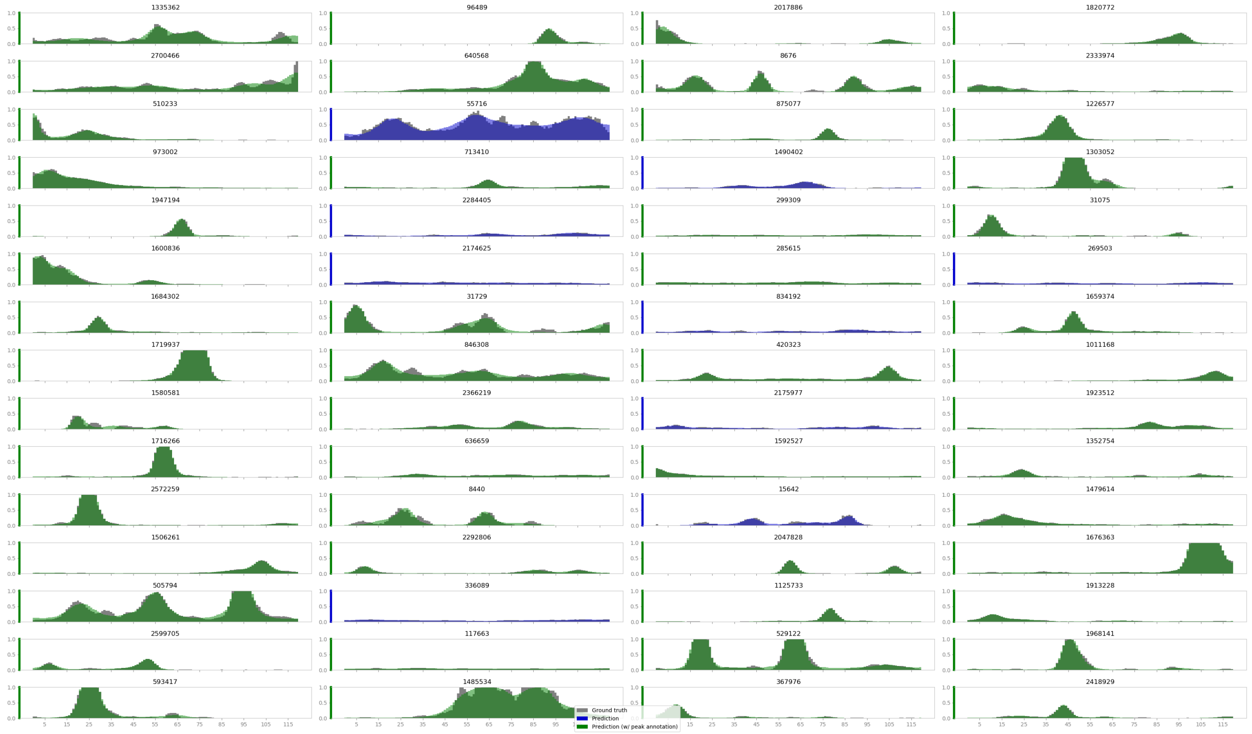


Figure S3: **Reconstruction of the 3 kb DNase-seq CAE**. We show a randomly chosen set of non-empty windows from the test set. Gray bars indicate the raw data and green and blue superimposed bars show the reconstruction. Green bars indicate windows that contain at least one computationally-derived peak annotation from the ENCODE project.

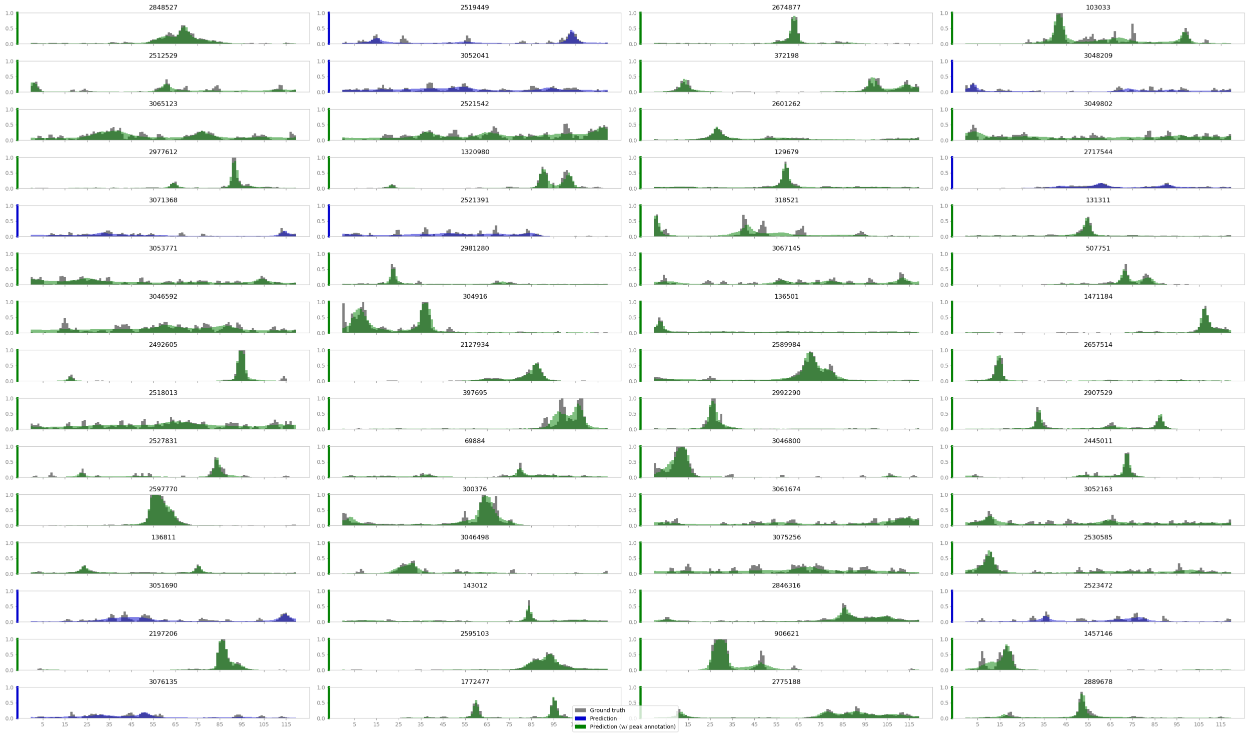


Figure S4: **Reconstruction of the 12 kb DNase-seq CAE**. We show a randomly chosen set of non-empty windows from the test set. Gray bars indicate the raw data and green and blue superimposed bars show the reconstruction. Green bars indicate windows that contain at least one computationally-derived peak annotation from the ENCODE project.

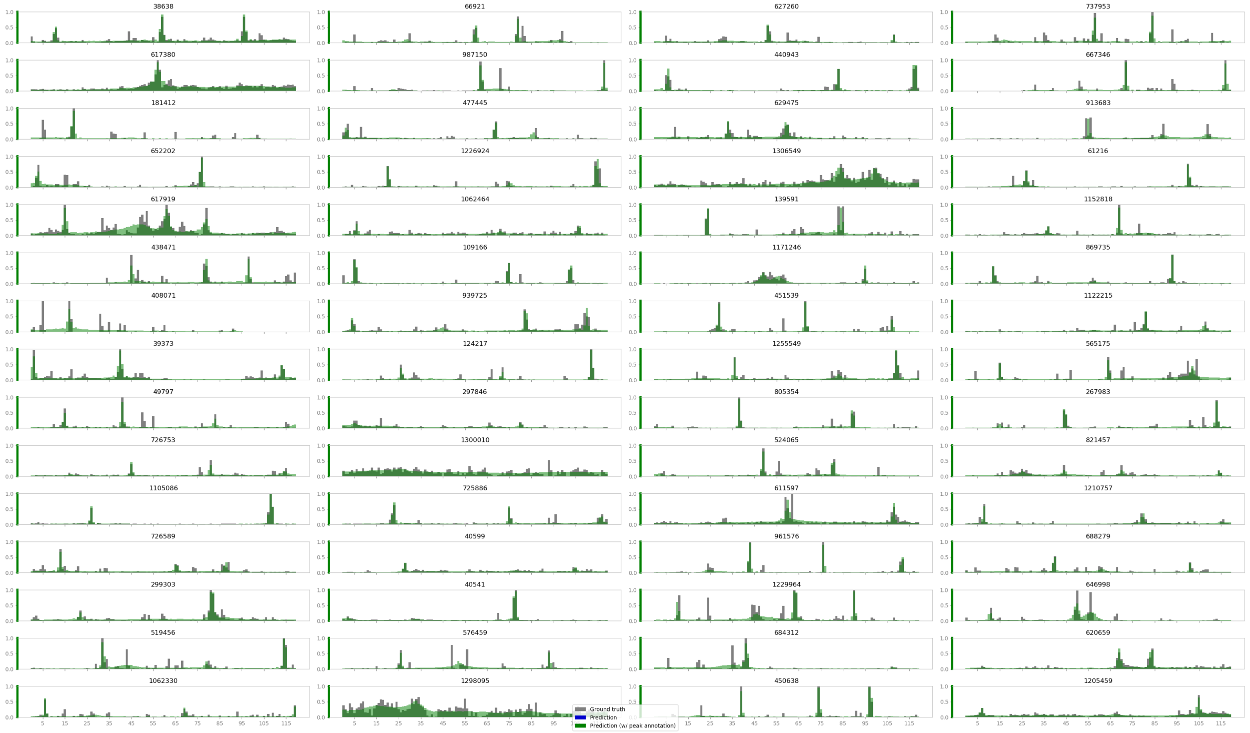


Figure S5: **Reconstruction of the 120 kb DNase-seq CAE**. We show a randomly chosen set of non-empty windows from the test set. Gray bars indicate the raw data and green and blue superimposed bars show the reconstruction. Green bars indicate windows that contain at least one computationally-derived peak annotation from the ENCODE project.

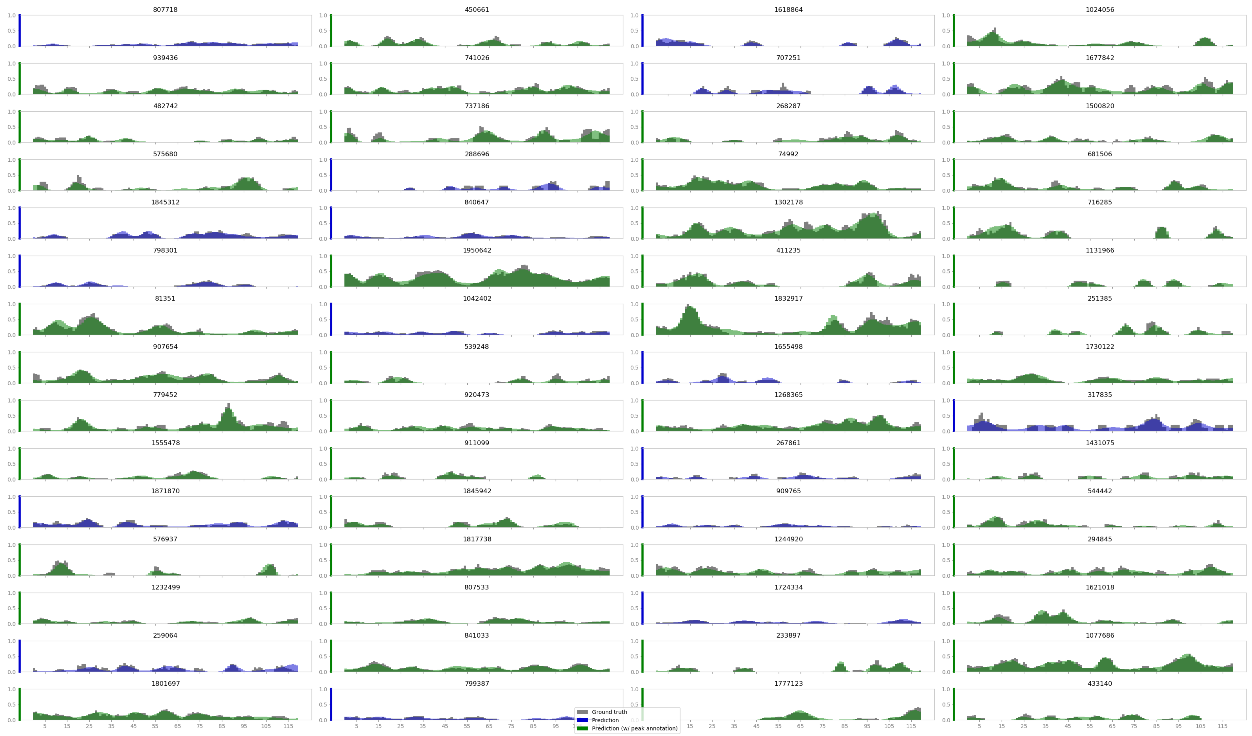


Figure S6: **Reconstruction of the 3 kb histone mark ChIP-seq CAE**. We show a randomly chosen set of non-empty windows from the test set. Gray bars indicate the raw data and green and blue superimposed bars show the reconstruction. Green bars indicate windows that contain at least one computationally-derived peak annotation from the Roadmap Epigenomics project.

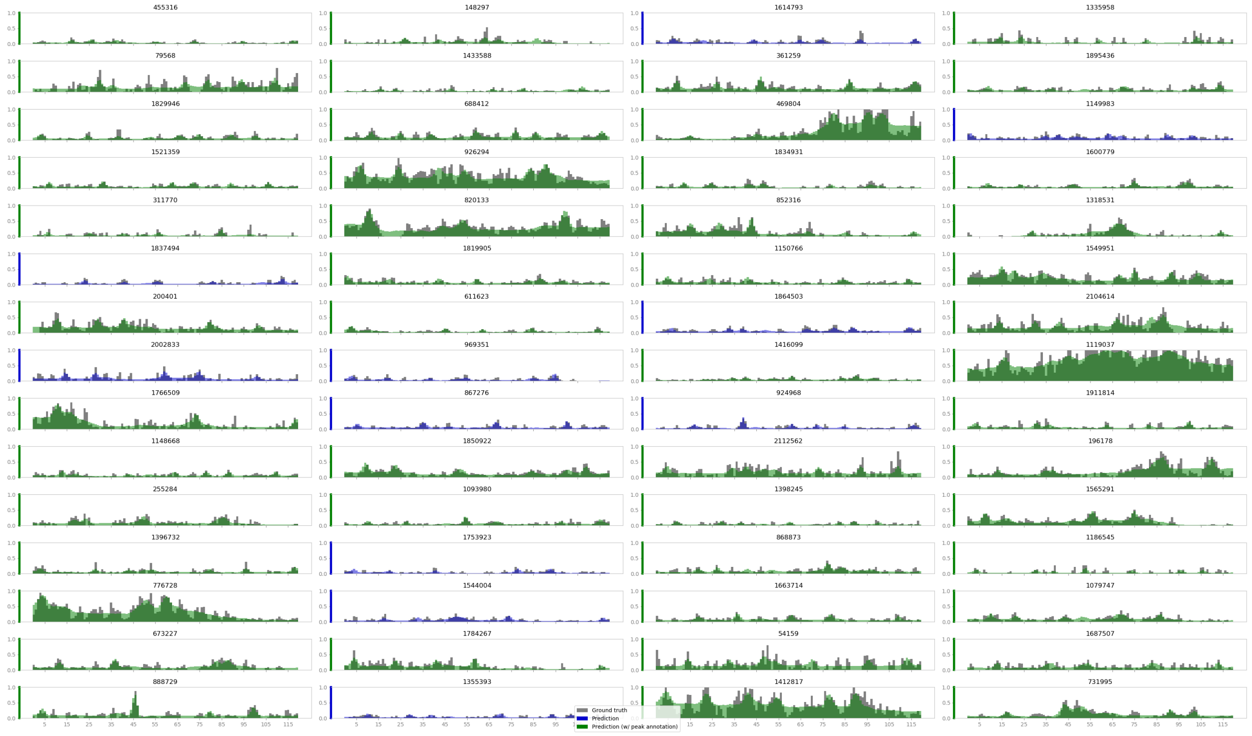

Figure S7: **Reconstruction of the 12 kb histone mark ChIP-seq CAE**. We show a randomly chosen set of non-empty windows from the test set. Gray bars indicate the raw data and green and blue superimposed bars show the reconstruction. Green bars indicate windows that contain at least one computationally-derived peak annotation from the Roadmap Epigenomics project.

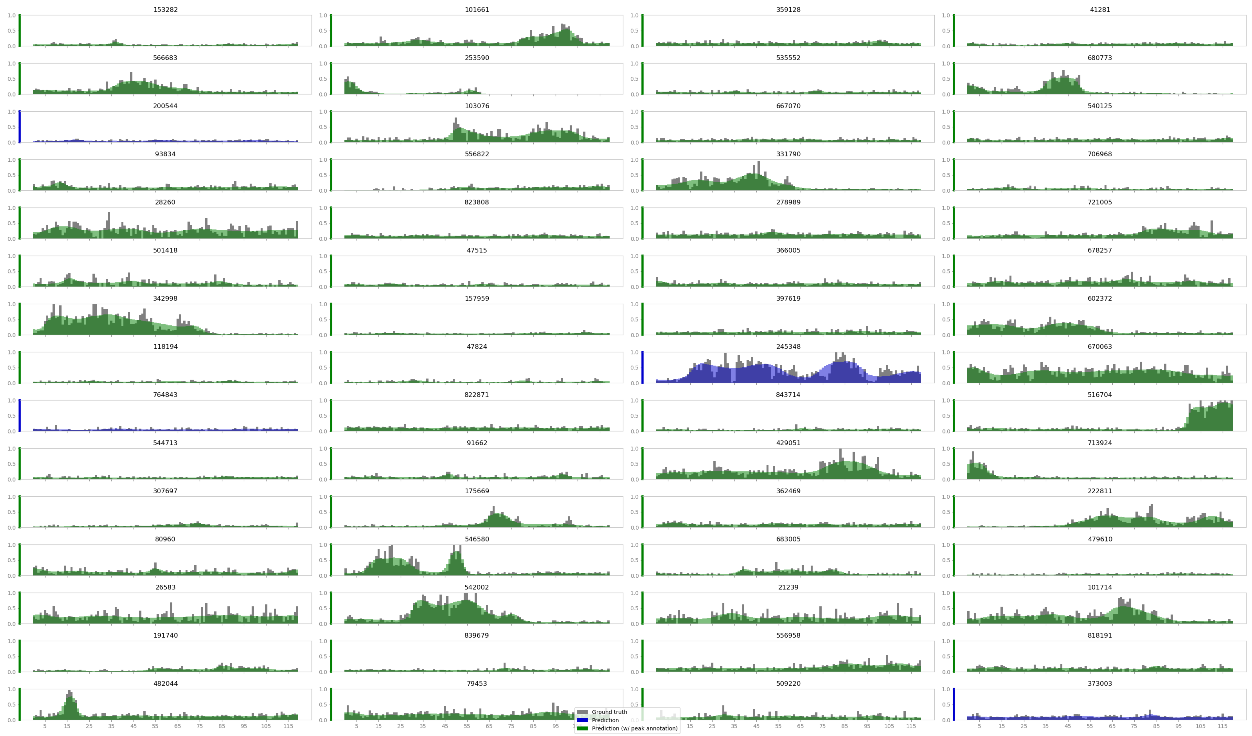


Figure S8: **Reconstruction of the 120 kb histone mark ChIP-seq CAE**. We show a randomly chosen set of non-empty windows from the test set. Gray bars indicate the raw data and green and blue superimposed bars show the reconstruction. Green bars indicate windows that contain at least one computationally-derived peak annotation from the Roadmap Epigenomics project.

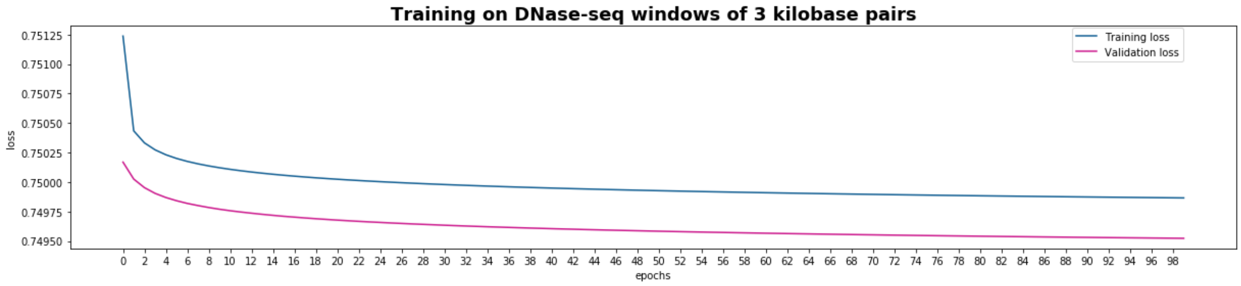


Figure S9: **Training and validation loss of the 3 kb DNase-seq CAE.** Total training time was 5 days and 13 hours.

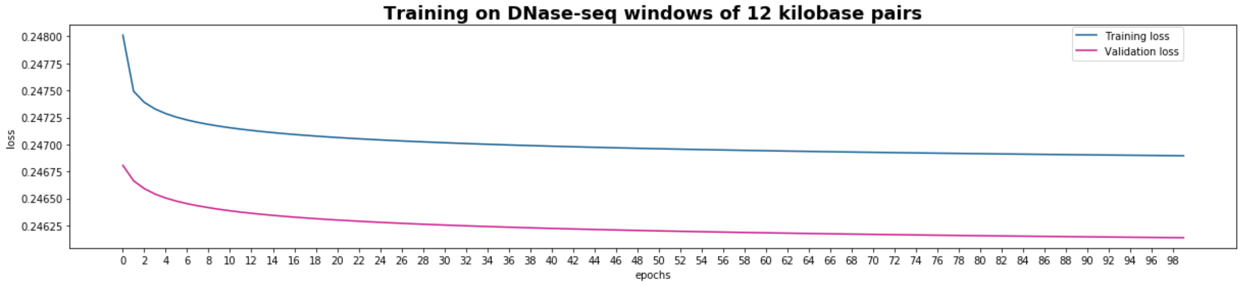


Figure S10: **Training and validation loss of the 12 kb DNase-seq CAE.** Total training time was 7 days and 15 hours.

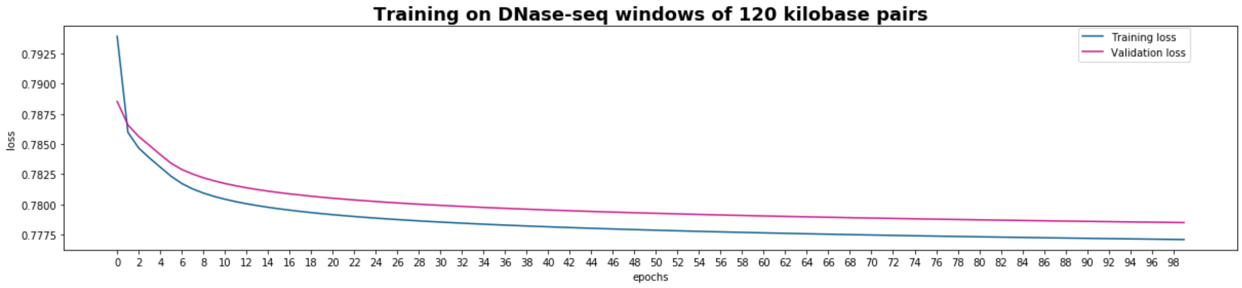


Figure S11: **Training and validation loss of the 120 kb DNase-seq CAE.** Total training time was 3 days and 5 hours.

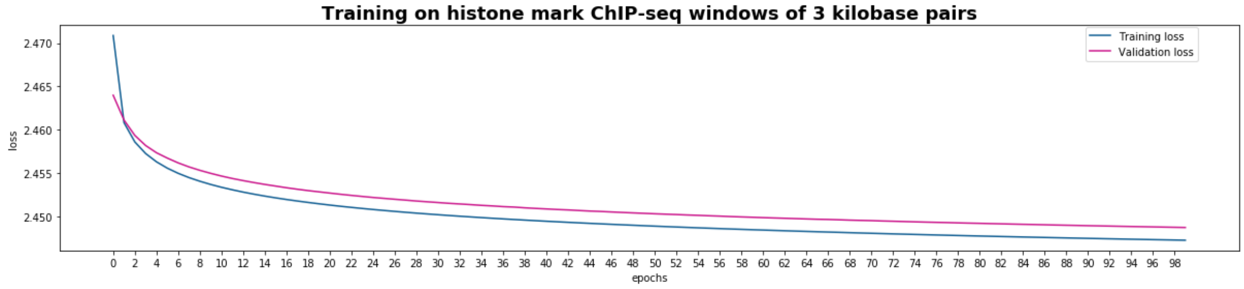


Figure S12: **Training and validation loss of the 3 kb histone mark ChIP-seq CAE.** Total training time was 4 days and 2 hours.

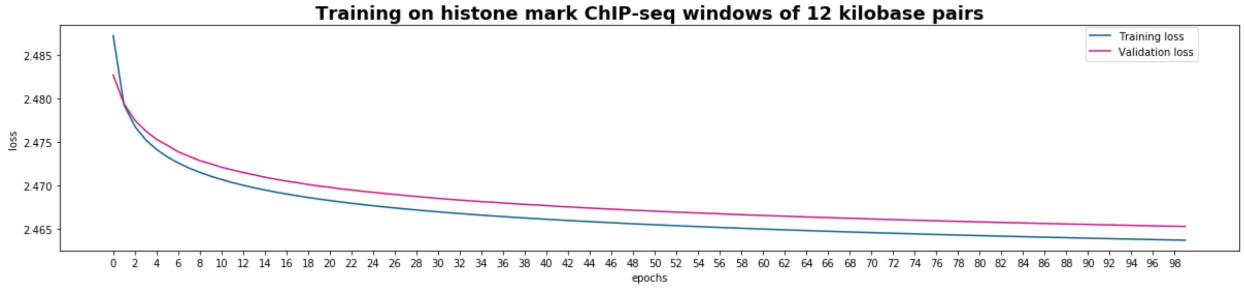


Figure S13: **Training and validation loss of the 12 kb histone mark ChIP-seq CAE.** Total training time was 5 days and 6 hours.

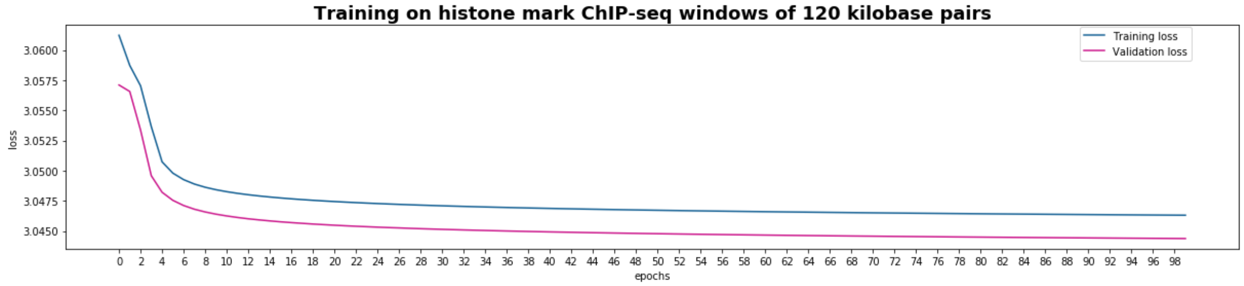


Figure S14: **Training and validation loss of the 120 kb histone mark ChIP-seq CAE.** Total training time was 1 days and 18 hours.

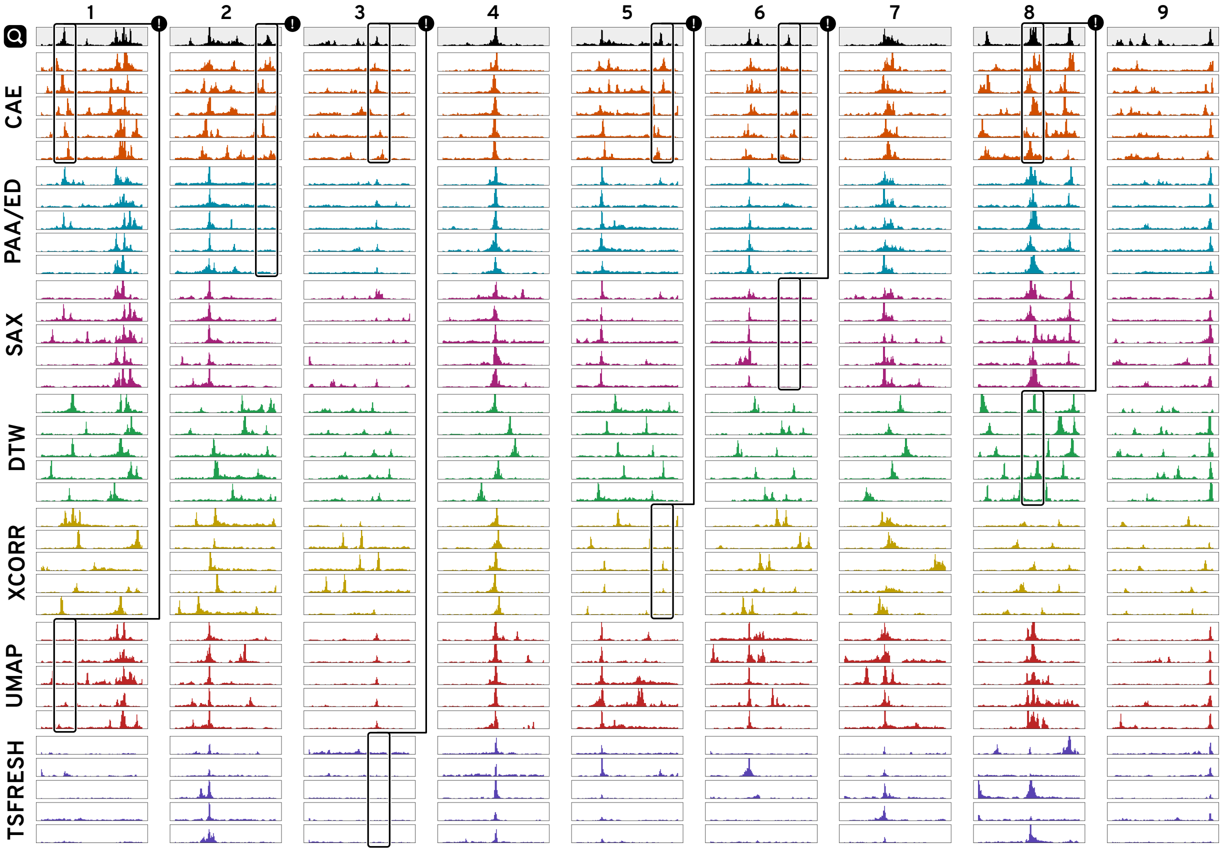


Figure S15: **Similarity Comparison of 12kb windows.**

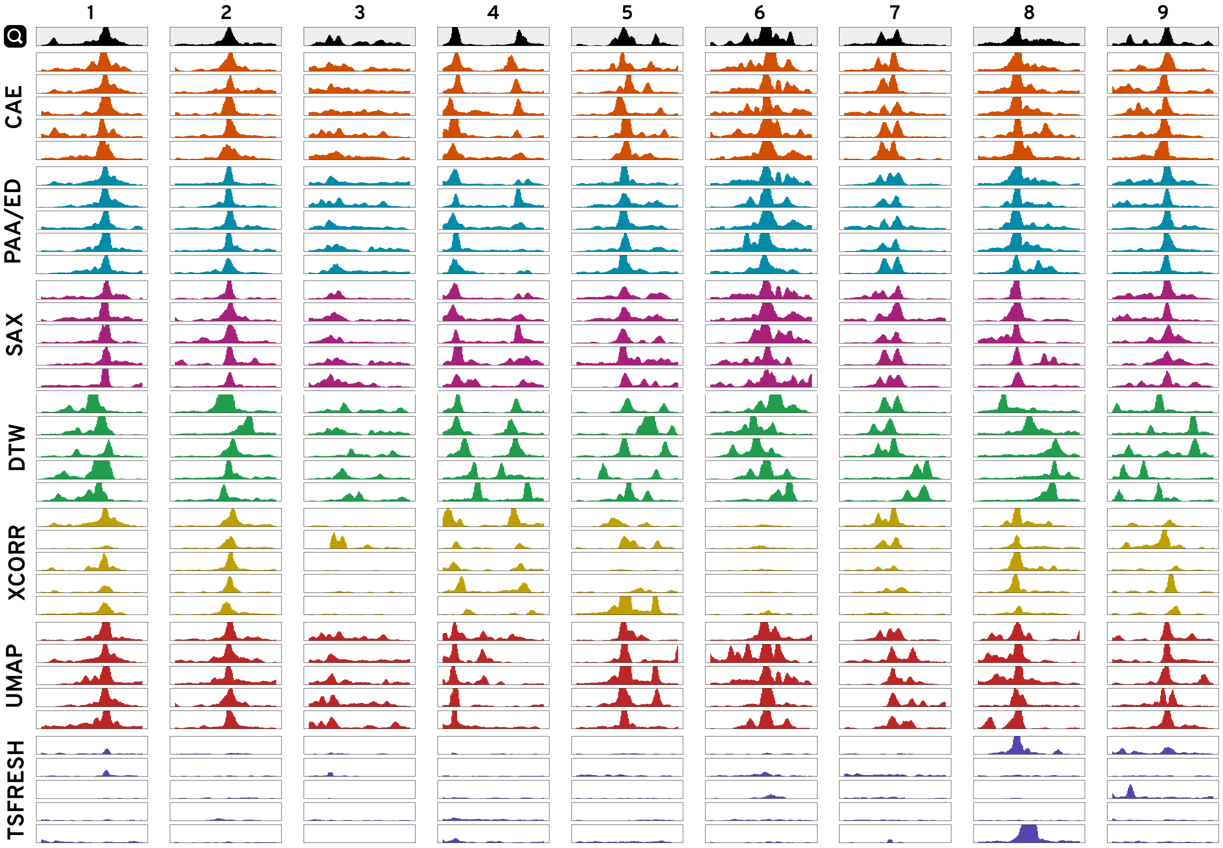


Figure S16: **Similarity Comparison of 3kb windows.**

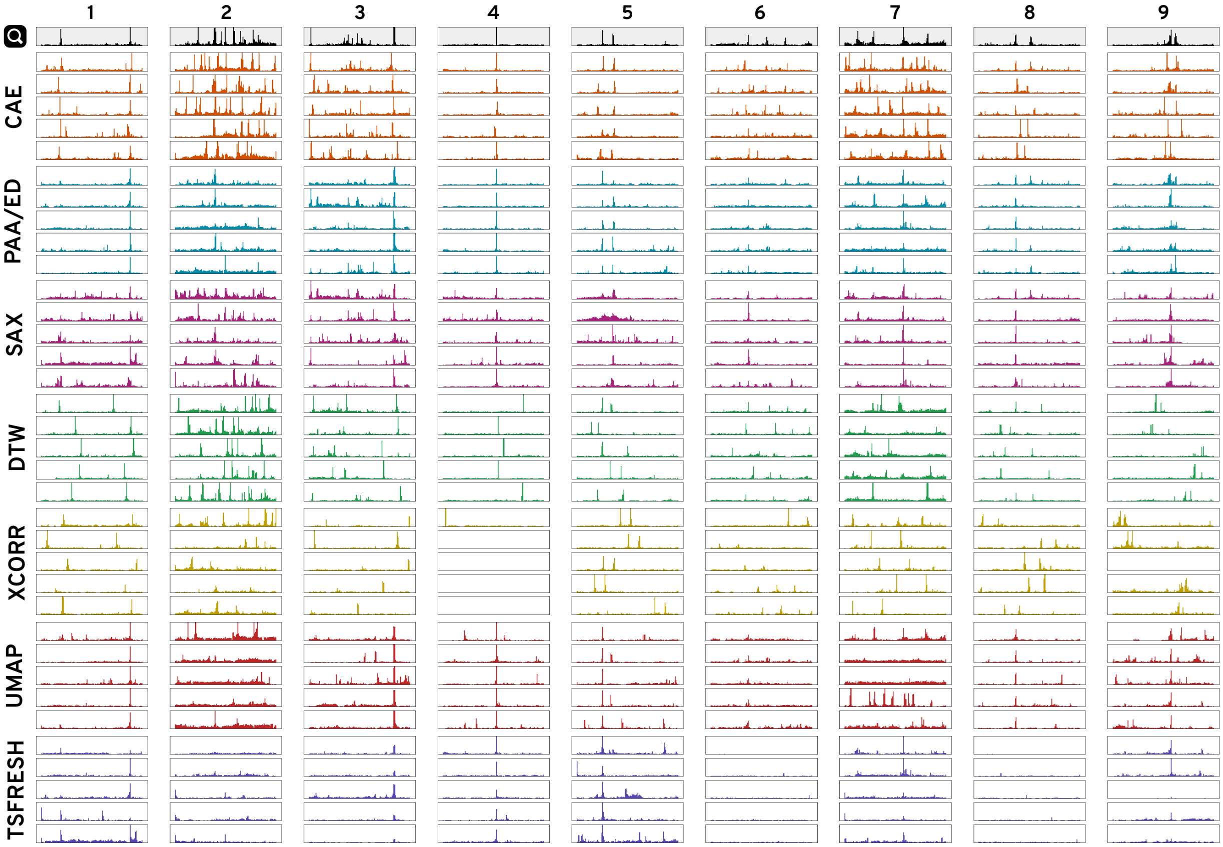


Figure S17: **Similarity Comparison of 120kb windows.**

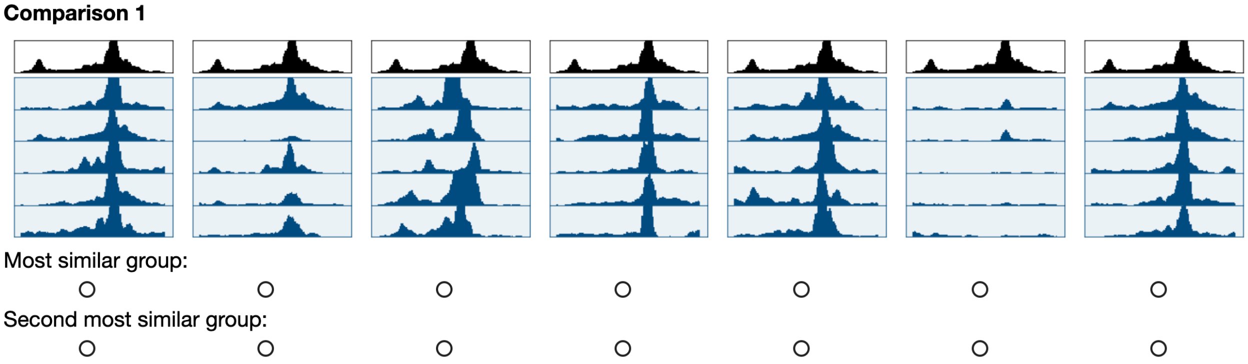


Figure S18: **User Study 1 Task.** An example task of the first user study for groupwise visual similarity comparison of the 5-nearest neighbors from all seven techniques. As a pre-study showed that comparisons can be hard when there are multiple group with good matches we ask the Amazon Mechanical Turkers to select the most and second most similar group of patterns.

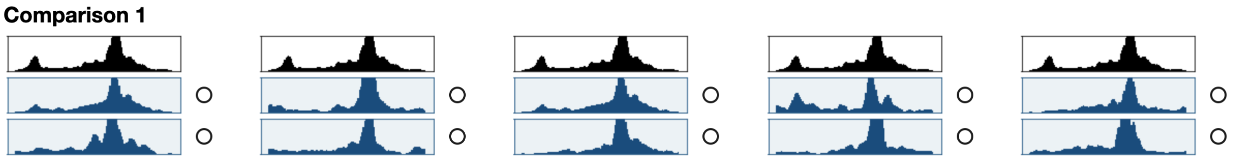


Figure S19: **User Study 2 Task.** An example task of the second user study for pairwise visual similarity comparison of the 5-nearest neighbors from our technique and Euclidean distance (ED).

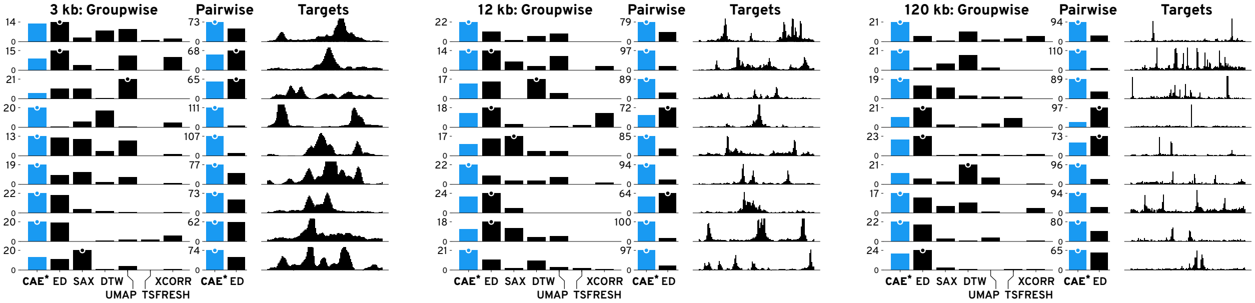


Figure S20: **Results of the Similarity Comparison.** Results for the subjective similarity comparison are shown as votes per technique per target pattern per pattern size. More votes mean that participants regarded the results to be visually more similar to the target. The highest votes are indicated by a dot. Results for our method (CAE) are drawn in blue.

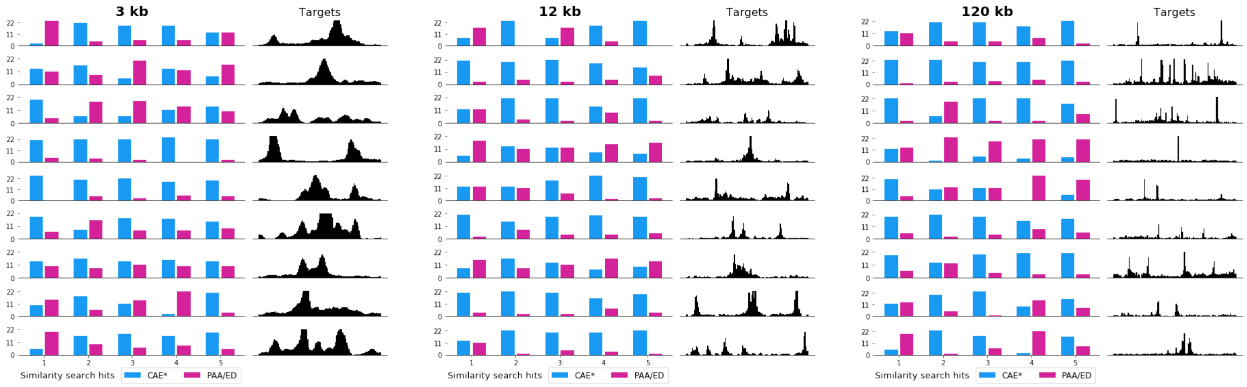


Figure S21: **User Study 2 Results per Nearest Neighbor.** Results of the second user study for all the five pairwise comparisons of the nearest neighbors between our model (CAE) and Euclidean distance (ED).

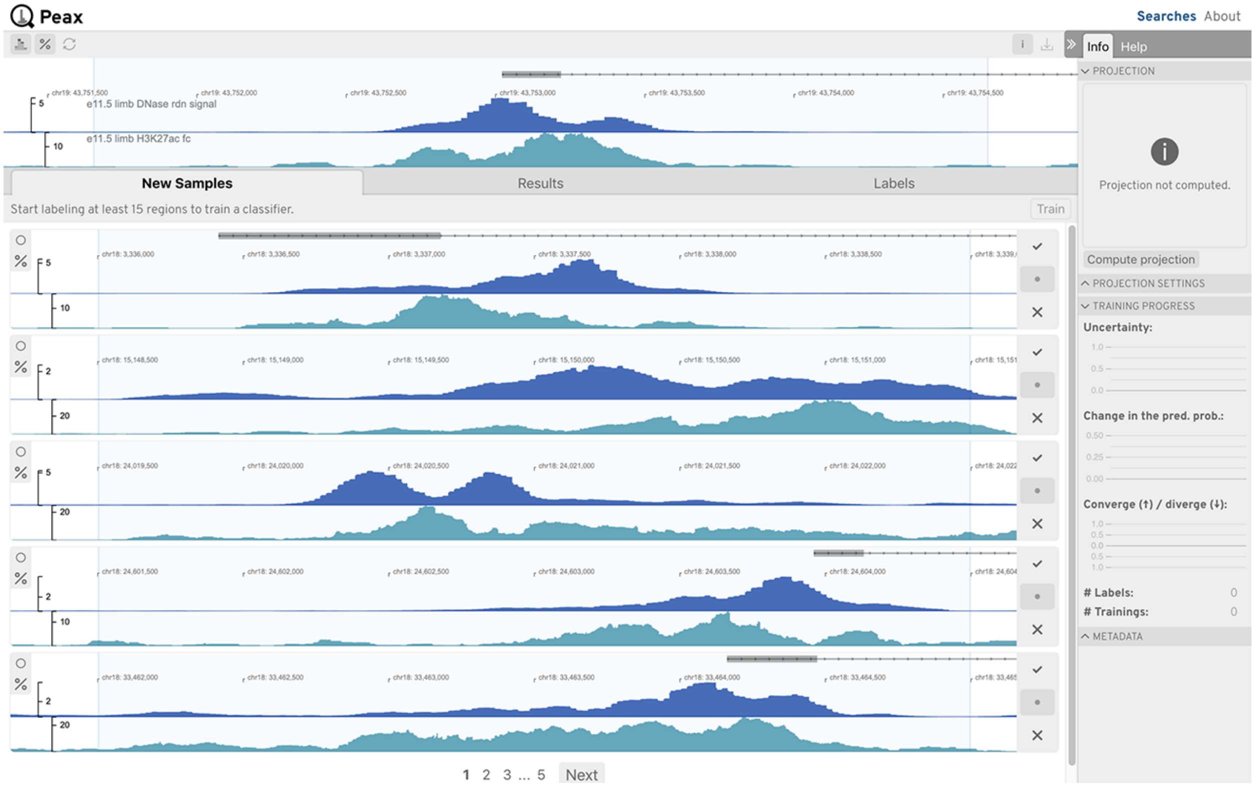


Figure S22: **Use Case 1: Asymmetrical Peak: Step 1.** Start search.

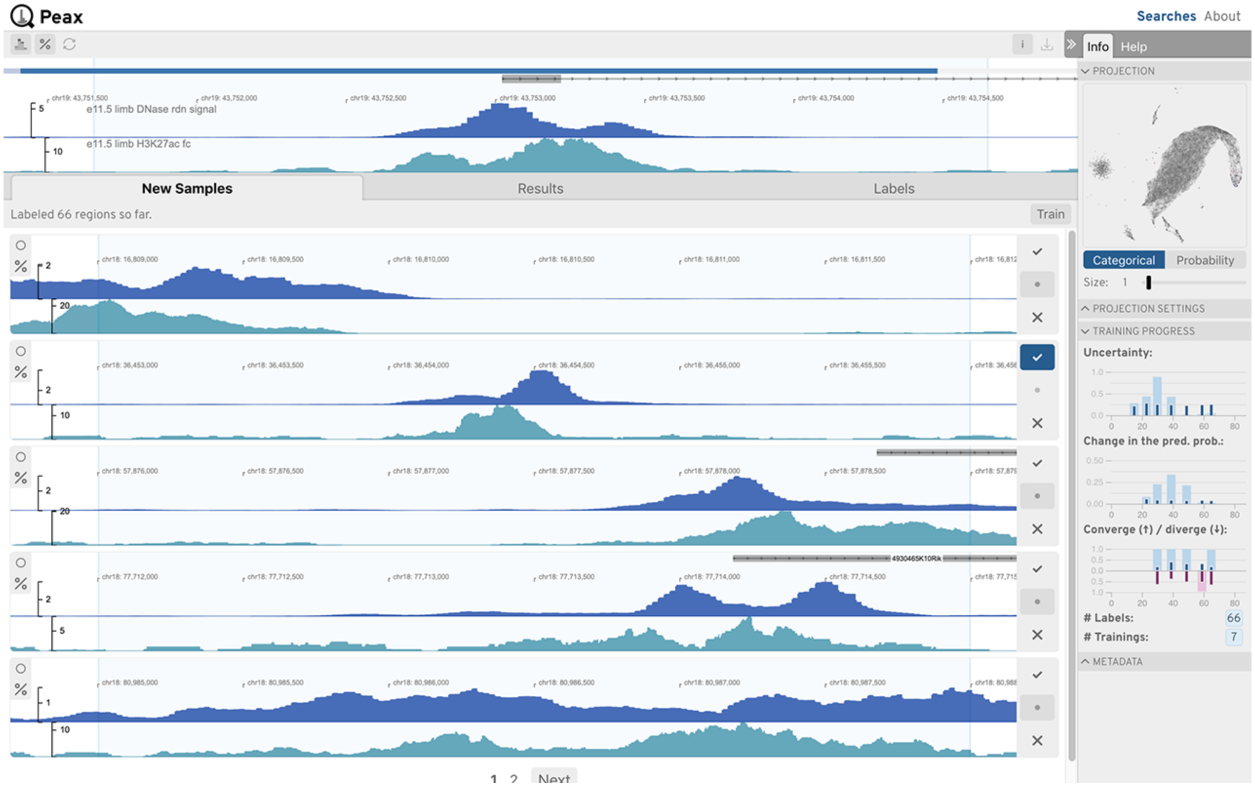


Figure S23: **Use Case 1: Asymmetrical Peak: Step 2.** Initial labeling is done. Inspecting the embedding view and the progress.

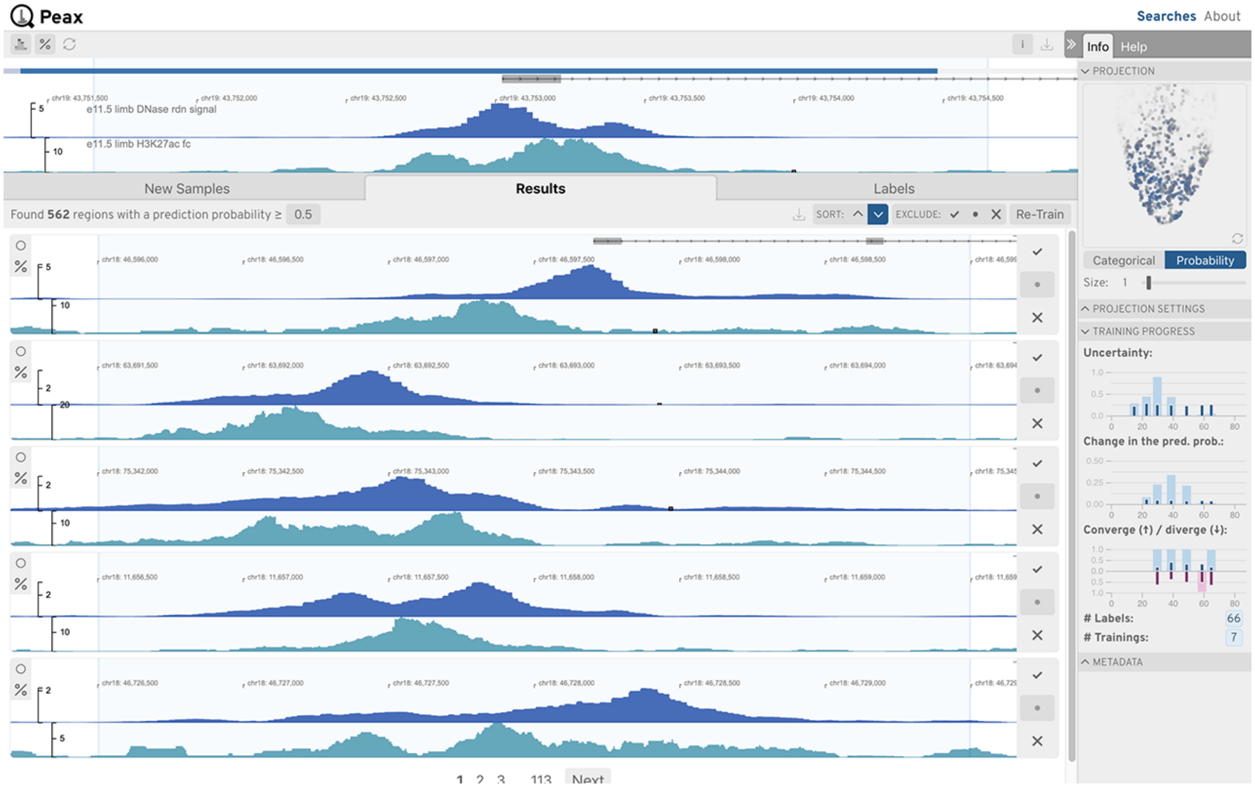


Figure S24: **Use Case 1: Asymmetrical Peak: Step 3.** Inspecting initial results.

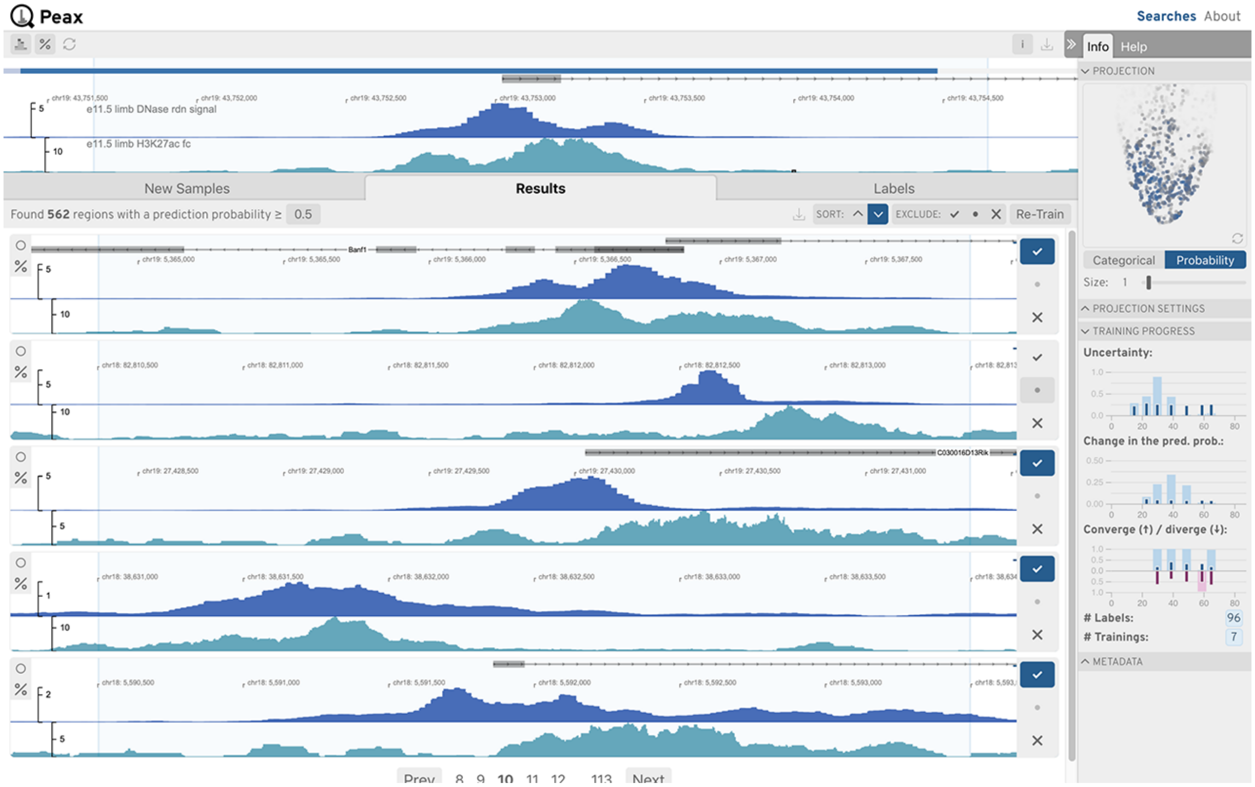


Figure S25: **Use Case 1: Asymmetrical Peak: Step 4.** Going through results, boosting true positives, and cleaning false positives.

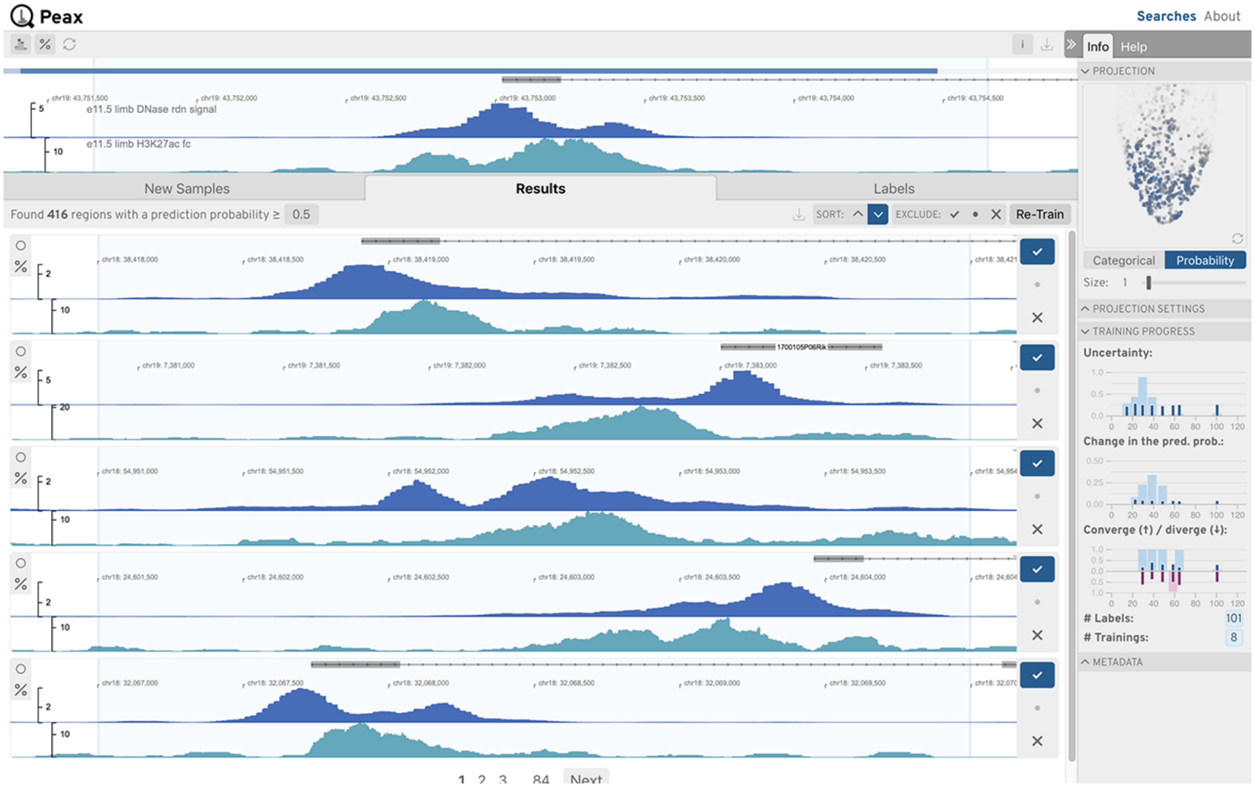


Figure S26: **Use Case 1: Asymmetrical Peak: Step 5.** Inspecting results after the retraining the classifier.

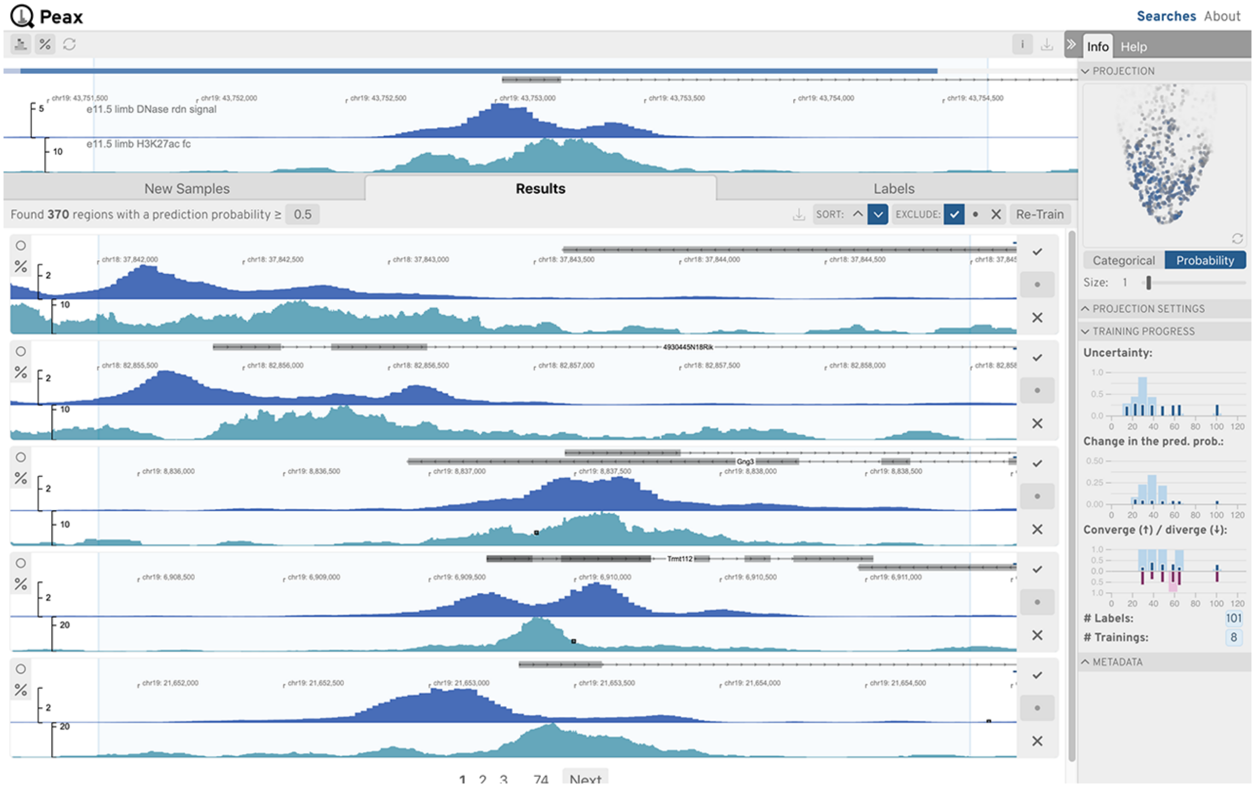


Figure S27: **Use Case 1: Asymmetrical Peak: Step 6.** Inspecting unlabeled results.

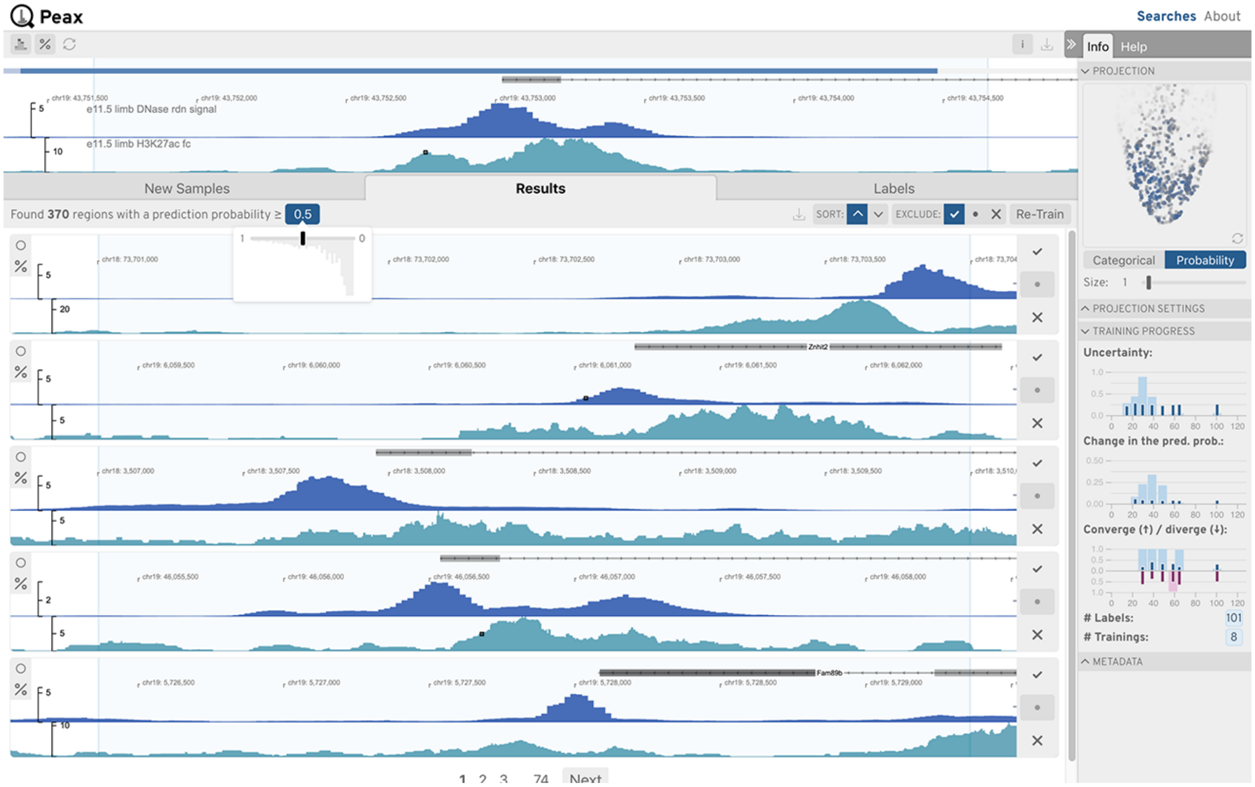


Figure S28: **Use Case 1: Asymmetrical Peak: Step 7.** Adjusting the prediction probability while looking at the boundary.

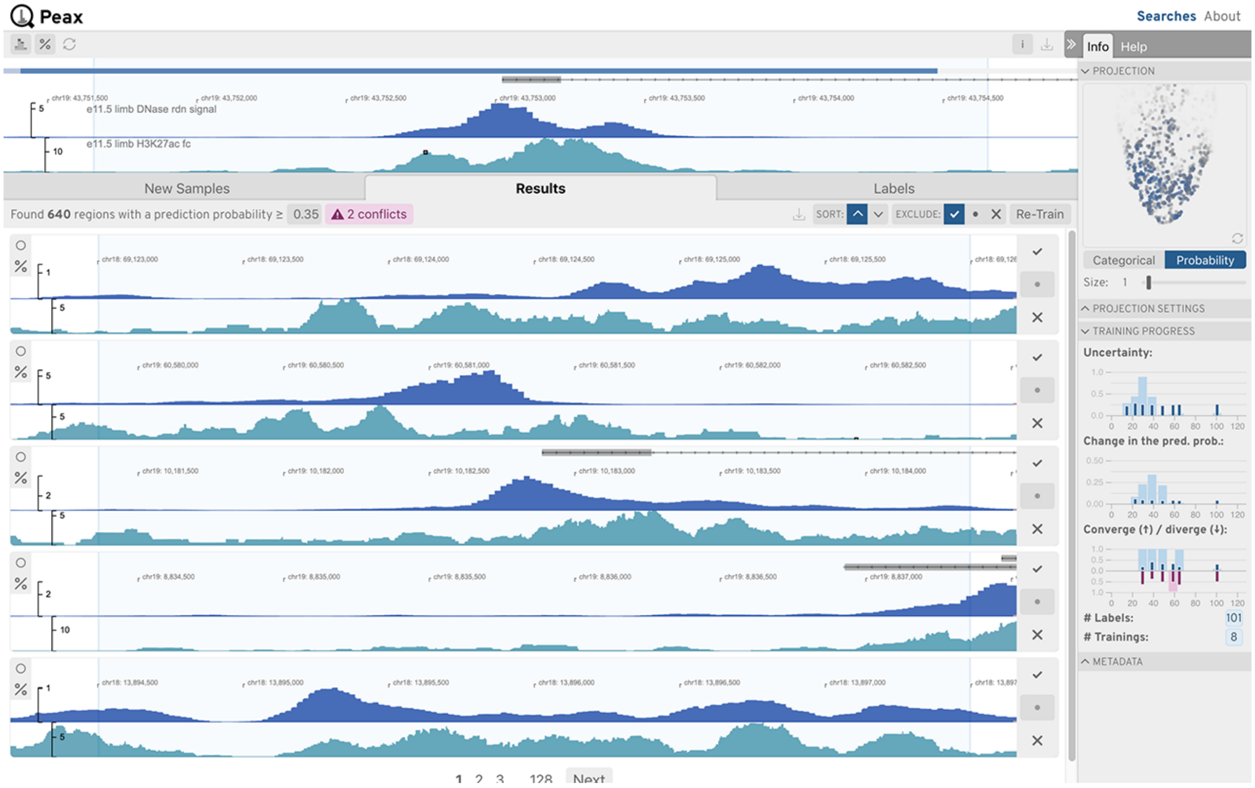


Figure S29: **Use Case 1: Asymmetrical Peak: Step 8.** Identify that conflicts exist at the new prediction probability threshold.

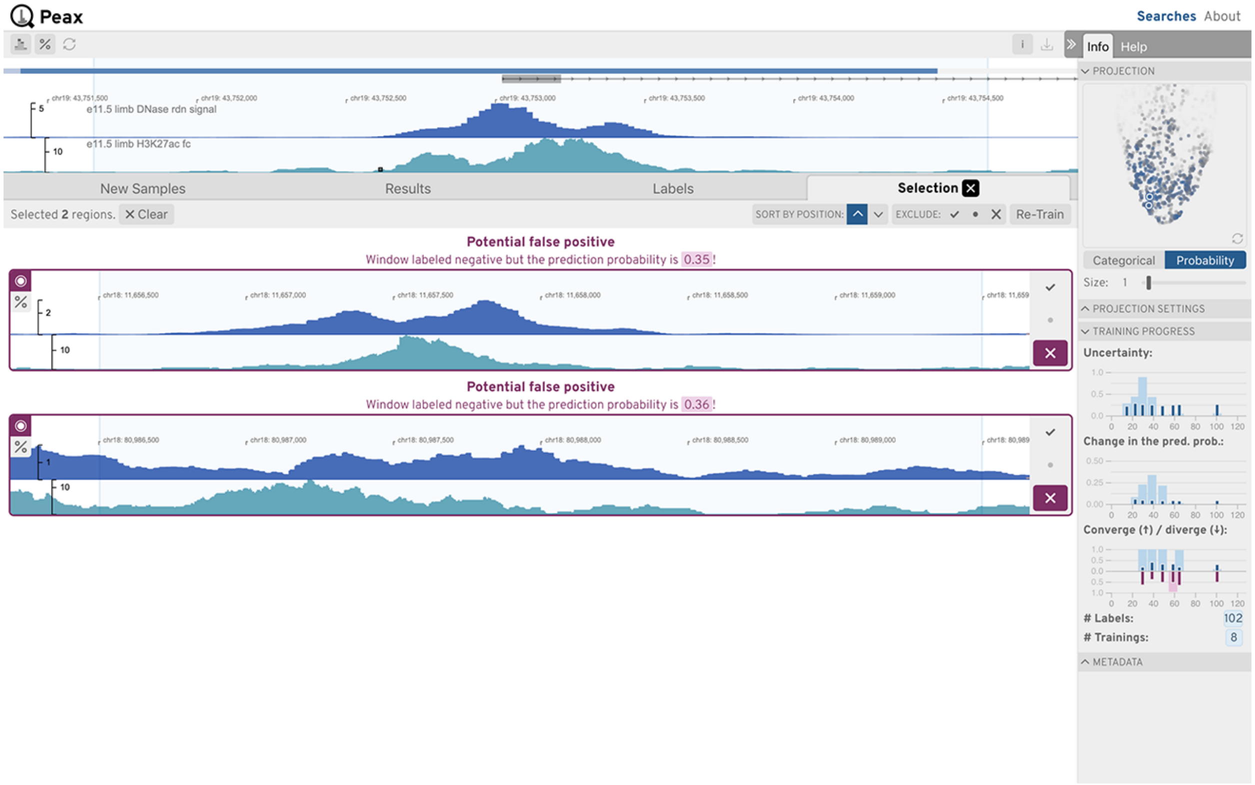


Figure S30: **Use Case 1: Asymmetrical Peak: Step 9.** Viewing conflicting windows.

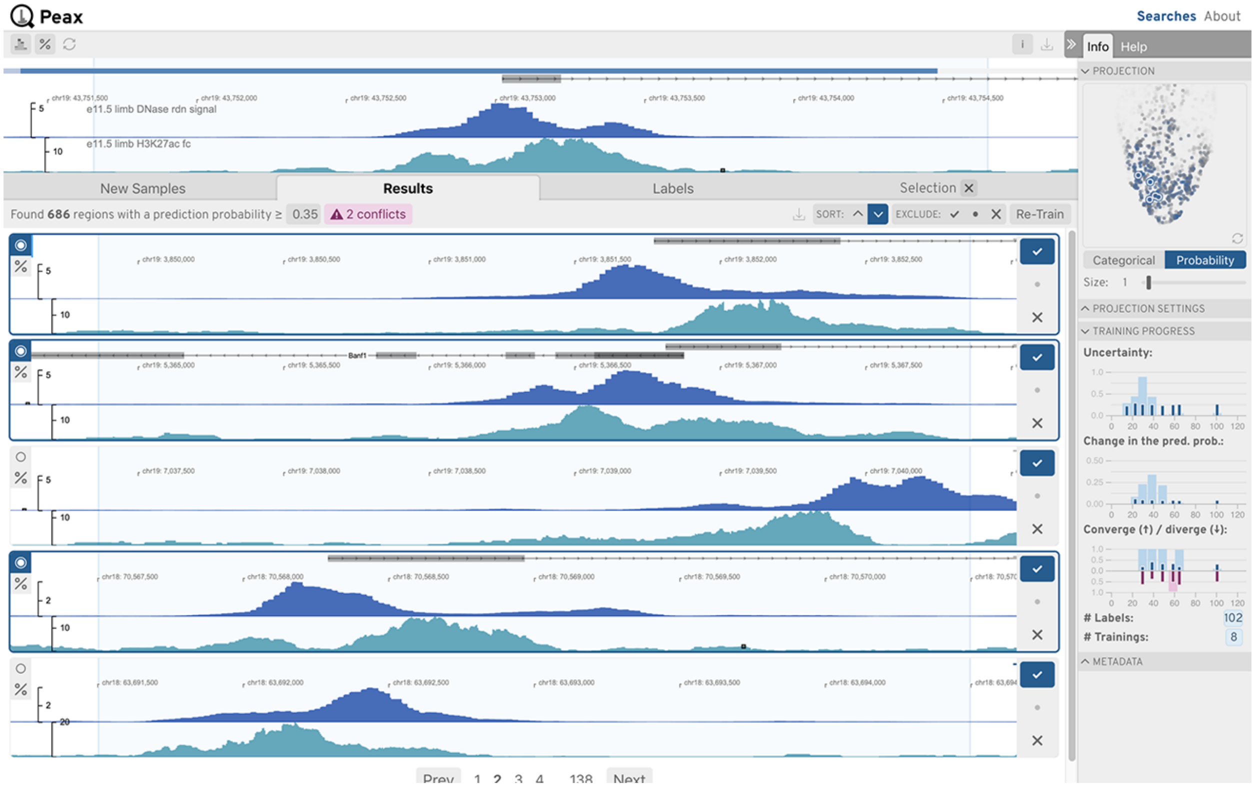


Figure S31: **Use Case 1: Asymmetrical Peak: Step 10.** Improving results using selected windows in a certain local neighborhood.

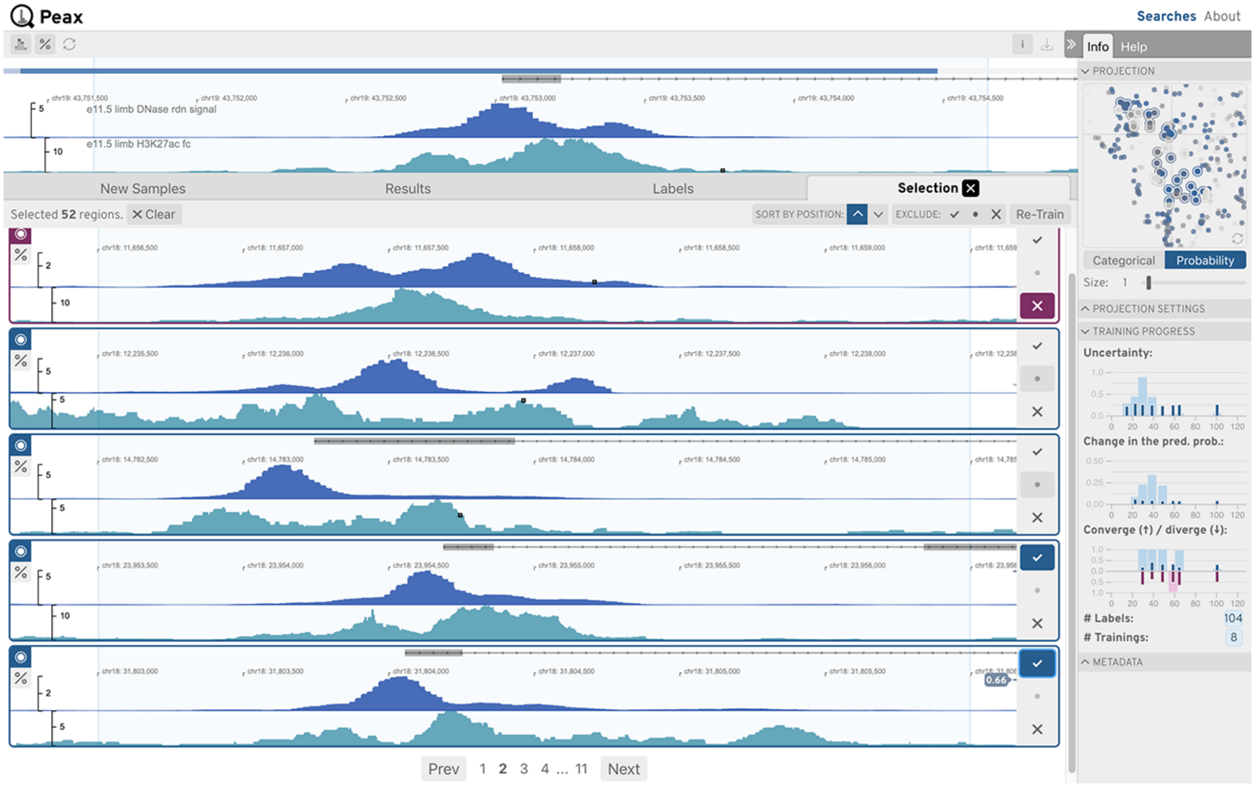


Figure S32: **Use Case 1: Asymmetrical Peak: Step 11.** Improving results further using selected windows in a certain local neighborhood.

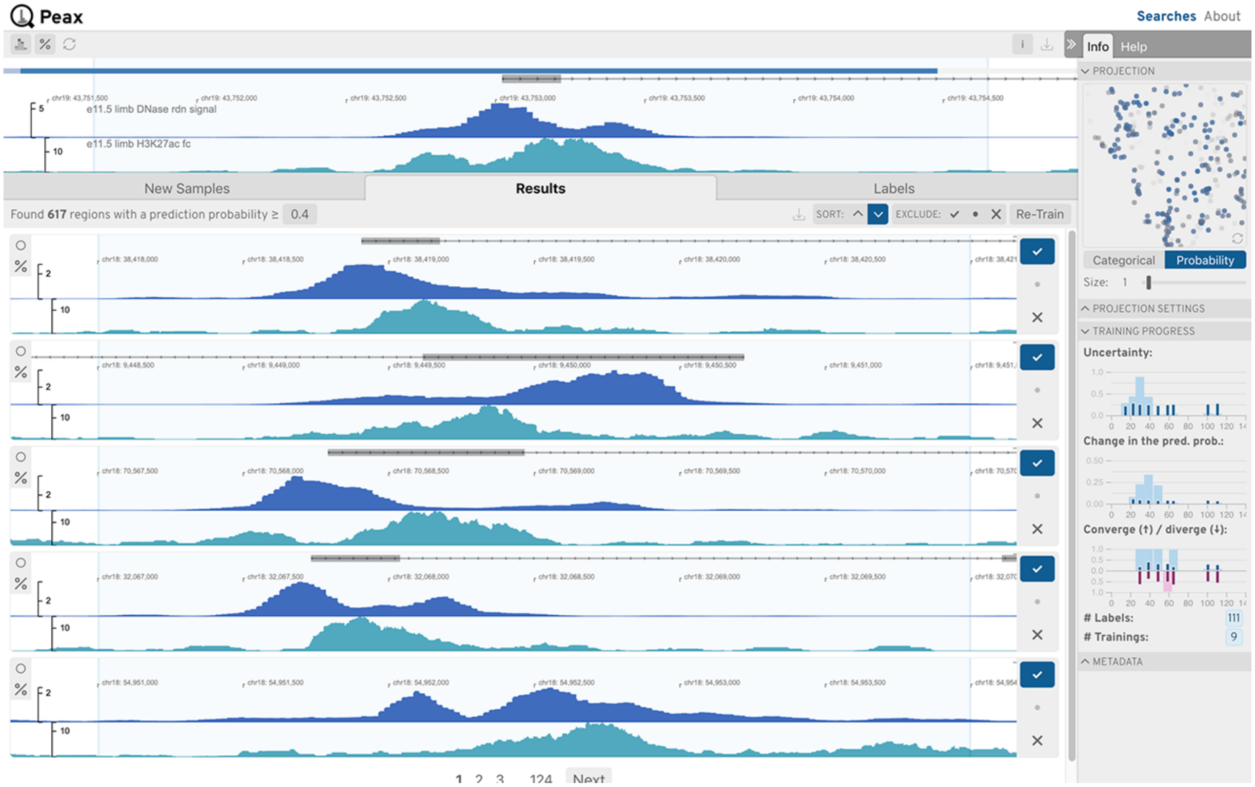


Figure S33: **Use Case 1: Asymmetrical Peak: Step 12.** Final result.

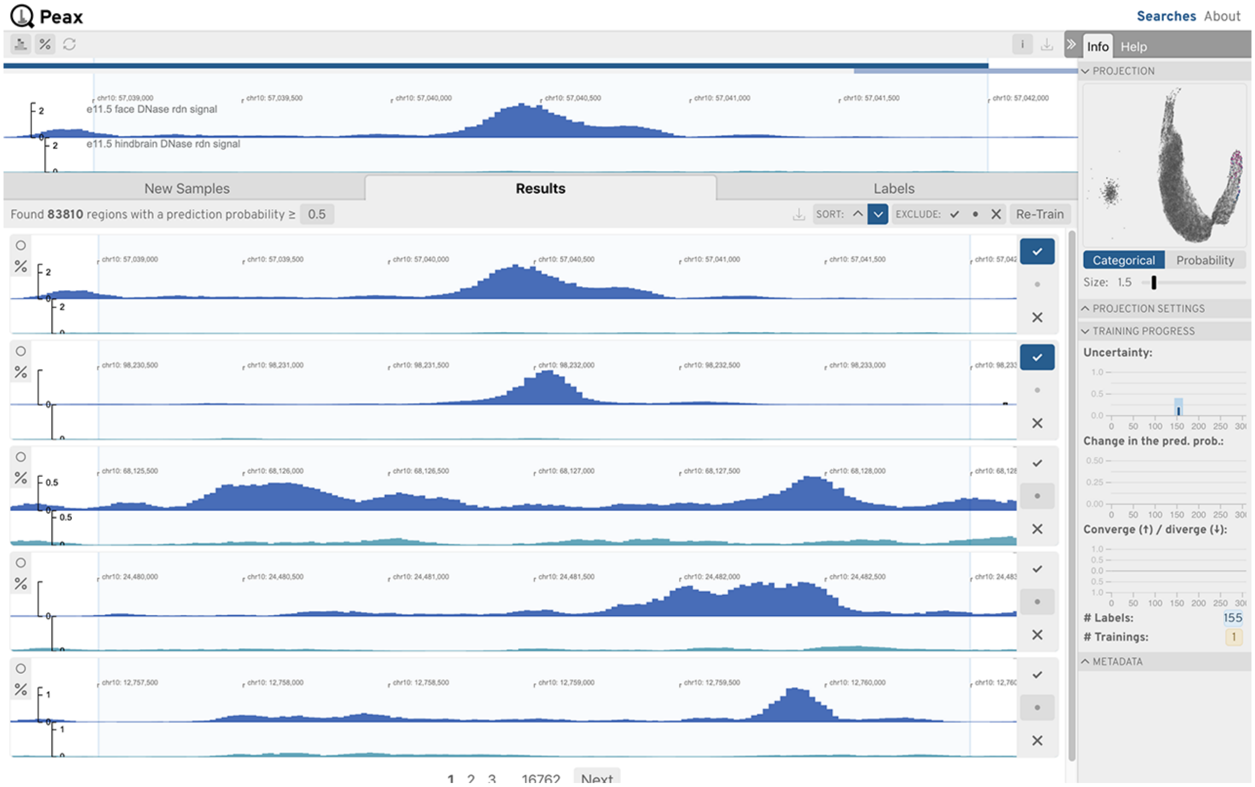


Figure S34: **Use Case 2: Differential Peak: Step 1.** Initial classifier after loading the labels.

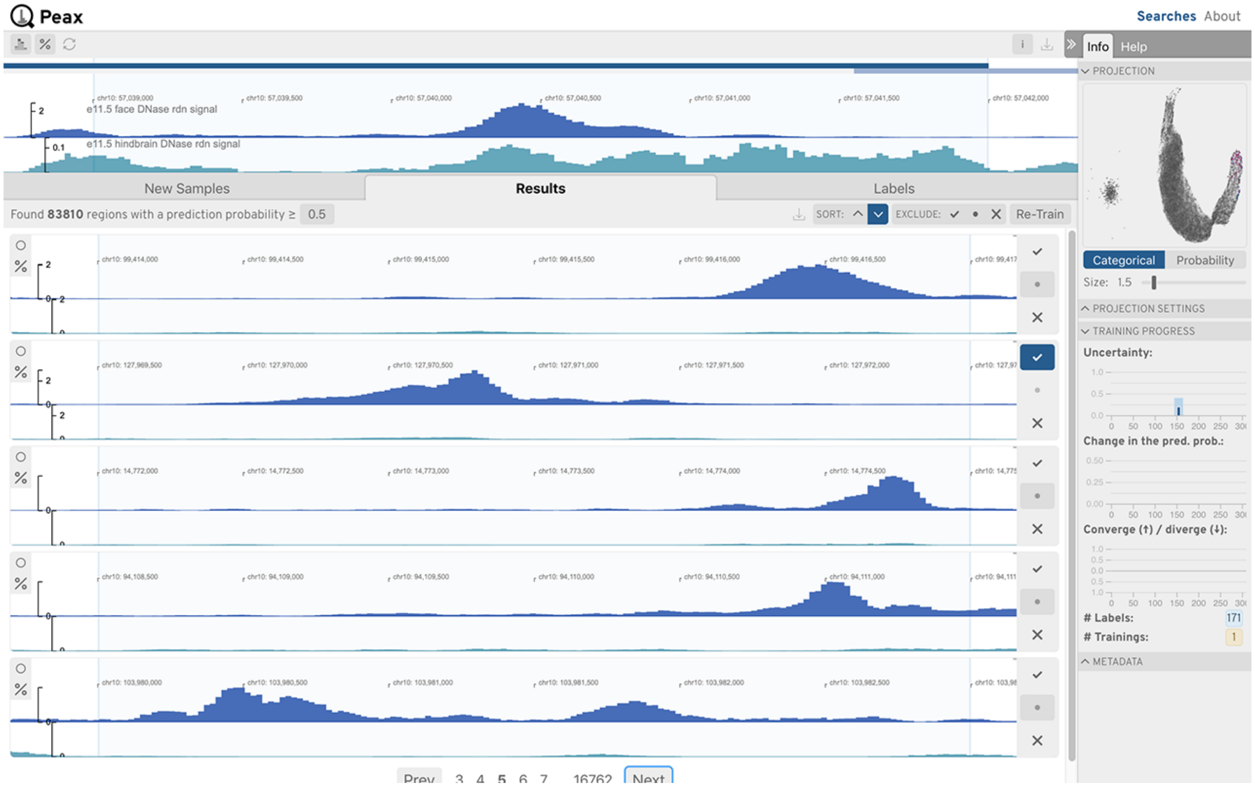


Figure S35: **Use Case 2: Differential Peak: Step 2.** Inspect top results.

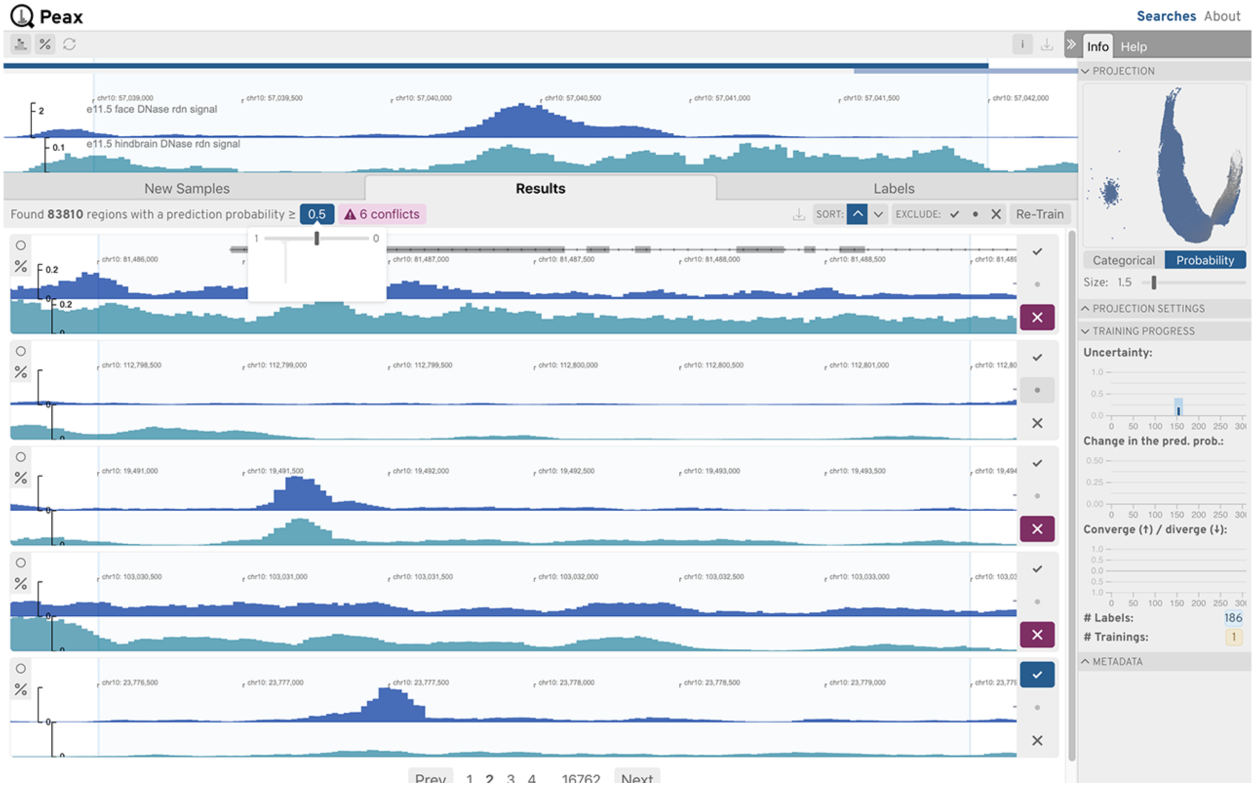


Figure S36: **Use Case 2: Differential Peak: Step 3.** Assess recall, identify high uncertainty, and add negative labels.

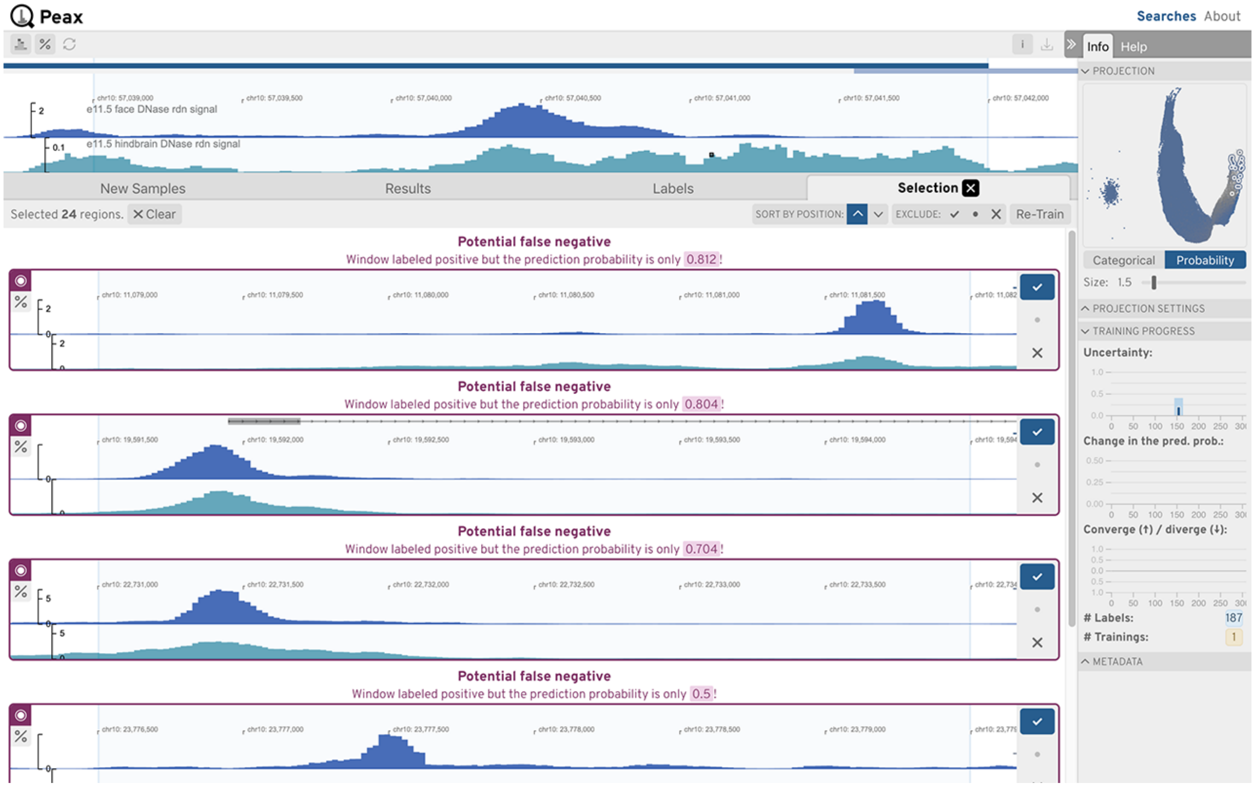


Figure S37: **Use Case 2: Differential Peak: Step 4.** Resolve conflicting labels from the peak annotations that defined the pre-loaded labels.

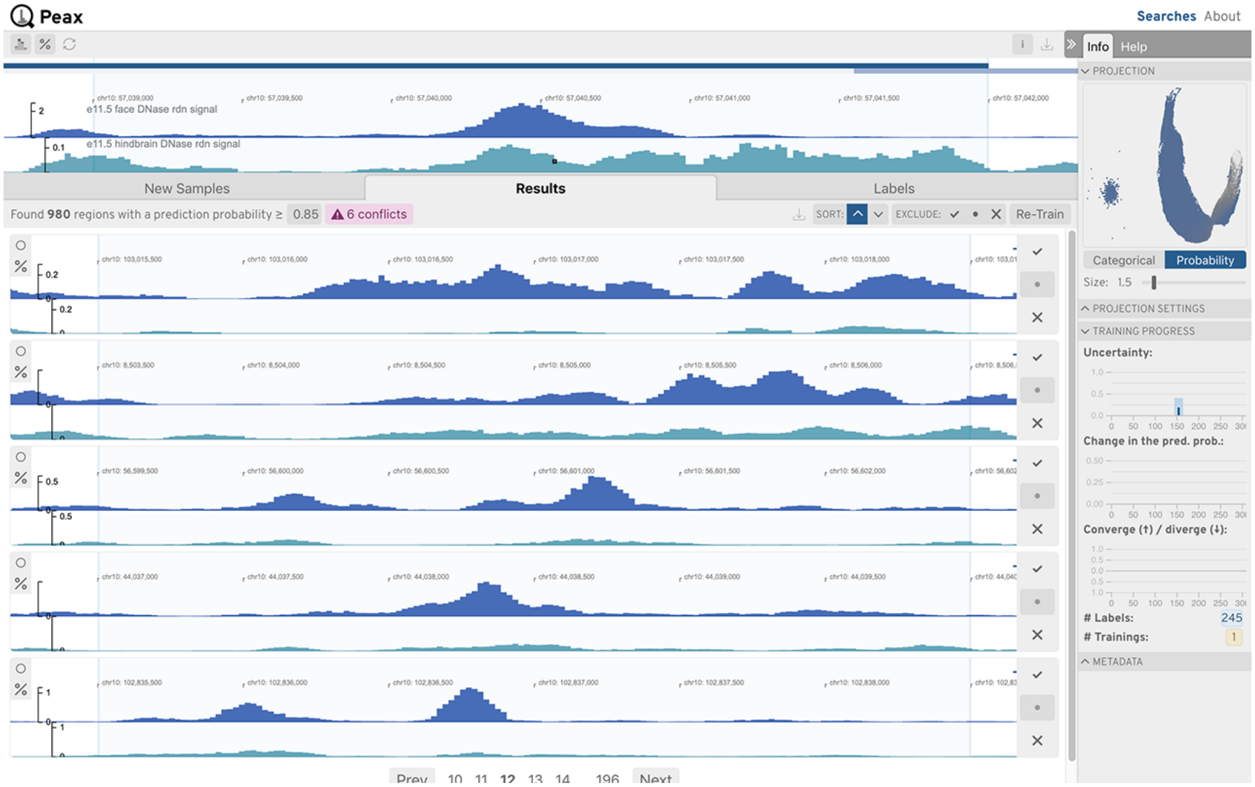


Figure S38: **Use Case 2: Differential Peak: Step 5.** Instances of unclear differential peak. While the lower DNase-seq contains much less signal peaks still overlap.

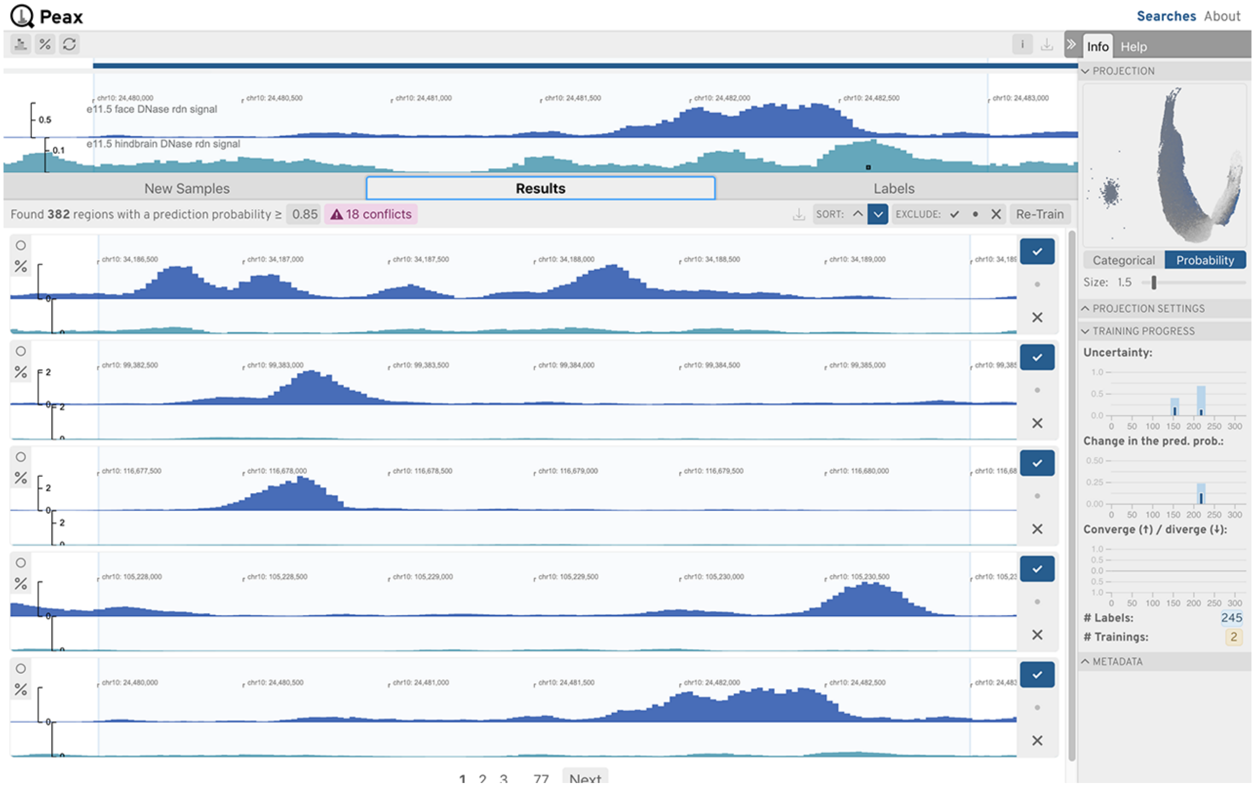


Figure S39: **Use Case 2: Differential Peak: Step 6.** Initial classifier after loading the labels.

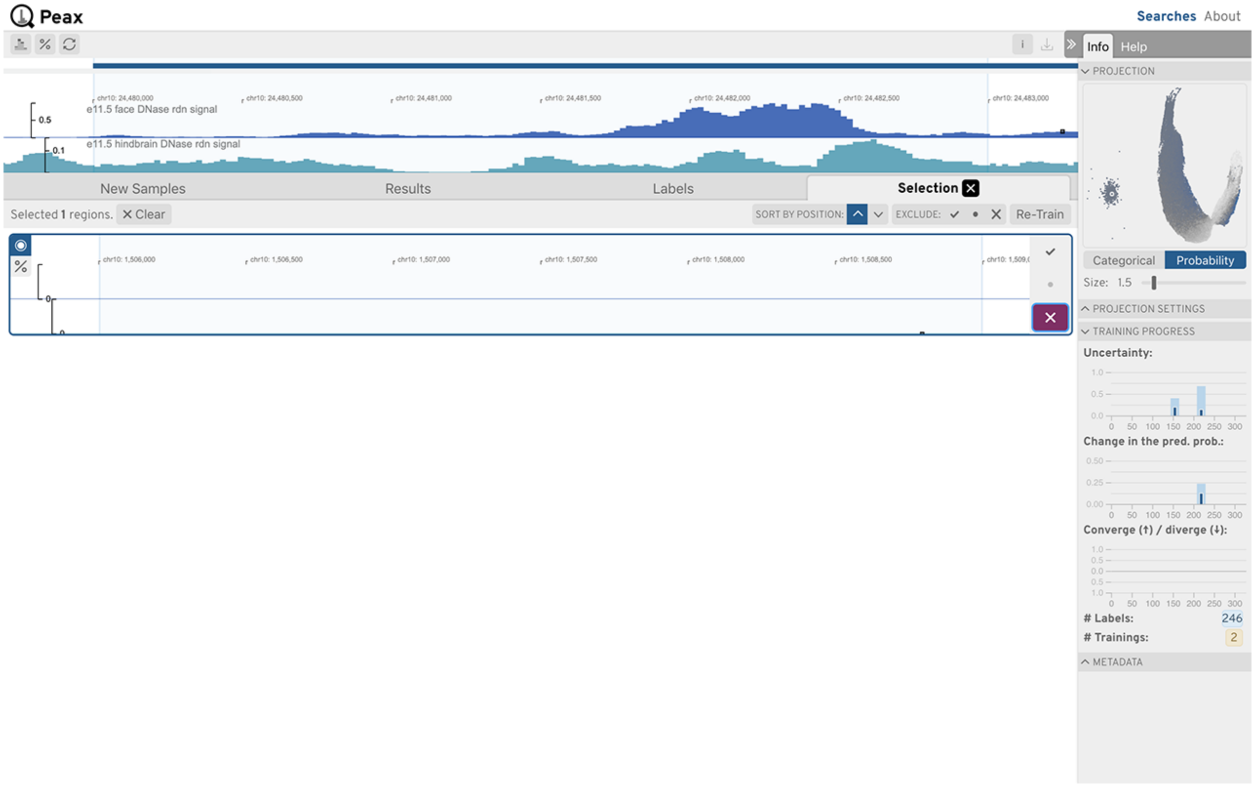


Figure S40: **Use Case 2: Differential Peak: Step 7.** After cleaning labels and retraining the uncertainty of borderline windows decreased substantially. See embedding view.

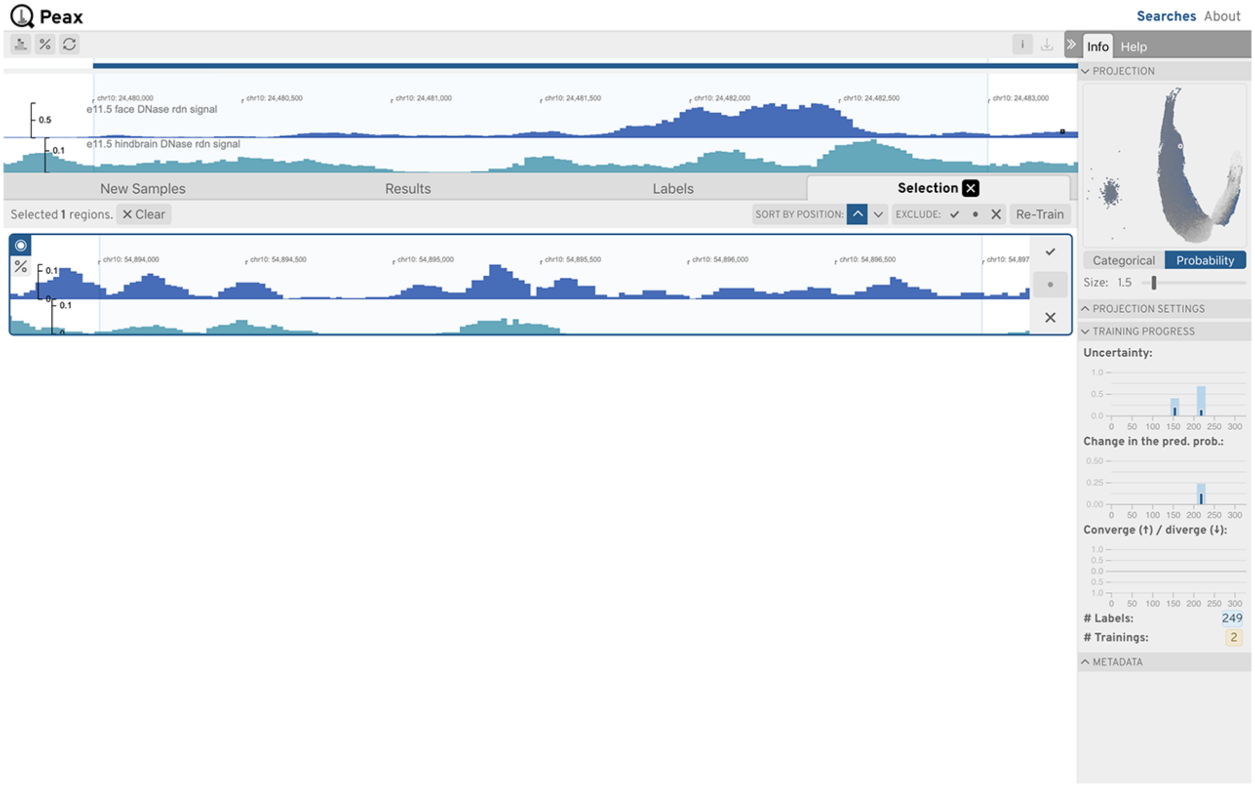


Figure S41: **Use Case 2: Differential Peak: Step 8.** Inspecting local uncertain neighborhoods with the embedding view.

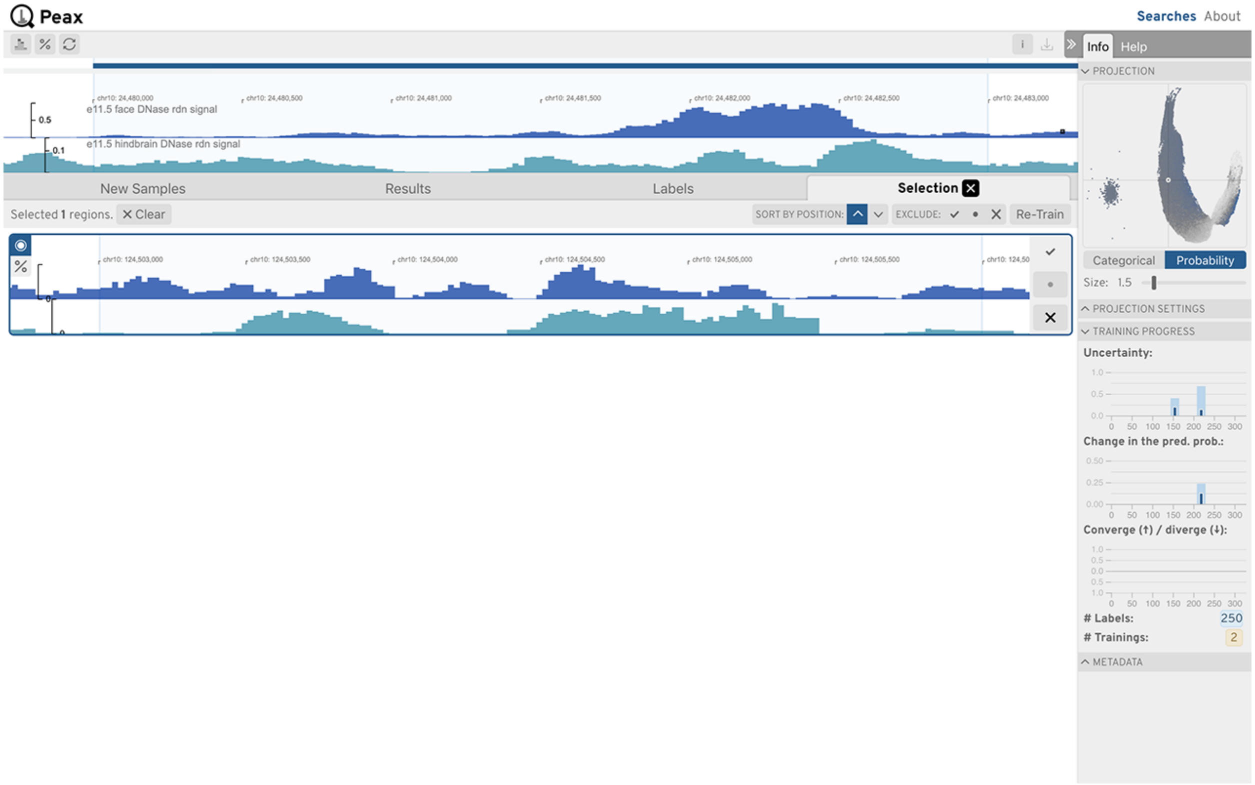


Figure S42: **Use Case 2: Differential Peak: Step 9.** Inspecting local uncertain neighborhoods with the embedding view.

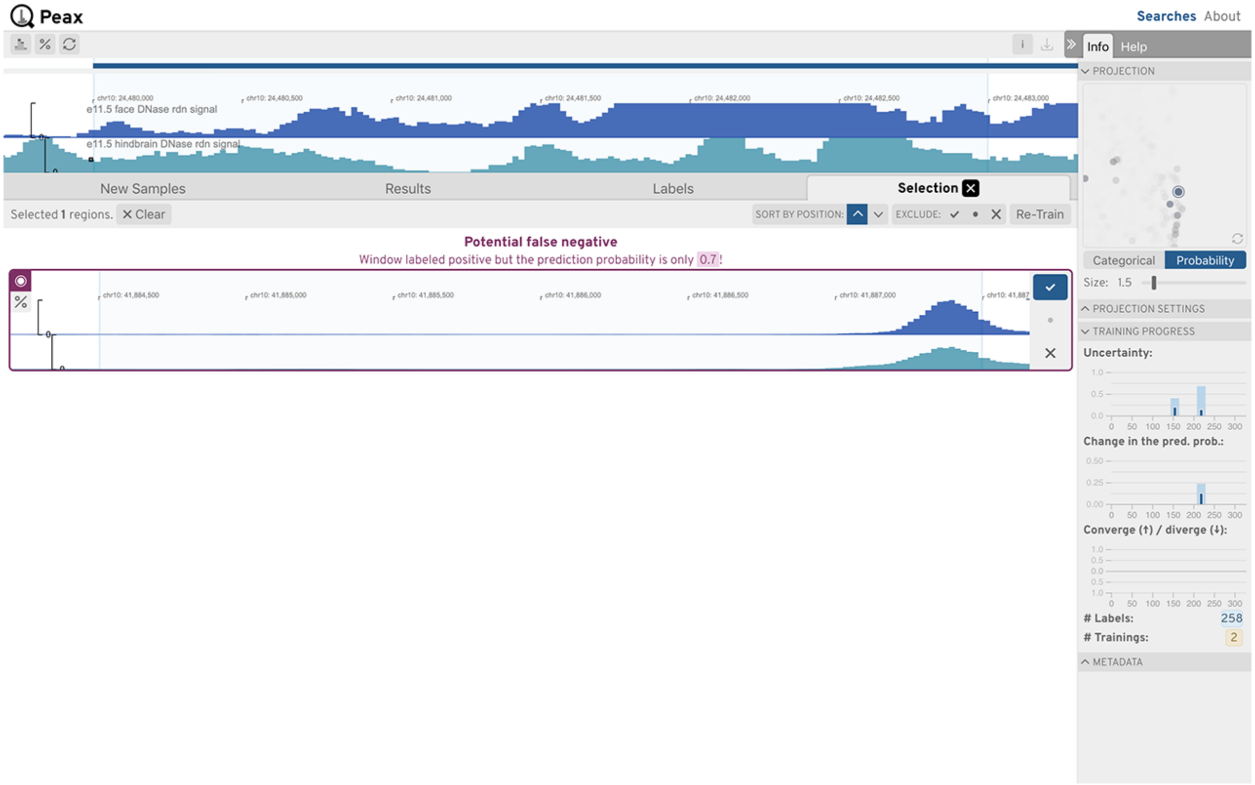


Figure S43: **Use Case 2: Differential Peak: Step 10.** Inspecting local uncertain neighborhoods with the embedding view.

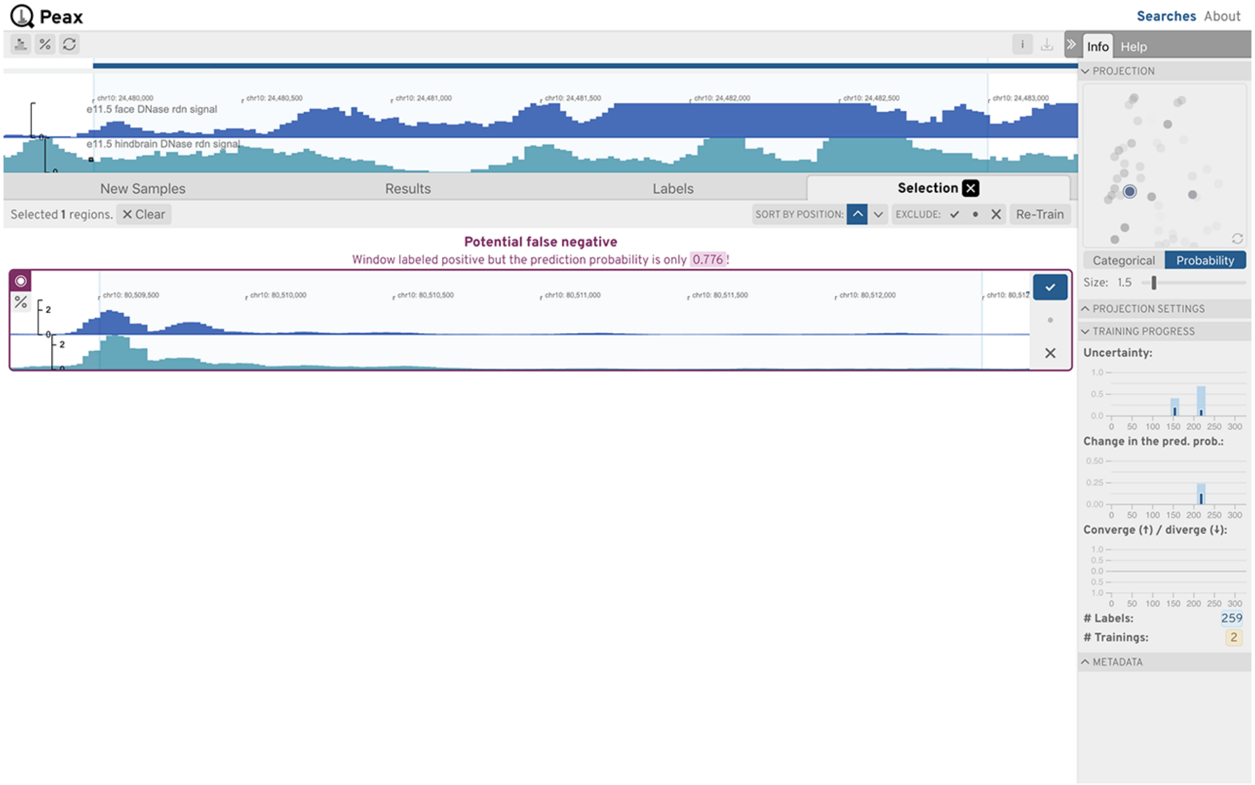


Figure S44: **Use Case 2: Differential Peak: Step 11.** Inspecting local uncertain neighborhoods with the embedding view.

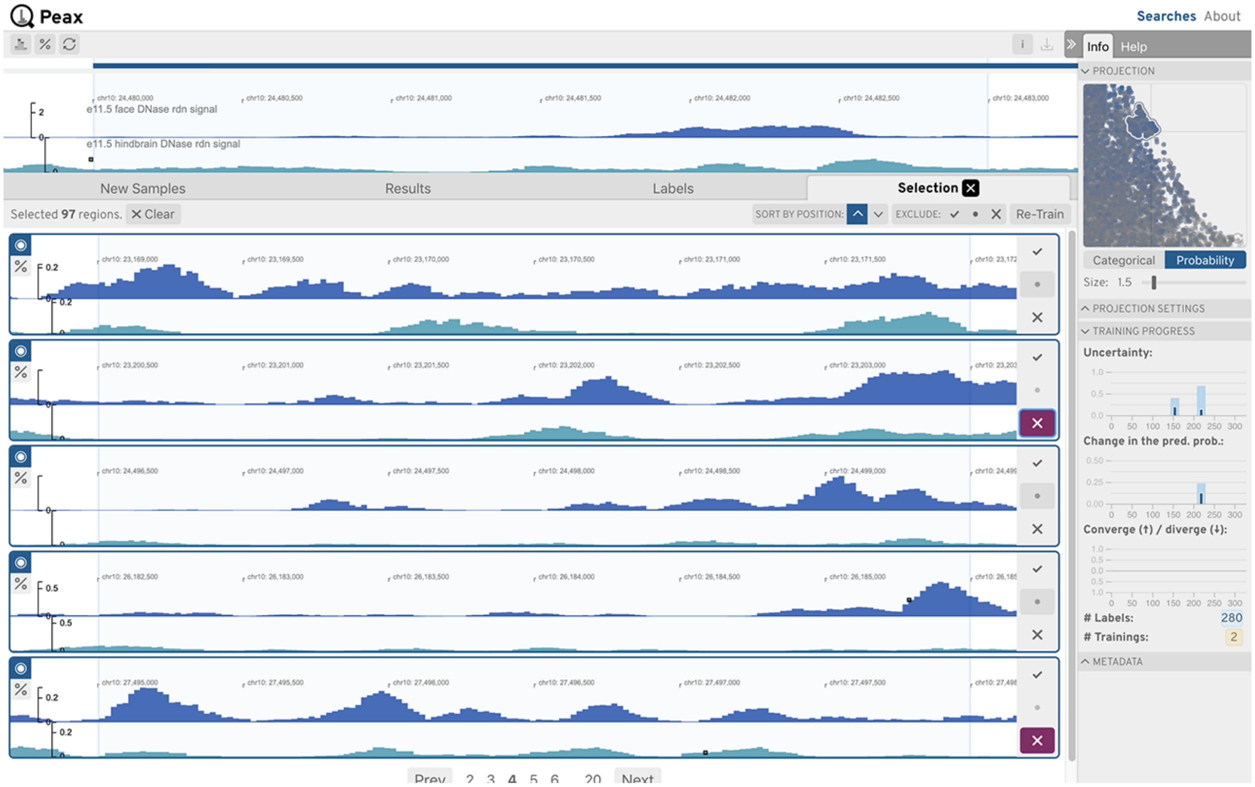


Figure S45: **Use Case 2: Differential Peak: Step 12.** Inspecting local uncertain neighborhoods with the embedding view.

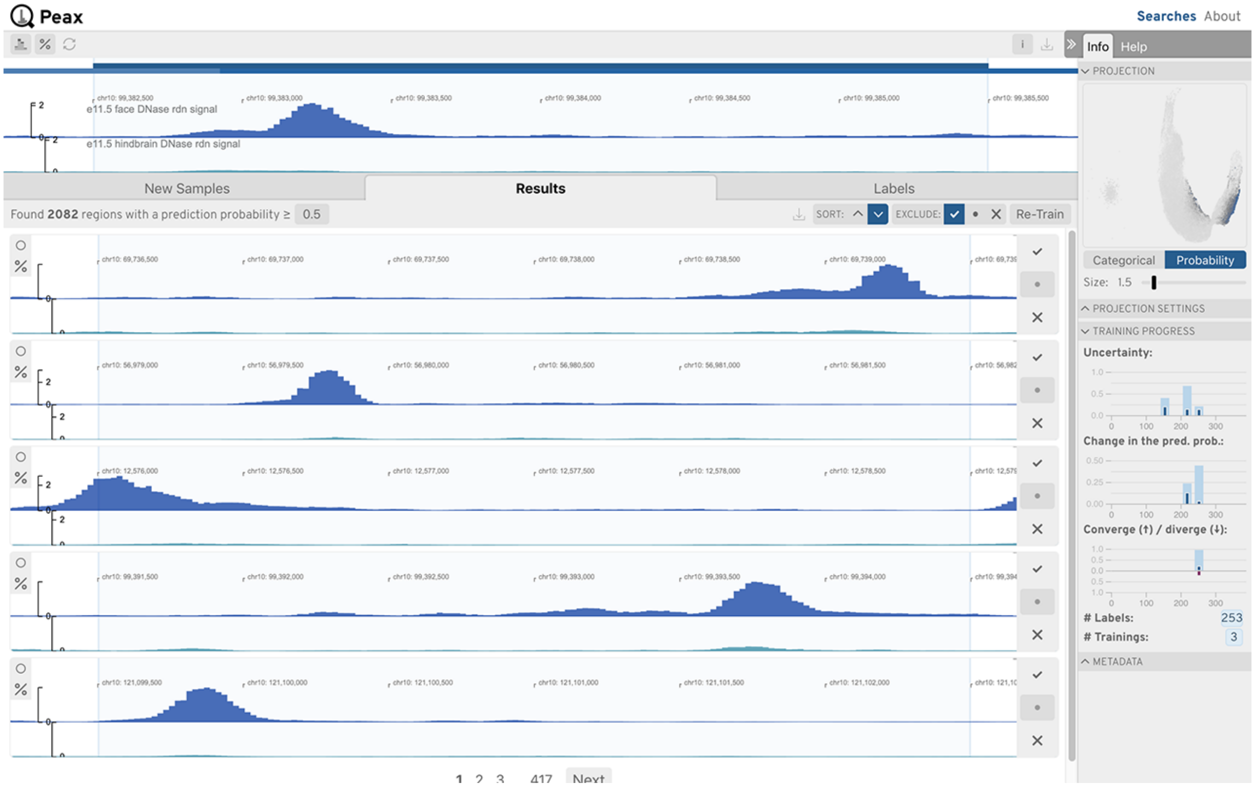


Figure S46: **Use Case 2: Differential Peak: Step 13.** Final results excluding the already positively labeled windows.

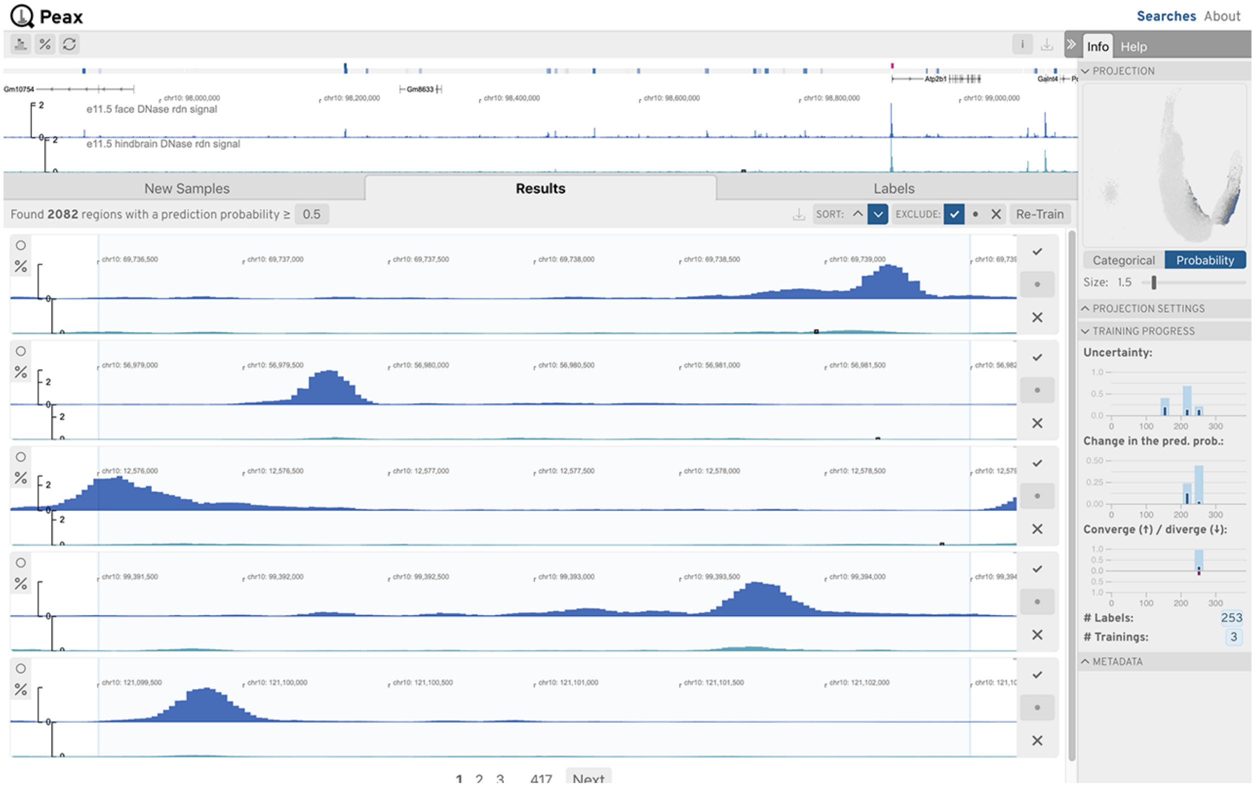


Figure S47: **Use Case 2: Differential Peak: Step 14.** Zooming out in the query view highlights many areas with potential differential peaks.

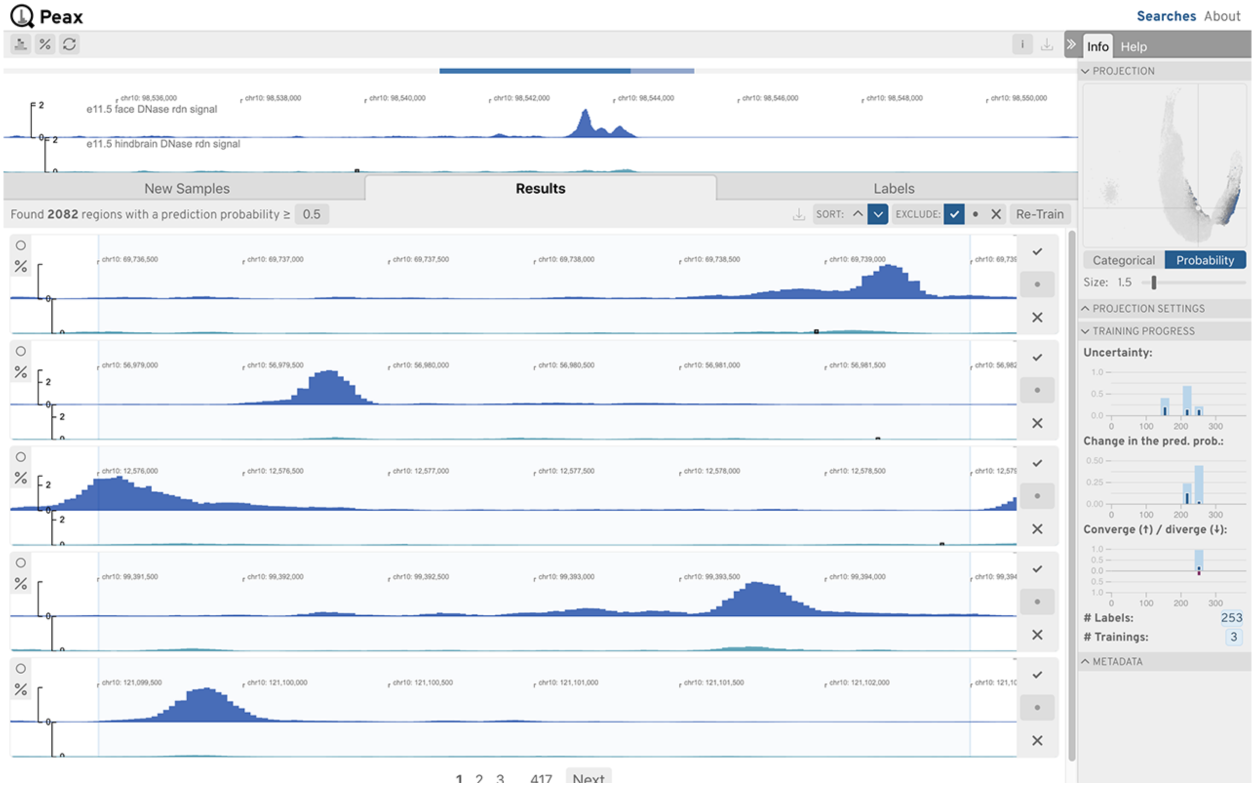


Figure S48: **Use Case 2: Differential Peak: Step 15.** Exploring those peaks in detail quickly reveals that the classifier works as expected.

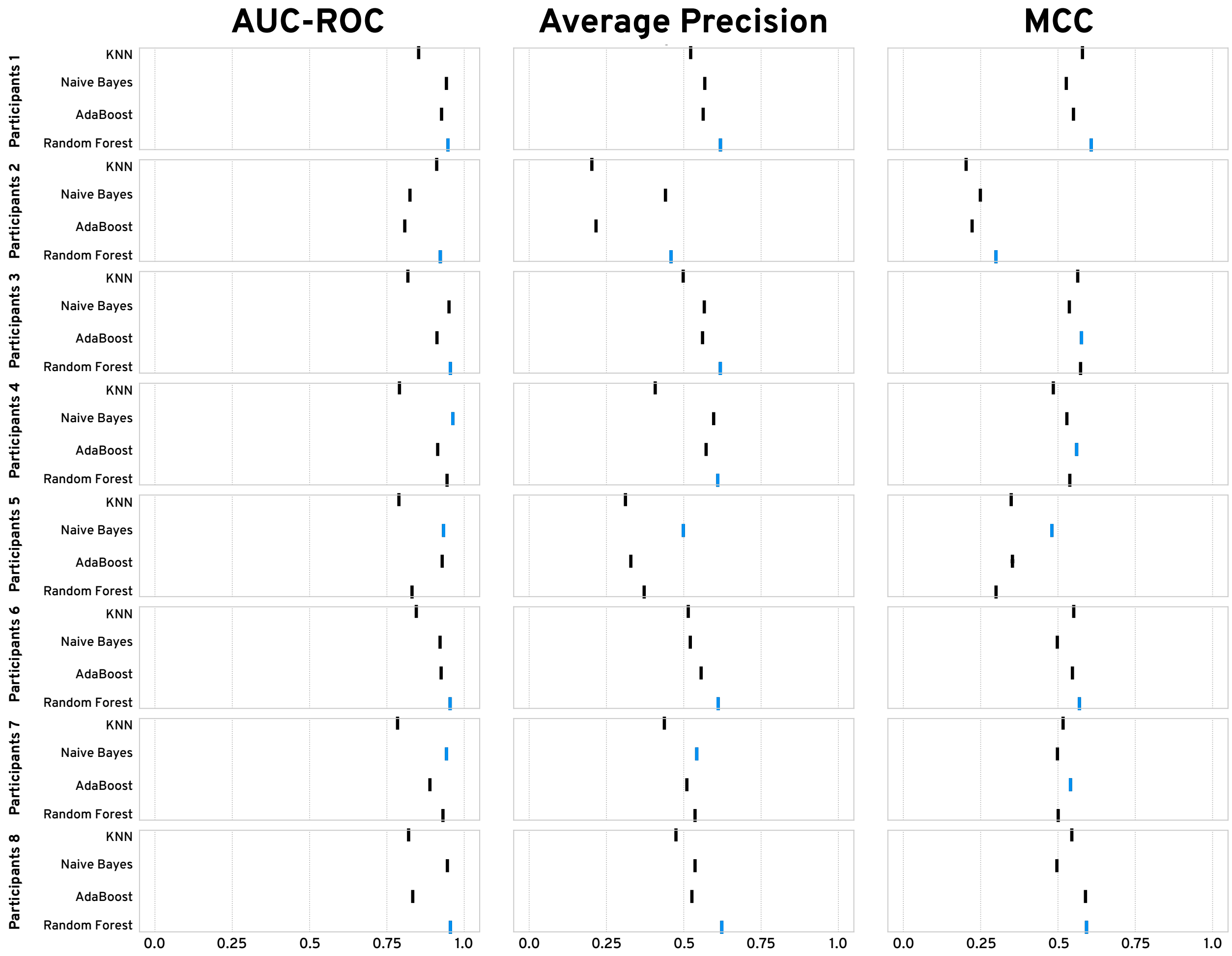


Figure S49: **Classifier Comparison:** Using our simulated experiments (Table S7) and the expert-defined labels from our in-person user study, we compared the performance of four commonly used binary classifiers: K-nearest neighbor search (KNN), Naive Bayes, AdaBoost, and Random Forest. For comparison, we choose the area under the receiver operating characteristic curve (AUC-ROC), average precision, and Matthews correlation coefficient (MCC) with a probability threshold of 0.5. Higher values indicate better performance. The best performing classifier is highlighted with a blue bar. Visually we find that on average, the Random Forest classifier outperforms all other classifiers except for the MCC score, where AdaBoost and Random Forest seem to do equally well. To confirm our intuition, we merged the rankings of the classifier across all metrics and participants with the Borda count method. The result confirms that the Random Forest classifier outperforms all other classifiers.

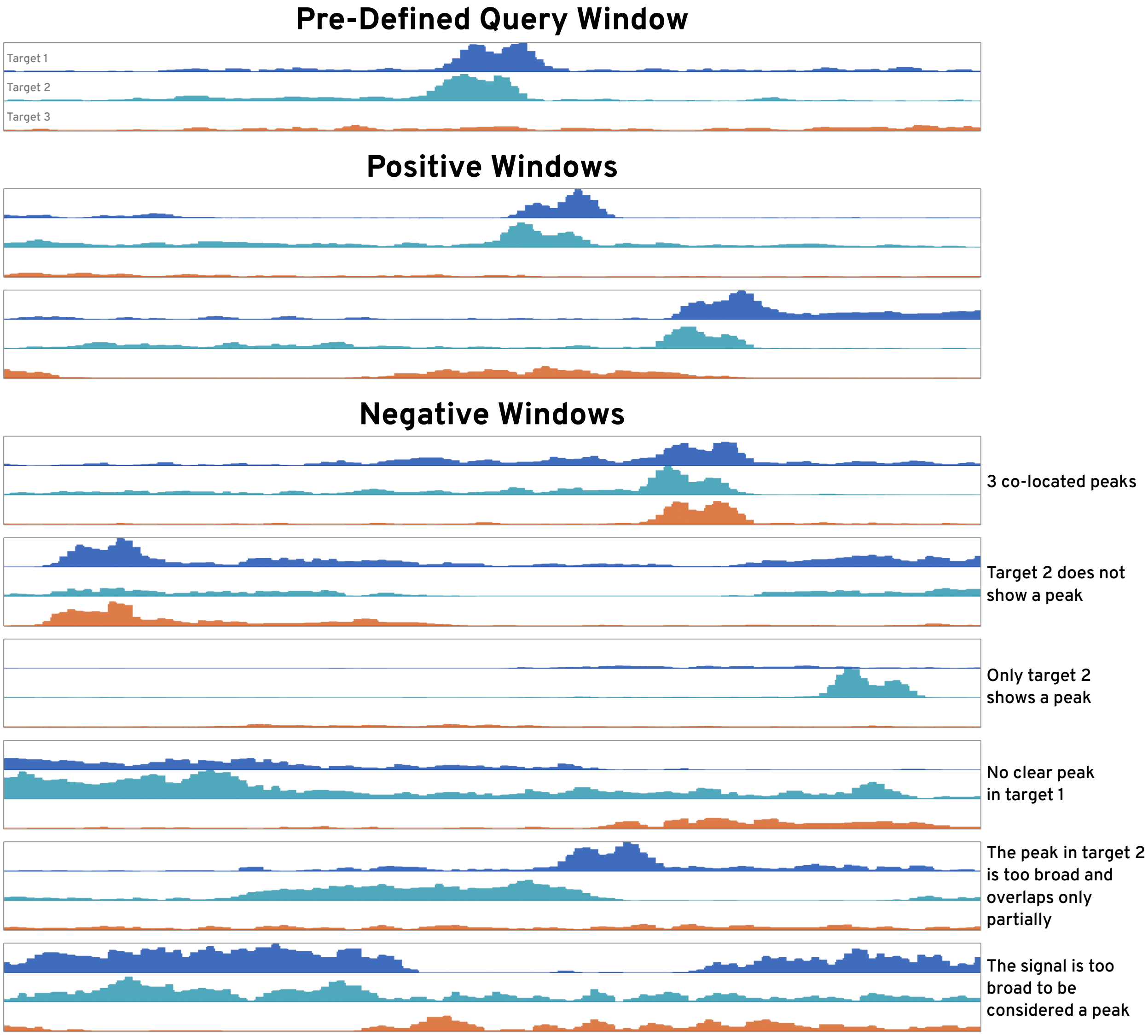


Figure S50: **In-Person User Study Task 1 Patterns:** We simulated three transcription factor binding ChIP-seq experiments (Table S7) and pre-defined a query window showing a pronounced co-located peak in target 1 and 2 and no peak in target 3. For context, we show a few positive and negative windows, i.e., windows that match the query window and windows that do not match the query window given the ground truth data.

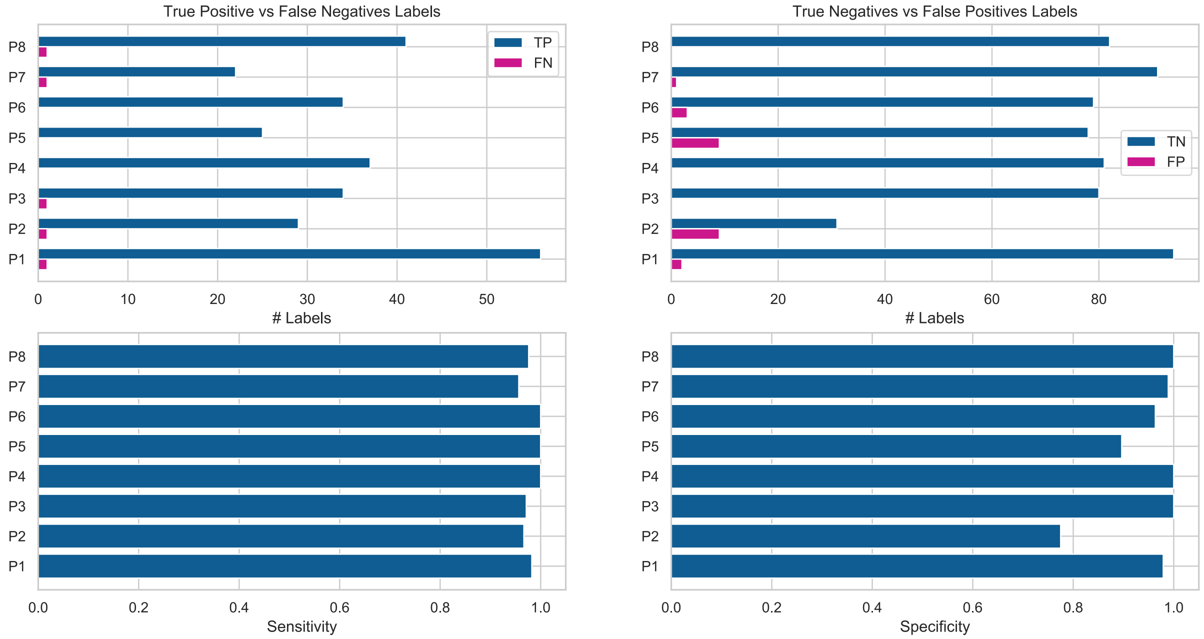


Figure S51: **In-Person User Study Task 1 Labels:** Using the simulated features as the ground truth, we can calculate the sensitivity and specificity of the experts' labels. Notice that P2 and P5 have higher false-positive labels compared to the other experts indicating that their imagined target pattern might have differed from ours. This is not necessarily unexpected as we did not ask them to find visually identical patterns but patterns that show similarity in terms of two peaks in the top two tracks and reduced signal in the bottom track.

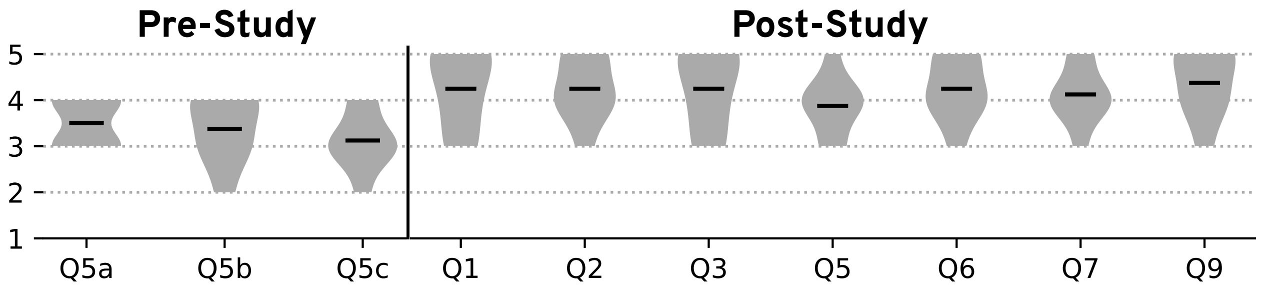


Figure S52: **Results of the Pre- and Post-Study Questionnaires.** Mean values are indicated by the black bars. For details see Table S9 and Table S10.

### Supplementary Tables

Table S1: **Training progress metrics.** To visually inform the user about the progress of the actively learned classifier we defined the following three progress metrics, where $(P)$ indicates the class prediction probabilities of all windows and **P3**, **P2**, and **P1** are the prediction probabilities of the current classifier, previous classifier, and the classifier before the previous classifier.

| |
|---|
| **Uncertainty:** |
| $$\sum_{i \in \mathbf{P}} \lvert \mathbf{P}_i - \text{round}(\mathbf{P}_i) \rvert / \lVert \mathbf{P} \rVert * 2$$ |
| **Prediction probability change:** |
| $$\sum_{i \in \mathbf{P1}} \lVert \mathbf{P1}_i - \mathbf{P2}_i \rVert / \lVert \mathbf{P1} \rVert$$ |
| **Convergence:** |
| $$\sum_{i \in \mathbf{P1}} \lfloor \lVert \text{sign}(\lfloor \mathbf{P2} \rceil_{\mathbf{d}} - \lfloor \mathbf{P1} \rceil_{\mathbf{d}}) + \text{sign}(\lfloor \mathbf{P3} \rceil_{\mathbf{d}} - \lfloor \mathbf{P2} \rceil_{\mathbf{d}}) \rVert / 2 \rfloor / \lVert \mathbf{P1} \rVert$$ |

Table S2: **Training Data.** ENCODE accession numbers and Roadmap Epigenomics experiment IDs for DNase-seq datasets and histone modification ChIP-seq datasets. The replicate number of DNase-seq experiments is listed in parentheses after the ENCODE accession numbers (ENCSR). For each dataset, we downloaded the fold change over background signal (Roadmap Epigenomics) or read-depth normalized signal (ENCODE) (bigWig), narrow peak annotations (bigBed), and broad peak annotations (bigBed)). We employed the peak annotations to balance the number of windows used for training that contain at least one peak annotation, that contain no peak annotation but are non-empty, and that are empty.

| **ENCODE accession number (ENCSR)** |
|---|
| 000EQB (2), 316UDN (1), 317SIH (1), 000EJO (1), 038XTK (1), 158VAT (1), 680SDS (1), 440FZS (1), 678PDD (1), 299INS (1), 000ENZ (1), 121ZSL (1), 796SJV (1), 035QHH (1), 515EWI (2), 000EQD (1), 271QSV (1), 426IEA (1), 000EPG (2), 000EIY (1), 595CSH (1), 000EQJ (1), 383BLX (1), 628IRM (1), 477RTP (1), 512CWR (1), 000EQI (1), 945RWN (1), 272RQX (1), 814KRX (1), 548MMD (1), 141VGA (1), 645GJD (1), 594NOE (1), 691MQJ (2), 000EPI (1), 381PXW (1), 468ZXN (1), 000EPE (2), 434OBM (1), 931UQB (1), 217RVH (1), 325LYJ (1), 004SUL (1), 035RVH (1), 217TAW (1), 184LMY (1), 940NLN (1), 000FFJ (1), 153LHP (1), 383SNM (1), 052AWE (1), 672EWY (2), 098PTC (2), 452DCM (1), 265TEK (1), 852TRT (1), 120LVW (1), 251UPG (1), 564JUY (1), 782XFY (1), 774RCO (1), 405TXU (1), 154ZNQ (1), 257BGZ (1), 148VUP (1), 593LTJ (1), 622TWS (1), 649KBB (1), 000ELO (1), 696TPW (1), 191EII (1), 019JDO (1), 000EML (1), 458LIB (2), 269SIA (2), 000EMR (1), 385AMY (1), 208DMX (1), 033STL (1), 683QJJ (1), 845CFB (1), 228VNQ (2), 517NHP (1), 337IRF (1), 000EPK (2), 554WUJ (1), 770DEN (1), 724CND (1), 911LTI (1), 857AEB (1), 959ZXU (1), 000EPD (1), 714DIF (1), 141NSQ (1), 083STA (1), 346IHH (1), 164WOF (1), 224FOA (2), 000EJQ (1), 621ENC (1), 228IKB (1), 954AJK (1), 206FSY (1), 275ICP (1), 552XJI (1), 445XYW (1), 166KPV (1), 935EVZ (1), 236SFP (1), 792ZXA (1), 426TPQ (1), 582IPV (2), 524OCB (1), 000EMV (1), 902XFY (1), 000EIS (1), 921NMD (1), 873ANE (1), 850YHJ (1) |
| **Roadmap Epigenomics experiment ID** |
| E003, E004, E005, E006, E007, E008, E011, E014, E015, E016, E017, E019, E020, E026, E038, E047, E049, E062, E063, E066, E067, E068, E069, E072, E073, E074, E075, E076, E087, E101, E102, E103, E108, E111, E114, E115, E116, E117, E118, E119, E120, E121, E122, E123, E124, E125, E126, E127, E128 |

Table S3: **Data Processing for DNase-seq and ChIP-seq data.** In the following we provide details about our applied data processing for DNase-seq and histone mark ChIP-seq data for the subsequent CAE training (Table S4).

| Step | Description |
|---|---|
| **Number of ChIP-seq dataset** | ChIP-seq 343 |
| **ChIP-seq targets** | H3K4me1, H3K4me3, H3K27ac, H3K9ac, H3K27me3, H3K9me3, and H3K36me3 |
| **Number of DNase-seq dataset** | 120 |
| **Window sizes** | 3 kb, 12 kb, 120 kb |
| **Step freq.** | 2, 3, 10 |
| **Step sizes** | 1.5 kb, 4 kb, 12 kb |
| **Resolution** | 25 bp, 100 bp, 1000 bp |
| **Filtering** | Since a large part of the epigenome contains little to no signal we employed the peak annotations to balance the number of windows used for training that contain at least one peak annotation, that contain no peak annotation but are non-empty, and that are empty. We chose to include two times more windows with peak annotations than non-empty windows without peak annotations, which were randomly sampled. Additionally, we added at most 5% empty windows to avoid biasing the autoencoder towards non-empty regions. Finally, since a large portion of non-empty regions are almost empty (very low signal) we additionally ensured that all windows with a total signal that is higher than the 50th percentile of the total signal of windows that contain peak annotations were included as well. |
| **Total number of ChIP-seq windows** | 3 kb = 11,515,378, 12 kb = 28,341,434, 120 kb = 54,415,566. |
| **Total number of DNase-seq windows** | 3 kb = 17,438,381, 12 kb = 41,026,177, 120 kb = 69,727,138. |

Table S4: **Model Training for DNase-seq and ChIP-seq data.** In the following we provide details about our CAE training on DNase-seq and histone mark ChIP-seq.

| Step | Description |
|---|---|
| **Loss** | log loss |
| **Optimizer** | ADADELTA [1] with the recommended default learning rate 1.0 and a learning rate decay of 0.001. |
| **Epochs** | 100 epochs with early stopping if the validation loss does not decrease for 10 epochs |
| **Batch size** | 256 |
| **Sample weighting** | We weight the loss of windows containing at least one peak annotation 8 times higher and additionally multiply the weighted loss by the total amount of signal such that the final loss of the $i$th window is given by. $$loss_i = \text{logloss}\, w_i * \ln S_i * \begin{cases} 8, & \text{if } w_i \text{ has at least one peak annotation} \\ 1, & \text{otherwise} \end{cases}$$ |
| **Data split** | We used 85% of the windows for training, 7.5% for validation, and the final 7.5% for testing. |
| **Environment** | NVIDIA Tesla V100 GPUs |

Table S5: **Hyperparameter Search.** We initially performed several iterations of hyperparameter search to determine which hyperparameters work well and which ones are irrelevant.

| Parameter | Tested Values | Insights |
|---|---|---|
| # conv. layers | 1, 2, 3, 4* | Until 3 layers the reconstruction quality increases. |
| Conv. sizes | 32, 64, 128, 256, 512, 1024 [32, 128], [64, 256], [32, 64, 128], [64, 128, 256], [64, 256, 1024], [32, 64, 128, 256], [64, 128, 256, 512], [128, 256, 512, 1024] | Larger conv. layers lead to lower loss but increase run time dramatically. |
| Conv. kernel sizes | 3, 5, 7, 9, 11 [3, 5], [5, 7], [3, 7], [3, 5, 7], [3, 7, 11], [3, 5, 7, 9], [3, 7, 11, 15] | Larger kernel sizes lead to lower loss but increase run time markedly |
| # dense layers | 0, 2*, 4, 5 | Dense units improve the reconstruction slightly until 2 layers. |
| Dense layer sizes | [512, 128], [1024, 256]*, [1024, 512, 128, 32], [1024, 512, 256, 64, 32] | The effect is similar to conv. layer sizes but less pronounced. |
| Embedding size | 6, 8, 10, 12, 15, 20 | 10, 12, 15, 20 have equal performance. Below 10 the reconstruction starts to decrease (very noticeable at 6) |
| Dropout | 0, 0.1, 0.2 | Dropout always increased the loss. Since there are no signs of overfitting we dropped dropout. |
| L1 regularization | 0, 0.00001, 0.001 | Gentle regularization (0.00001) does not hurt but doesn't help either |
| Loss functions | MSE, logcosh, log loss* | MSE and log loss led to lowest loss while log loss offered to more predictable training. MSE can get stuck after the first epoch. |
| Optimizer | sgd, nesterov, rmsprop, adagrad, adadelta*, adam, amsgrad, adamax, nadam | RMSprop and adadelta consistently lead to faster training than other optimizers. |
| Learning rate | 1.0*, 0.1, 0.01, 0.001 | The effectiveness of the learning rate depended on the optimizer. We found that the default recommendation works best. |
| Learning rate decay | 0, 0.001* | Gentle decay resulted in mildly lower loss. |
| Batch norm. | yes, no* | No effect |
| Sample weight by peaks | 1, 2, 4, 8* | Minimal effect but weighting seems to speed up learning. |
| Sample weight by signal | none, linear, log2, logn*, log10 | Minimal effect but weighting seems to Speed up learning. |

Table S6: **Similarity Measures.** We compared the latent representation computed by our autoencoder models against five other commonly use similarity measurements. For a fair comparison we did not include relevance feedback for feature-based methods and instead based extracted the five nearest neighbors in L2 space. Note that all methods operate on the piecewise aggregate approximation (PAA) representation of windows given their respective bin sizes as specified in Table S4

| | Methods | Type | Dim | Metric |
|---|---|---|---|---|
| **CAE** | Convolutional autoencoder (our method) | F | 10 | L2 |
| **UMAP** | Uniform Manifold Approximation and Projection [2] | F | 10 | L2 |
| **TSFRESH** | Time Series FeatuRe Extraction on basis of Scalable Hypothesis tests [3] | F | 10 | L2 |
| **ED** | Euclidean distance | D | 1 | *own* |
| **SAX** | Symbolic aggregate approximation [4] | D | 1 | *own* |
| **DTW** | Dynamic time warping [5] | D | 1 | *own* |
| **XCORR** | Zero-normalized cross correlation | D | 1 | *own* |

Table S7: **Data Simulation.** In the following we provide details about the data simulation for Task 1 of the in-person user study. Using ChIPsim [6] we simulated a 12 mega base pair-sized genome together with three transcription factor binding site experiments. The code for simulating the data is available at `https://github.com/Novartis/peax/tree/develop/experiments/simulation`

| Aspect | Description |
|---|---|
| **Genome** | 12 mega base pairs |
| **Read Length** | 50 base pairs |
| **Number of Read** | 1 million |
| **Transcription Factors** | 3 (TF1, TF2, and TF3) |
| **Features** | 7 (TF1 only, TF2 only, TF3 only, TF1+TF2, TF1+TF3, TF2+TF3, TF1+TF2+TF3) |
| **Feature Binding Probability** | 0.6% |
| **Read Alignment & Coverage** | We use BWA [7] to index the genome and align the reads. With Samtools [8] and Deeptools [9] we compute the read RPGC-normalized converage and generate bigWig [10] files. |

Table S8: **In-Person User Study Procedure.**

| Duration | Description |
|---|---|
| 5 mins | **Welcome**<br>Consent<br>Pre-study questionnaire |
| 10 mins | **Introduction**<br>Play 8-min introductory video<br>Briefly show where the prediction probability score and the navigational cues are displayed, explain what the light blue bounding box is standing for, and demonstrate how to activate the lasso tool. |
| 15–20 mins | **1. Task: Build a Classifier From Scratch**<br>*Background:* We loaded three transcription factor ChIP-seq datasets: target 1, 2, and 3 and already selected a search target for you. The search target pattern consists of co-occurring peaks in target 1 and target 2 and no peak in target 3! It typically requires 50–100 labels before the systems provides reasonable results.<br>*Instructions:* Train a classifier that can detect this pattern type by labeling regions in the datasets that contain patterns similar to the search target. It doesn't matter where the peaks in targets 1 and 2 are located in regards to the genomic window as long as they co-occur at the same location. And it also doesn't matter if other peaks are present elsewhere in the genomic window. Finally, there is no fixed end point. To the best of your knowledge, build a classifier that captures the above-described pattern as accurately as possible. |
| 15–20 mins | **2. Task: Improve a Pre-trained Classifier**<br>*Background:* We loaded two mouse e11.5 DNase-seq datasets from Encode together with their official peak annotations coming from a peak caller for two tissues: face and hindbrain. Given these peak annotations, we extracted differentially-accessible peaks in face to find tissue-specific regulatory elements. I.e., ideally you should see a peak in the face track but not in the hindbrain track.<br>*Instructions:* Evaluate the quality of the algorithmically-generated differentially-accessible peak annotations and improve the classifier to the best of your knowledge by validating and expanding the pre-assigned labels. |
| 5 mins | **Post-study questionnaire** |

Table S9: **Pre-study questionnaire of the in-person user study.** Each “X” stands for the answer of one of the participants. All questions were optional.

| | | | | | |
|---|---|---|---|---|---|
| **1. What is your gender?** | | | | | |
| Male: | | Female: | | Non-binary: | |
| XXXXX | | XXX | | | |
| **How old are you?** | | | | | |
| 18-24 | 25-34: | 35-44: | 45-54: | 55-64: | 65-74: |
| | XXXXXX | XX | | | |
| **3. What degree (if any) do you hold?** | | | | | |
| Bachelor: | | Master: | | PhD/MD: | |
| X | | | | XXXXXXX | |
| **4. What is your current profession?** | | | | | |
| Bachelor student: | Master student: | PhD/MD student: | Post-Doc: | Research associate: | Professor: |
| | | X | XXXXXX | XX | |
| **5. How much do you trust the results from...** | | | | | |
| | 1: Not at all | 2 | 3 | 4 | 5: Completely |
| A) Peak caller? | | | XXXX | XXXX | |
| B) Differential peak caller? | | X | XXX | XXXX | |
| C) Feature caller? | | X | XXXXX | XX | |

Table S10: **Closing questionnaire of the first user study.** Each "X" stands for the answer of one of the participants. Free text responses are paraphrased and grouped. All questions were optional.

| **1** | **2** | **3** | **4** | **5** |
|---|---|---|---|---|
| **1. How easy was it to understand the main idea behind Peax's method for visual pattern search?** | | | | |
| *very hard to understand* | | | | *very easy to understand* |
| | | XX | XX | XXXX |
| **2. How easy is it to learn how to use the user interface?** | | | | |
| *very hard to learn* | | | | *very easy to learn* |
| | | X | XXXX | XXX |
| **3. How easy is it to use the user interface for visual pattern search?** | | | | |
| *very hard to use* | | | | *very easy to use* |
| | | XX | XX | XXXX |
| **4. Are there any parts of the user interface that were not easy to understand, learn, or use?** | | | | |
| • Progress view is not clear (2x)<br>• More detailed instructions for training would be useful (2x)<br>• Manual interactions to label the data is not ideal<br>• The embedding view is not immediately clear<br>• Re-train button isn't immediately noticeable | | | | |
| **5. How much do you trust the results from Peax' classifier?** | | | | |
| *not at all* | | | | *completely* |
| | | XX | XXXXX | X |
| **6. How useful is visual pattern search for exploring/labeling epigenomic data?** | | | | |
| *not useful at all* | | | | *very useful* |
| | | X | XXXX | XXX |
| **7. How useful is the tool in its current form to you?** | | | | |
| *not useful at all* | | | | *very useful* |
| | | X | XXXXX | XX |
| **8. Which features (if any) are missing that would make this tool more useful?** | | | | |
| • Exporting results (2x)<br>• Integration with HiGlass [11]<br>• Show embedding view in a separate window<br>• Allow panning & zooming in the results panel<br>• Show the number of positive and negative labels<br>• Help with inconclusive patterns<br>• Provide case studies for initial guidance | | | | |
| **9. Imagine all missing features are implemented, how useful would do you think would this tool be to the research community?** | | | | |
| *not useful at all* | | | | *very useful* |

| | | | | |
|---|---|---|---|---|
| | | X | XXX | XXXX |

**10. Is this kind of visual pattern search currently possible with other tools/methods?**

- None (8x)
- The best one can do is manually browse in a genome browser (2x)

**11. If so, how does Peax compare against the other methods in terms of performance (A) and features (B)?**

*No responses. See question 10.*

**12. Which search strategy do you think is more useful?**

| | |
|---|---|
| 1. Starting with one target from scratch: | |
| 2. Starting with an existing set of annotations: | XX |
| 3. Both: | XXXX |
| 4. None/other: | XX |
| - The use case will depend on the context | |

**13. Which goal do you think is more common and/or important?**

| | |
|---|---|
| 1. Find the k-most similar genomic regions: | |
| 2. Extract all genomic regions that contain a similar pattern: | XXXXX |
| 3. Both: | XXX |
| 4. None/other: | |

**14. Do you want to tell us anything else?**

- It's great to that the classifier is refined interactively (4x)
- Very intuitive and usable after watching just the introduction video
- A biology-focused introductory video would be useful
- This kind of search tool would be very useful for chromatin interaction data, e.g., Hi-C (2x)
- Noticed that their own judgment changes over time. It would be great to test the "stability" of users over time. I.e., how much variation is there within a human's visual judgment?
- Remove low signal regions automatically
- It takes some rounds of labeling before the classifier finds useful things

Table S11: **In-Person User Study Task 1 Protocol.** Chronological overview of the participants' actions in task 1.

**Participant P1:**

• Confirms target pattern
• Clicks on *Getting Started* to load sample regions for labeling
• Labels regions as positive, negative, and inconclusive
• Trains their first classifier and keeps labeling
• At 100 samples (8min), P1 noted to do one more round of training before switching to the results
• Is undecided between the *Results* tab and *Compute Projection* button
• Switches to the *Results* tab and notes that they found 488 regions with a prediction probability threshold of 0.5 and 5 potential conflicts
• Investigates the first page with the top results and notes that these look pretty good
• Inverts the ordering of results to see regions with a prediction probability close to the threshold of 0.5
• Decides to first look at the conflicts
• Confirms that the labeling of the false positives conflicts is correct so they switch back to the *Results* tab and increase the prediction probability threshold to 0.6
• Looks at newly appeared false-negative conflicts and decides to re-train the classifier after confirming that the assigned labels are correct
• After the last training, only one potential conflict is left, which P1 ignores after looking at it
• Computes the projection for the embedding view
• Adjusts the dot size and switches to the prediction probability colormap
• Identifies a neighborhood of positive regions in the embedding view
• Lasso-selects the previously found neighborhood and assigns labels to the selected regions
• Re-trains the classifier
• Investigates the results with the highest prediction probability
• Inverts the ordering of results to see regions with a prediction probability close to the threshold of 0.6.
• Acknowledges that the results look good but sees a few false positive hits and increases the prediction probability to 0.65
• Notices that they could do a bit more refinement but states that the results look great

**Participant P2:**

• Clicks on *Getting Started* to load sample regions for labeling
• Labels regions as positive, negative, and inconclusive
• Is looking how to see results and finds the *Train & Load More* button to train their first classifier
• Briefly switches to the *Results* tab
• Switches to the Label tab and recognizes the regions as their input
• Switches back to the *Results* tab and recognizes the regions as the predictions
• Recognizes that they need to expand the labels as the predictions don't look good yet. P2 notices that most predictions are wrong and that their training label set is too small
• Labels results and re-trains the classifier for two rounds
• Notices that the results improve
• Computes the embedding view
• Is unsure about the embedding view but then switches to the prediction probability colormap and recognize a neighborhood with regions that potentially match the target pattern
• Selects regions in the previously identified neighborhood in the embedding view and confirms her hypotheses

**Participant P3:**

• Clicks on *Getting Started* to load sample regions for labeling
• Labels regions as positive, negative, and inconclusive
• Trains the first classifier

• Keeps labeling and training for another 3 round until they labeled 85 windows
• Switches to the *Results* tab
• Acknowledges that the top hits look good and immediately inverses the sort order to look at the prediction probability boundary
• Keeps labeling regions with a prediction probability close to 0.5
• Retrains the classifier
• Computes the embedding view
• Pans & zooms in the embedding view and lasso-selects several regions of close to investigate neighborhoods close to positively and negatively labeled regions

**Participant P4:**
• Goes to the *New Sample* tab to sample regions for labeling
• Labels regions as positive, negative, and inconclusive
• Trains the first classifier and continues labeling
• Asks about a value scale normalization and uses the window-specific normalization button to compare a window with low signal to other windows
• Continues labeling regions and training classifiers for another four rounds
• Switches to the *Results* tab and investigates the top hits
• Recognizes that the classifier learned to find the target pattern
• Added more positive labels to good results that aren't labeled yet
• Switches to potential false-positive conflicts but concludes that the assigned labels are correct
• Switches back to the results and increases the prediction probability threshold to 0.7 to avoid false-positive hits
• Investigates false negatives regions and fixes the labeling of one region that is tricky to label
• Switches back to the results and confirms that the effect of the prediction probability threshold by inverting the sort order of the hits to see regions with a prediction probability close to 0.7
• Reverses the sort order again and adds more labels to positive regions
• Re-trains the classifier and acknowledges that the most confident results are showing the target

**Participant P5:**
• Goes to the *Labels* tab but realizes that it's not the right tab
• Goes to the *New Sample* tab to sample regions for labeling
• Labels regions as positive, negative, and inconclusive
• Trains the first classifier
• Continues labeling regions and training classifiers for another five rounds
• Switches to the *Results* tab and looks at the top results
• Recognizes that some results look good but that there are a few false-positive hits
• Assigns labels to false-positive hits

**Participant P6:**
• Goes to *New Samples* to sample regions for labeling
• Labels regions as positive, negative, and inconclusive
• Trains the first classifier
• Continues labeling regions and training classifier
• After skipping regions with low and broad differential signal, P6 notices that the number of regions they perceive as inconclusive increases
• Notice that the uncertainty graph in the training process changes but that it doesn't decrease
• Concludes that they should label inconclusive low and broad differential signal as negatives instead of inconclusive because these windows don't show narrow peaks
• Continues sampling and labeling regions and training classifiers for another three rounds
• Computes the embedding view and loads the results
• Looks at one potential false-positive region but confirms the previously assigned label

• Goes back to the results and looks at the first two pages
• Acknowledges that the top results look good and goes to the last page to look at regions close to the prediction probability threshold
• Labels regions at the prediction probability boundary and re-trains the classifier

**Participant P7:**
• Clicks on *Getting Started* to load sample regions for labeling
• Labels regions as positive, negative, and inconclusive
• Trains the first classifier
• Continues labeling regions and training classifiers for three more rounds
• Notes that it would be useful to see how many positive and negative labels have been assigned so far
• Notices that the overall uncertainty is low after four rounds of training and switches to the *Results* tab
• Notices and inspects one potential false-negative conflict
• Fixes the positive label to inconclusive and re-trains the classifier
• Computes the embedding view
• While the embedding view is computed, P7 examines the first six pages of the results and assigns labels to what they think are true and false positive hits
• Re-trains the classifier and inspects the top results and the results close to the prediction probability threshold of 0.5
• Acknowledges that the results look "really good"
• Briefly inspects the embedding view to look at which neighborhoods contain positive and negative labels

**Participant P8:**
• Pans the query views
• Goes to *New Samples* to load sample regions for labeling
• Labels regions as positive, negative, and inconclusive
• Trains the first classifier
• Continues sampling and labeling regions and training classifiers
• Briefly checks the uncertainty in the progress view and notes to themselves that the uncertainty looks "okay"
• Continues sampling and labeling regions and training classifiers
• After training four classifiers, P8 notes that the uncertainty in the progr ess view goes down and switch to the *Results* tab
• Goes through the results pages and acknowledges that the results look good so far
• Improves the labels by assigning positive labels to true-positive regions and negative labels to false-positive regions
• Re-trains the classifier after going through all nine pages of results
• Spots a false-positive conflict but confirms that the assigned label is correct
• Computes the embedding view
• Switches to the prediction probability colormap and examines an uncertain region in a neighborhood of regions that are predicted to be negative

Table S12: **In-Person User Study Task 2 Protocol.** Chronological overview of the participants' actions in task 2.

**Participant P1:**

- Confirms regions that they perceive as true positive by assigning labels
- Acknowledges that the top results look great
- Notices the very high number of predicted hits and inverts the order to look at results close to the prediction probability threshold
- Acknowledges that the results close to the prediction probability threshold don't show a differentially-accessible peak
- Decides to first assign negative labels before they increase the prediction probability threshold
- After assigning labels and re-training, they notice that the number of found regions didn't go down dramatically, so they increase the prediction probability threshold to 0.65
- Labels more regions at the prediction probability threshold
- Increases the prediction probability threshold further to 0.8 and labels more regions at the prediction probability threshold
- After re-training, they switch to the conflicts and fix several incorrectly pre-loaded labels.
- After retraining, they acknowledge that the results look pretty good so far and they decide to compute the projection for the embedding view
- During the computation, they investigate the training process views
- Examines the projection view and zooms into a neighborhood with regions predicted to be positive results
- Lasso-selects several regions and recognizes them as true positive hits

**Participant P2:**

- Acknowledges that the classifier is pre-trained and starts expanding the labels of the results
- Acknowledges that the first couple of pages look all "pretty good" and can't find any negative hits on the first 15 pages
- Tries to label some regions in the middle of the results, by increasing the prediction probability threshold to 0.85 and inverting the sort order of the results
- Labels regions at the prediction probability boundary
- Re-trains the classifier
- Increase the prediction probability threshold to 0.9
- Continues labeling regions and acknowledges that the results look much better now
- Re-trains the classifier
- Computes the embedding view
- Inspects the embedding view but is unsure how to interpret the shape of the clusters
- Explores the embedding space by looking at regions (through mouse click-based selection) of potential negative hits
- Switched to the probability colormap and identifies and confirms the primary neighborhood with true positive matches

**Participant P3:**

- Looks at the results on the first seven pages
- Acknowledges that the results look good and realizes that they are predicted from the pre-loaded labels
- Switches back to page one of the results and assigns labels to regions showing the target pattern on the first two pages of the results
- Inverts the sort order to look at the prediction probability boundary
- Labels regions at the prediction probability boundary and re-trains the classifier
- Continues labeling at the prediction probability boundary and re-trains the classifier
- Computes the embedding view
- Explores the embedding view and lasso-selects regions in a separated cluster of dots

• Discovers that all regions of this cluster are empty (i.e., the value of the sequential data is zero across the entire region)
• Lasso-selects another neighborhood and discovers regions with low signal. P3 also recognizes that the classifier is uncertain about these regions
• Labels regions with low signal as negative and re-trains the classifier
• Adjusts the prediction probability threshold to 0.8

**Participant P4:**
• Stays in the *Results* tab as they notice that the classifier is trained already with the pre-loaded labels
• Investigates the first two pages of the top results and labels the predicted results
• Inverts the sort order to look at regions that are close to the prediction probability boundary
• Labels regions at the prediction probability boundary
• Looks at false negatives but confirms the correctness of the assigned labels
• Switches back to results and find more conflicts, which turn out to be recently assigned labels
• Re-trains the classifier
• Assigns labels to regions with a high prediction probability
• Noticed that the classifier changed and that it shifted the target pattern
• Assigns labels to the top results
• Increases the prediction probability threshold to 0.8
• Computes the embedding view
• While the embedding view is being computed, P4 notices the high divergence in the progress view
• Switches to the embedding view, identifies the neighborhood with currently labeled regions, zooms into it, and lasso-selects dots in this neighborhood
• Switches to the prediction probability colormap
• Identifies a neighborhood with regions that are predicted to match the search target
• Consecutively lasso-selects small regions and assigns more labels

**Participant P5:**
• Starts in the *Results* tab and assigns labels to regions that match the target pattern but are not yet labeled
• Briefly looks at the already labeled regions but switches back to the results
• Looks at two potential false-negative conflicts but confirms that the correctness of the assigned labels
• Switches back to the results and inverts the sort order to look at regions close to the prediction probability boundary
• Assigns labels to regions close to the prediction probability boundary
• Increases the prediction probability threshold to 0.8
• Re-trains the classifier

**Participant P6:**
• Starts in the *Results* tab and assigns labels to regions that match the target pattern but are not yet labeled
• After investigating the first three pages, P6 acknowledges that the top results look “very good” and switches to the last page to look at regions that the classifier is uncertain about
• Re-trains the classifier
• Computes the embedding view
• While the embedding view is being computed, P6 looks at the progress view and mentions that they are not sure about their definition
• Investigates the embedding view and zooms into a neighborhood that contains positively and negatively labeled regions
• Lasso-selects multiple regions in the previously-found neighborhood
• Expands and fixes labels of the selected regions and re-trains the classifier
• Adjusts the point size of the embedding view to identify a neighborhood with regions that are predicted to match the target

• Assigns labels to regions that they perceive as true positives
• Switches to new samples to look at regions Peax's classifier is unsure about as they do not perceive the region with the highest prediction probability matches the target pattern well
• Labels two pages of new samples and re-trains the classifier
• Switches to the *Results* tab
• Increases the prediction probability threshold to 0.9 and looks at regions close to the prediction probability thresholds close to the prediction probability threshold

• Labels regions at the prediction probability boundary and re-trains the classifier

---

**Participant P7:**
• Starts in the Results tab and assigns labels to regions that match the target pattern but are not yet labeled
• After looking through the first two pages, P7 investigates the potential conflicts highlighted by Peax but confirm the correctness of the assigned labels
• Switches back to the Results tab and continues assigning positive labels to top results
• After assessing the first six pages, P7 switches to the last page to assess regions close to the prediction probability boundary
• Labels regions on the last six pages and then re-trains the classifier
• Assigns negative labels to unlabeled false-positive regions of the top results
• After labeling false positives, P7 realizes that they forgot to assign positive labels to unlabeled true positive regions. They switch back the first page of the results and quickly assigns positive labels to regions that they perceive as true positives
• Re-trains the classifier
• Switches to regions close to the prediction probability threshold and labels several false-positive regions
• Switches back to the first page of the results and labels several true positive regions
• Re-trains the classifier
• Notices three potential false-negative conflicts and inspects them
• Fixes one label and confirms the correctness of the other two assigned labels
• Re-trains the classifier
• Increases the prediction probability threshold to 0.7
• Inspects regions with a prediction probability close to or equal to 0.7
• Assigns some more negative labels and re-trains the classifier
• Inspects the results and mention they are happy with them

---

**Participant P8:**
• Starts in the Results tab and assigns labels to regions that match the target pattern but are not yet labeled
• Continues assigning labels to the first eight pages of the top search results
• Computes the embedding view
• Switches to the prediction probability colormap
• Finds out that most of the chromosome is considered to be a match by the classifier
• Re-trains the classifier
• Increases the prediction probability threshold to 0.75
• Spots the warning for ten potential false-negative regions and investigates them
• Fixes some labels and confirms the correctness of other assigned labels
• Re-trains the classifier
• Fixes assigned labels for newly appearing conflicts
• Re-trains the classifier
• Notices that the number of potential hits went down 3-fold

• Inspects the updated prediction probability colormap of the embedding and notes that still a lot of regions are considered positive hits

Table S13: **Usability Improvements After the In-Person User Study.**

| |
|---|
| Allow sorting selection regions by prediction probability. |
| Show an indicator if the prediction probability is below the prediction probability threshold. |
| Add legends for the colormap to the embedding view. |
| Increase the contrast of the *Re-Train* and *Compute Projection* buttons. |
| Split *# Labels* into the number of positive and negative labels. |
| Add title to the x-axis of the training progress plots. |
| Add *question mark* icons next to the 3 training progress plots to provide an explanation for each of the three bar charts. |
